# Putative Role of Norrin in Neuroretinal Differentiation Revealed by bulk and scRNA Sequencing of Human Retinal Organoids

**DOI:** 10.1101/2024.11.15.623746

**Authors:** Kevin Maggi, David Atac, Jordi Maggi, Silke Feil, Samuel Koller, Wolfgang Berger

## Abstract

Pathogenic variants in the X-linked gene *NDP* (Norrie disease protein) have been associated with a variety of non-syndromic and syndromic human retinal diseases, including Norrie disease and familial exudative vitroretinopathy. The gene codes for Norrin, a secreted angiogenic molecule which binds to FZD4 and its co-receptors LRP5/6 and TSPAN12 and activates Wnt-signaling. Additionally, it also potentiates Wnt-signaling by binding to the LGR4 receptor. Norrin was also found to exert a neuroprotective function in the retina, specifically for retinal ganglion cells. Furthermore, it was suggested to be involved in neurodevelopmental processes such as early neuro-ectodermal specification and differentiation, as well as maintenance of cochlear hair cells. To better understand the putative role of Norrin in neuronal cells of the retina we generated *NDP* mutant and eGFP-expressing *NDP* reporter human induced pluripotent stem cells, which were differentiated to retinal organoids. Bulk RNA sequencing and fixed single-cell RNA sequencing revealed alterations in gene expression as well as cellular composition, with increased proportions of retinal progenitors as well as Müller glia cells in *NDP^KO^* retinal organoids. Differential expression of genes related to glutamate signaling, Wnt and MAPK signaling, as well as neurogenesis was detected. Furthermore, genes associated with functions in the extracellular matrix were also differentially expressed. The considerable decrease in retinal neurons found in our *NDP^KO^* organoids suggest that Norrin is also important for retinal neurogenesis, which may precede the vascular manifestations in *NDP*-associated diseases.

## Introduction

The Norrie Disease Pseudoglioma gene (NDP, OMIM: 300658) is located on the short arm of the human X-chromosome and was identified in 1992 by positional cloning [1]. Since its identification, hundreds of different mutations in NDP were reported as being associated with distinct retino-vascular conditions of variable severity, including X-linked familial exudative vitroretinopathy (FEVR, OMIM: 305390), Coat9s disease (OMIM: 300216), persistent hyperplastic primary vitreous (PHPV) and Norrie disease (ND, OMIM: 310600)[237]. In ND patients, the ocular phenotype is often accompanied by extraocular symptoms which include progressive deafness, intellectual disability (ID) as well as other neurological and non-neurological manifestations [8].

*NDP* codes for Norrin, a secreted protein with a cystein-knot motive that acts primarily as a WNT analogue, binding to a receptor complex that includes FZD4, LRP5/6 and TSPAN12 [9311], leading to cytosolic enrichment of ³-catenin and activation of TCF/LEF transcriptional programs [12,13]. Some studies indicate that Norrin could also operate through alternative mechanisms. Specifically, the protein was found to bind and activate LGR4, resulting in Wnt-signaling potentiation [14]. In addition, Norrin was described to act as a BMP2/4 antagonist, leading to inhibition of TGF-³ signaling through SMAD1/5/8 [15].

Initial reports on an *Ndp^KO^* mouse model described cytoarchitectural abnormalities in the neuroretina, including abnormal retinal layering and disorganization and loss of the retinal ganglion cells (RGC) layer [16]. *Ndp*^KO^ mice also revealed a progressive loss of sensory hair cells (HC) in the cochlea and of RGC in the retina in addition to the abnormal vasculature [17,18]. This loss was found to be only partially caused by the abnormal cochlear vascular development. Indeed, it was demonstrated to be a direct consequence of the lack of Norrin signaling in HC, as this was found to be essential for their development and maintenance [19]. Furthermore, involvement of Norrin in early neuroectodermal specification has also been proposed [15].

Most of the current knowledge on the functions of the protein and pathomechanisms involved in traits associated with Norrin derive from animal studies, while human studies are still limited, mainly due to the inaccessibility of the affected tissues. Therefore, species-specific information on the processes underlying the pathological changes observed in patients are lacking. Human induced pluripotent stem cells (hiPSC) offer unprecedented opportunities in this regard, as they can be used to generate three-dimensional cell-aggregates known as organoids, which recapitulate developmental and cytoarchitectural aspects of specific organs, and serve therefore as powerful *in-vitro* models to study human disease [20]. Over the last few years, several studies used retinal organoids to study different ophthalmic diseases, including retinitis pigmentosa and Leber congenital amaurosis [21,22], or to investigate pathogenic mechanisms associated with mutations of specific genes such as *ATOH7* [23].

In this study we used human retinal organoids, which are devoid of vascular components, to study the function of Norrin in neuroretinal development and differentiation. We describe, for the first time, the transcriptomic consequences resulting from depletion or strong reduction of *NDP* transcripts levels in hiPSC-derived retinal organoids. Combining bulk RNA sequencing (bulk RNA-seq) and fixed single-cell RNA sequencing (scRNA-seq) in human retinal organoids (Figure 1), we detected early transcriptomic events in neuroretinal development and differentiation, which may underlie the pathological processes associated with ND and other *NDP*-associated conditions. We identified alterations in the expression of genes involved in Wnt-, MAPK- and NOTCH-signaling, as well as several genes associated with functions in the extracellular matrix (ECM). In addition, our data suggest that Norrin may participate in the modulation of the glutamatergic signaling system, which was recently described to be an important prerequisite for retinal angiogenic processes and maturation of the blood-retinal barrier. Interestingly, scRNA-seq also reveals a significant alteration in cell composition in avascular neuroretinal organoids. Consequently, this study offers the first evidence that Norrin signaling may play a role in the development and differentiation of neuroretinal cells independent from its function in retinal angiogenesis.

**Figure 1:**
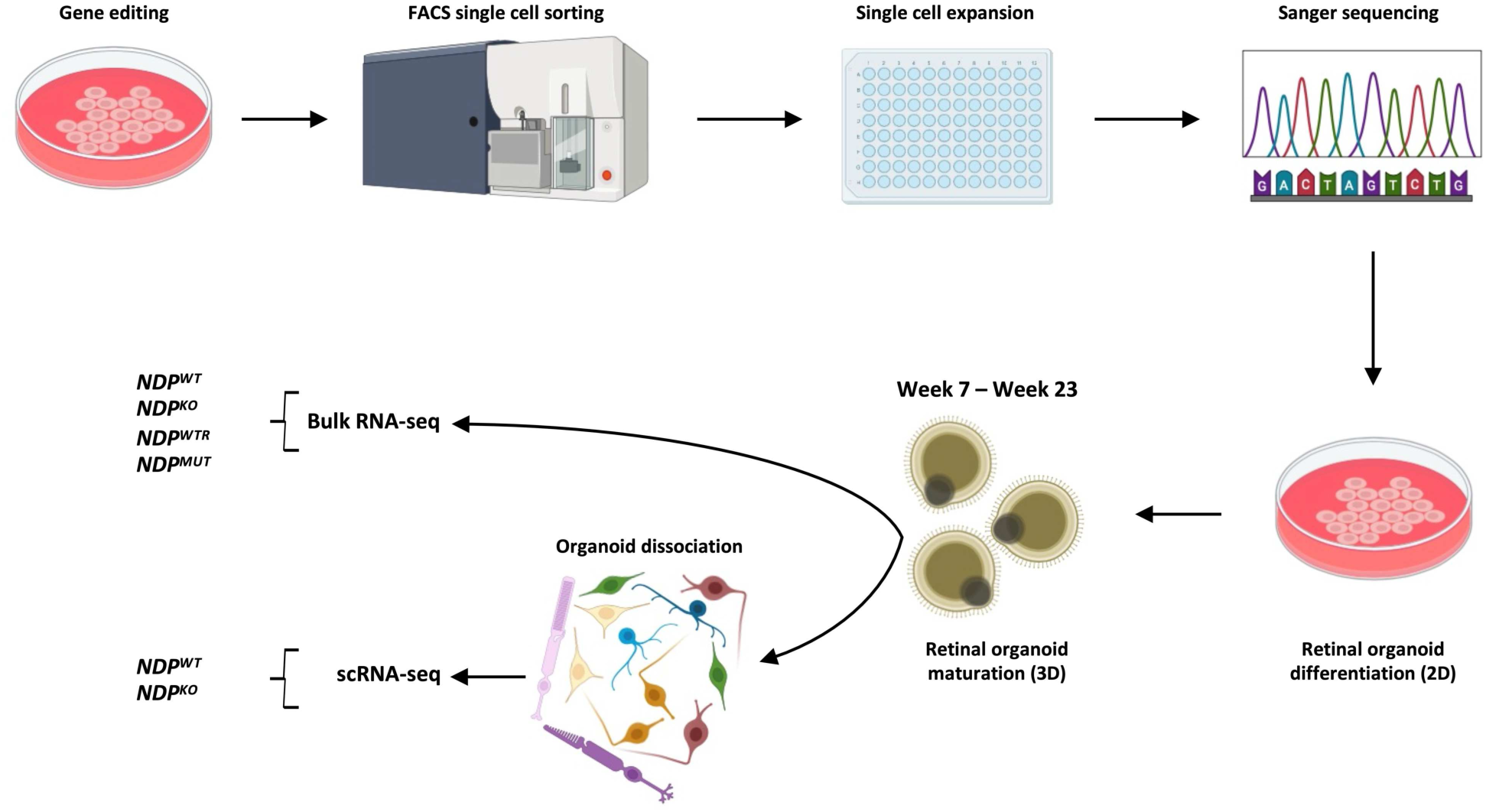
Schematic representation of the experimental setup. CRISPR/Cas9 was used to edit the *NDP* locus in hiPSC. Single cells were sorted from the edited population and expanded as clones. After genotyping by Sanger sequencing, selected clones were differentiated to retinal organoids. During the maturation process, samples were collected at different timepoints for downstream experiments. For the scRNA-seq experiments, 10 organoids were pooled and dissociated to single cell suspensions. Abbreviations: bulk RNA-seq, bulk RNA sequencing; FACS, Fluorescence-Activated Cell Sorting; KO, knockout; MUT, mutant; hiPSC, human induced pluripotent stem cells; scRNA-seq, single-cell RNA sequencing; WT, wildtype; WTR, WT-reporter.

## Materials and methods

### Human iPSC (hiPSC) culture

The fibroblast-derived female hiPSC line UCSD167i-99-1 was acquired from the WiCell Research Institute (WiCell, Madison, WI, USA). The cells were cultured in two-dimensional colonies in E8 Flex Medium (Thermo Fisher Scientific, Weltham, MA, USA) on truncated recombinant human vitronectin (rhVTN-N; Thermo Fisher Scientific, Weltham, MA, USA) at 37°C and 5.0% [CO_2_]. Medium was changed every other day and double medium volumes were used for over-weekend culture, according to the manufacturer9s instructions. Cells were split 1:4-1:12 using Versene Solution (Thermo Fisher Scientific, Weltham, MA, USA) 2-3 times per week. Cells were cryopreserved in Bambankers Standard freezing medium (Nippon Genetics Europe GmbH, Düren, Germany) according to the manufacturer9s instructions.

### CRISPR/Cas9 based targeted genome editing

#### Plasmid design

single guide RNAs (sgRNAs) ûanking exon 2 (sg1 and sg2) and the protein-coding sequence (CDS) of exon 3 (sg3 and sg4) were designed, which would lead to deletions of approximately 460 bp (exon 2) and 370 bp (exon 3 CDS), respectively (Figure S01A,B). The sequences of the guides and the top 10 oû-target regions for each guide are summarized in Table S01A-D. The sgRNAs were synthesized as single-stranded oligonucleotides (Microsynth AG, Balgach, Switzerland), dimerized and subsequently cloned into the Cas9 expressing EF1³-PX459 vector, a modified version of the pSpCas9(BB)2A-Puro (PX459) V2.0 plasmid (Addgene, Watertown, MA, USA), in which the cytomegalovirus (CMV) enhancer and the chicken ³-actin promoter have been replaced by the elongation factor 1 alpha (EF1³) promoter [23,24]. Separately, plasmids harboring repair templates for homology-directed repair (HDR) for the generation of *NDP*-eGFP reporters (*NDP^WTR^* [*NDP*-V5-P2A-EGFP], Data 01; *NDP^MUT^* [*NDP*^—ex2-CDS^- eGPF-P2A], Data 02) were designed. These comprised an *NDP*-P2A-eGFP expression cassette ûanked by 600-900 bp homology arms, which included protospacer adjacent motif (PAM)-site mutations and sites for plasmid linearization (Figure S01B). The repair templates were synthesized and cloned into a pUC57-mini vector (GenScript Biotech, Piscataway, NJ, USA). During CRISPR/Cas9 editing experiments, the BCL2L1_pLX307 plasmid (Addgene, Watertown, MA, USA) was transiently co-expressed to increase the yield of surviving edited cells [25].

#### Gene editing

hiPSC were dissociated using Versene Solution (Thermo Fisher Scientific, Weltham, MA, USA), gently resuspended to a single cell suspension in StemFlex# Medium (Thermo Fisher Scientific, Weltham, MA, USA) and seeded at 0.3M cells/well on rhVTN-N coated 12-well plates. Reverse-transfection was performed using a total of 1 µg of DNA, in proportions EF1³- PX459:pUC57-mini:BCL2L1_pLX307 2:2:1, and the Lipofectamine Stem# Transfection Reagent (Thermo Fisher Scientific, Weltham, MA, USA) following the manufacturer9s instructions. After seeding cells, transfection medium was supplemented with RevitaCell# Supplement (Thermo Fisher Scientific, Waltham, MA, USA) for 24 hours and, subsequently, antibiotic selection was performed for 3 days using 1 µg/ml puromycin containing StemFlex# Medium. After transfected cells were allowed to recover, these were dissociated using 0.75X TrypLE# Express Enzyme (Thermo Fisher Scientific, Weltham, MA, USA), and single cells were sorted into Corning Matrigel hESC-Qualified (Corning, New York, NY, USA) coated 96-well plates, filled with StemFlex# Medium supplemented with 1X RevitaCell#, using a FACSAria III 4L (BD Biosciences, Franklin Lakes, NJ, USA). After 3 days, medium was replaced with fresh StemFlex# and cells were allowed to expand for additional 7-10 days.

#### Colony PCR screening

Primers flanking the regions of interest were designed (exon 2: NDP_e2_59_fw and NDP_e2_39_rev; exon 3: NDP_e3_59_fw, NDP_e3_39_rev; Table S2A). Genomic DNA was directly extracted from pelleted cells using QuickExtract# DNA Extraction Solution (LGC Biosearch Teochnologies, Hoddeson, UK), following the manufacturer9s instructions. For each PCR reaction 2.5 ¿l of the extracted gDNA were used. PCRs were set up in volumes of 25 µl, containing 0.5 U Phusion HF Polymerase (New England Biolabs, Ipswich, MA, USA), 0.2 mM dNTPs (Solis BioDyne, Tartu, Estonia), 1X Phusion HF GC Buffer (New England Biolabs, Ipswich, MA, USA) and 0.5 ¿M primers and run with the following conditions: 1 cycle of 98°C for 30 sec; 35 cycles of [98°C for 10 sec, 70°C for 30 sec and 72°C for 2 min]; 1 cycle of 72°C for 10 min; hold at 10°C. After PCR termination, products were visualized on 1% agarose gels and edited clones were selected for further screening by sequencing. PCRs were also performed to screen for potential oû-targets effects. Primers were designed to screen for the four top oû-target sites of each sgRNA used for CRISPR/Cas9 gene editing (Table S2B).

#### Sanger sequencing

For the sequencing of the selected clones, BigDye# Terminator v1.1 Cycle Sequencing Kit (Thermo Fisher Scientific, Weltham, MA, USA) was used following the manufacturer9s instructions. Cycle sequencing reaction products were then purifieded using Sephadex# G-50 (Sigma-Aldrich, St. Louis, MO, USA) and sequenced on a SeqStudio# Genetic Analyzer (Thermo Fisher Scientific, Weltham, MA, USA).

### Retinal organoids differentiation

Differentiation of hiPSC to retinal organoids was performed using the 2D/3D differentiation protocol described previously by Gonzalez-Cordero et al and Cuevas et al [26,27]. Briefly, after reaching 60-70% hiPSC culture confluency, medium was changed to Essential 6 Medium (Thermo Fisher Scientific, Weltham, MA, USA) for 2 days, after which medium was switched to neural induction medium (NIM). At day 21 (D21) of differentiation, medium was switched to retinal differentiation medium (RDM) and between D28 and D35, organoids were manually dissected by 19G needles and transferred to suspension culture. From D35, medium was changed to fetal bovine serum- (FBS, Thermo Fisher Scientific, Waltham, MA, USA) and taurine-containing (Sigma Aldrich, St. Louis, MO, USA) retinal differentiation medium (RDM1), which was replaced at D63 with RMD1 containing retinoic acid (RA, Sigma Aldrich, St. Louis, MO, USA) (RDM2) and at D77, medium was then replaced with retinal differentiation medium 3 (RDM3). All retinal differentiation media were supplemented with 1X antibiotic-antimycotic (Thermo Fisher Scientific, Weltham, MA, USA).

### Bulk RNA-seq of human retinal organoids

#### Sequencing

RNA was extracted from 8-10 pooled organoids per sample using the NucleoSpin RNA kit (Macherey-Nagel, Düren, Germany) following the manufacturer9s instructions. Extraction was performed at two time points, week 7 and week 23 of differentiation. For week 7, RNA from six separate differentiations per condition was sequenced, whereas four differentiations per condition were used for week 23. Sequencing libraries were prepared with the Truseq total RNA (Illumina, San Diego, CA, USA) protocol with ribosomal depletion and sequenced as 100 bp single-end reads on a Novaseq 6000 sequencer (Illumina, San Diego, CA, USA).

#### Data pre-processing and mapping

raw data was pre-processed and assessed on the Galaxy Europe platform (http://usegalaxy.eu<u>, accessed 15.10.2023</u>). Reads quality assessment was performed using FASTQC v0.12.1 , whereas cutadapt v4.2 with the following settings was used for trimming [28]: --adapter sequence AGATCGGAAGAGCACACGTCTGAACTCCAGTCA \ --minimum- length 30 \ --quality-cutoff 20,20 \--trim-n. The trimmed reads were pseudoaligned with salmon v1.10.0 quant function [29], with the following settings: --ûdMean 150 \--ûdSD 70 \ -- validateMappings. As index, concatenated RefSeq GRCh38p14 DNA and RNA assemblies (gentrome) were used, whereas the corresponding RefSeq GRCh38p14 DNA assembly was used as decoy.

#### Differential expression analysis

Analysis of gene expression was performed in Rstudio v2022.12.0+353, running R v4.2.2. Filtering of genes with low expression was performed with edgeR::filterByExpression v3-40-2, while DESeq2 v1.38.3. was used to analyse differential expression in each condition [30]. Fold change correction was performed with DESeq2::lfcShrink using ashr v2.2-54. Benjamini-Hochberg FDR < 0.05 was used as thresholds to determine differentially expressed genes.

#### Over-representation analysis

Over-representation analysis (ORA) of gene ontology (GO) terms as well as of pathway enrichment, was performed using WEB-based Gene SeT Analysis Toolkit (WebGestalt; https://2024.webgestalt.org, accessed 10.04.2024), using the following parameters; functional database = GO biological process (GO:BP), GO Cellular Compartment (GO:CC), GO Molecular Function (GO:MF), pathway KEGG, pathway Reactome, pathway WikiPathways; minNum = 30; maxNum = 500, referenceSet = <genome_protein-coding=, multiple test adjustment = Benjamini-Hochberg FDR < 0.05. Results were then reduced using REVIGO (http://revigo.irb.hr/, accessed 13.04.2024), where needed.

### Fixed single-cell RNA sequencing (scRNA-seq) of human retinal organoids

#### Sample preparation

The paraformaldehyde (PFA)-fixed and hybridization-based scRNA-seq method following the Chromium Fixed RNA Profiling Technique (10X Genomics, Pleasanton, CA, USA) was selected to avoid composition biases due to potential loss of more sensitive retinal neuronal cells. Dissociated samples were prepared using the Chromium Next GEM Single Cell Fixed RNA Sample Preparation Kit (10X Genomics, Pleasanton, CA, USA), following the manufacturer9s instructions. The dissociation of the organoids to single-cell suspension was performed using the Papain Dissociation System (Worthington Biochemical Corporation, Lakewood, NJ, USA), utilizing approximately 1:10 of the reagents volumes recommended in the kit9s manual. Briefly, 10 organoids per sample were incubated in Papain/DNase mix on a thermomixer at 37°C for 35 min with shaking at 700 RPM. Subsequently, DNase and ovomucoid solution were added and organoids were gently triturated using a p200 micropipette with a wide-bore pipette tip, resulting in at least 1M cells with a viability of >85%. Cells were then fixed overnight with 4% PFA and supplemented with the fixative reagent. After washing the fixed cells twice, excessive debris was removed performing a discontinuous density gradient step. The cells were then resuspended in quenching buffer with the addition of the enhancer reagent and glycerol (10% final concentration) before cryostorage.

#### Sequencing and data analysis

Sequencing of the fixed single cells was performed on a Novaseq 6000 (Illumina, San Diego, CA, USA) as paired-end reads. Read mapping as well as counts quantification were performed with Cell Ranger multi v7.2.0, using the GENCODE GRCh38p13 genome assembly as reference and targeting 189075 genes. Data processing was then performed on the single-cell data analysis platform Trailmaker® (Parse Biosciences, Seattle, WA, USA; https://app.trailmaker.parsebiosciences.com, accessed 15.07.2024). Count matrices were uploaded, and data were filtered with the following parameters: barcode FDR <0.01 / minimum number of Unique Molecular Identifiers (UMI) per cell > 2000 / mitochondrial fraction <10% / scDblFinder probability threshold = 0.4. Integration of samples for batch effect correction was then performed by Harmony [31], using log normalization on 2000 highly variable genes and the number of principal components was set to explain >90% of the datasets variation. Dimensionality reduction for data representation was performed using the UMAP method and clustering was performed with the Luvain algorithm [32,33].

### Quantitative reverse-transcription PCR

#### Total RNA extraction and cDNA synthesis

8-10 organoids per extraction were pooled and RNA was extracted using the NucleoSpin RNA kit (Macherey-Nagel, Düren, Germany) following the manufacturer9s instructions and cDNA synthesis was performed using Superscript# III First- Strand Synthesis SuperMix (Thermo Fisher Scientific, Weltham, MA, USA) with oligo(dT)_20_ primers. The synthesized cDNA was then diluted to a concentration equivalent to 7.5 ng/µl of the input RNA.

#### Quantitative reverse transcription PCR

RT-qPCR was performed with SYBR# Select Master Mix (Thermo Fisher Scientific, Weltham, MA, USA) in 20 µl total volume. The reaction mix contained 0.4 µM RT-qPCR primers (see Table S2C) and 1 µl cDNA. The thermocycling reaction was set up as follows: 1 cycle of 50°C for 2 min; 1 cycle of 95°C for 2 min; 45 cycles of 98°C for 15 sec and 60°C for 1 min; and was performed using a LighCycler® 480 Instrument II (F. Hoffman-La Roche AG, Basel, Switzerland). The amplified products were subsequently confirmed by agarose gel electrophoresis.

#### Data analysis and visualization

Ct values were determined by 2nd derivative maximum analysis in the LightCycler 480 Software v 1.5.1.62 SP3 (F. Hoffmann-La Roche AG, Basel, Switzerland). The averaged Ct values of technical triplicates were normalized using the mean GAPDH expression and calibration using normalized expression values in hiPSC was performed, following the 2^2——Ct^ method [34]. Statistical analysis as well as data visualization and representation were performed using GraphPad PRISM v6.07 (GraphPad Software, San Diego, CA, USA).

## Results

### CRISPR/Cas9-based editing of the *NDP* locus in hiPSC

A dual sgRNAs (sg1 and sg2) approach was used to excise the entire exon 2 sequence by non-homologous end joining (NHEJ) to mimic pathogenic variants detected in some ND patients and the extensively studied *Ndp^KO^* mouse model [16]. In the process, unedited, isogenic wildtype hiPSC lines (*NDP^WT^*) were isolated, that were used as controls. Additionally, *NDP* reporter iPSC lines (*NDP^MUT^* and *NDP^WTR^*) were generated by targeting exons 2 and 3, respectively. For the *NDP^MUT^* reporter line, the same combination of sg1 and sg2 was used. The excised sequence was replaced by a plasmid-delivered eGFP-P2A sequence by HDR (Figure S01A). For the *NDP^WTR^*, the exon 3 coding sequence (e3CDS) was replaced by an NDP*^e3CDS^*-P2A-eGFP sequence using a different combination of sgRNAs (sg3 and sg4) (Figure S01B). Inspection of target sites, as well as of putative off-target (OT) sites of the sgRNAs confirmed the correct modifications in the selected clones and the absence of unwanted OT modifications (Figure S02-04). A schematic of the *NDP* locus in the cell lines used in this study is depicted in Figure 2.

**Figure 2:**
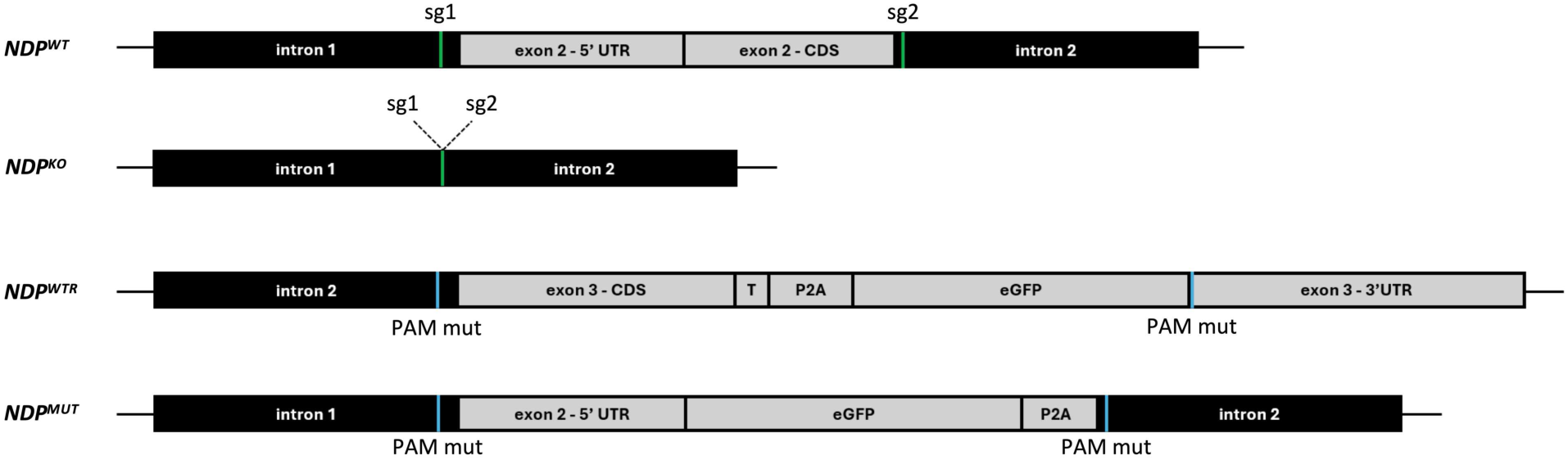
Schematic representation of the *NDP* locus of the hiPSC lines after editing, based on Sanger sequencing results. These included KO and WT isogenic clones (*NDP^KO^* and *NDP^WT^*) and WT and KO reporter lines expressing the green fluorescent protein under control of the endogeneous *NDP* promoter (*NDP^WT-GFP^* and *NDP^MUT^*). Abbreviations: CDS, coding sequence; eGFP, enhanced GFP; hiPSC, human induced pluripotent stem cells; KO: knockout; MUT, mutant; P2A, porcine teschovirus-1 2A self-cleaving peptide; PAM, protospacer-adjacent motif; PAM mut, PAM mutation; sg, small guide RNA; T, V5 Tag; UTR untranslated region; WT, wildtype; WTR, WT reporter.

### *NDP^KO^* organoids display comparable macroscopic morphology

Retinal organoids were generated following the protocol described in Gonzalez-Cordero et al and Cuevas et al [26,27]. The organoids were inspected at different stages of differentiation and were characterized by their distinct neuroretinal rim region, comprising radially organized cells (Figure S05A,B). No obvious macroscopical differences in morphology were detectable between WT and KO/MUT organoids, neither during the early (week 7) nor in the late (week 23) stage of differentiation (Figure S05A,B). *NDP* expression was detected in the *NDP^WT^* organoids at both timepoints, with increased levels in week 23 organoids. With the combination of primers used, no *NDP* transcripts were detected in KO organoids at any of the timepoints (Figure S05B,D). In the MUT organoids, qPCR revealed a strong decrease in *NDP* transcripts. In week 7 organoids we detected an approximately 98% reduction and in week 23 approximately 80 % reduction (Figure S05C). The strong reduction of WT *NDP* transcripts observed in the *NDP^MUT^* organoids was confirmed by bulkRNA-seq data. According to these observations, *NDP* expression begins early during retinal differentiation, at low levels, and increases with time. Additionally, *NDP^KO^* organoids do not seem to display obvious morphological aberrations in comparison with their WT counterparts in any timepoint analysed.

### *NDP* expression in human retinal organoids mainly localizes to retinal progenitor cells, a subset of RGCs and Müller glia cells

Early reports investigating the expression patterns of *Ndp* in murine retinae used RNA *in situ* hybridization, and detected signals in the inner nuclear layer (INL) and retinal ganglion cell layer [16]. Later studies used an *Ndp* alkaline-phosphatase (AP) reporter mouse strain and detected AP signals spanning the whole thickness of the retina, implying that Müller glia cells (MGC) are the main source of Norrin in the murine retina [13]. To better understand the expression patterns of the *NDP* gene in the human retina, we performed PFA fixed scRNA-seq on week 7 and week 23 *NDP^WT^ and NDP^KO^* dissociated organoids (3 replicates per genotype) using the Chromium Fixed RNA Profiling method (10X Genomics, Pleasanton, CA, USA). After applying quality filters, different numbers of cells were recovered from the sequenced samples (week 7: 10279-12091; week 23: 6365-10260, Figure S06). For processing and analysis, we used the Trailmaker platform (https://app.trailmaker.parsebiosciences.com, accessed 15.07.2024 [Parse Biosciences, Seattle, WA, USA]), using the Harmony algorithm for sample integration, with 26 and 27 principal components, for week7 and week 23 samples respectively, explaining >90% of the expression variation.

In the early timepoint (week 7), seven different main cell clusters were identified based on Luvain clustering (Figure 3A). Upon inspection of their transcriptional profiles, these were labeled as proliferating retinal progenitor cells (pRPC), characterized by markers such as *MKI67* and *TOP2A*, retinal progenitor cells (RPC) expressing marker genes such as *SFRP2* and *FZD5*, transient retinal progenitor cells (tRPC) expressing genes such as *ATOH7* or *GADD45A*, photoreceptor precursors (PRp) expressing for example *CRX* and *OTX2*, horizontal and amacrine cell precursors (H&A) characterized by the expression of e.g. *TFAP2A* and *ONECUT2* and early and late retinal ganglion cells (eRGC and lRGC), expressing higher levels of *POU4F2* and *SLIT1* as well as *STMN2* and *GAP43,* respectively (Figure S07)[23,26,35337]. *NDP* expression was found to be isolated to a small group of cells (2.82% in WT), which were localized mostly in the RPC (55.5%) and pRPC (16%) clusters with sub populations of tRPC (6.9%), eRGC (13%) and lRGC (6.24%) also showing some expression of the *NDP* gene (Figure 3B, Data 03).

**Figure 3:**
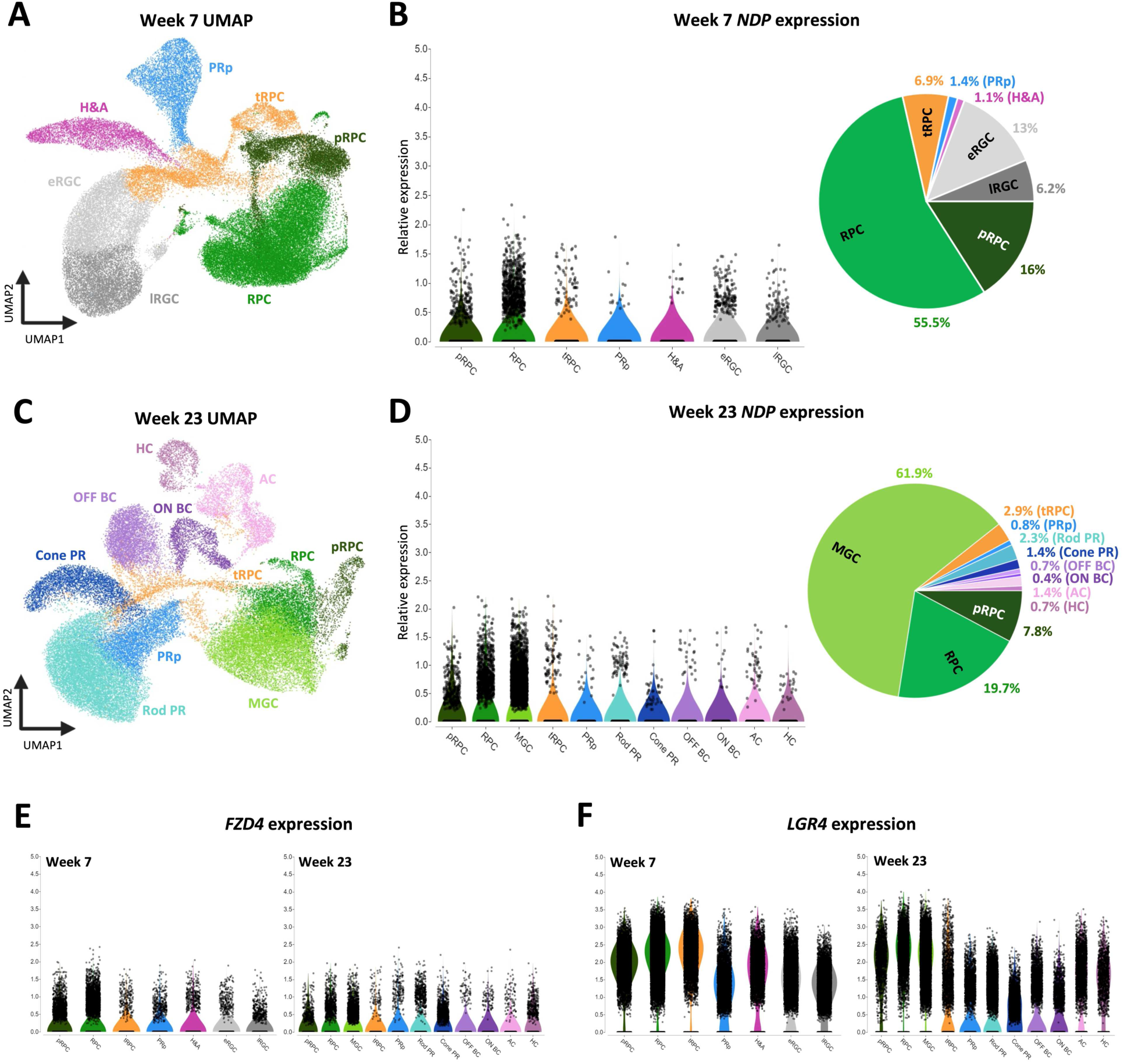
Expression of *NDP* and its receptors *FZD4* and *LGR4* in scRNA-seq clusters. (**A**) UMAP representation of retinal clusters from week 7 *NDP^WT^* and *NDP^KO^* organoids. The following clusters were identified: proliferative retinal progenitor cells (pRPC), retinal progenitor cells (RPC), transient retinal progenitor cells (tRPC), photoreceptor precursors (PRp), horizontal and amacrine cells (H&A), early retinal ganglion cells (eRGC) and late retinal ganglion cells (lRGC). (**B**) Violin plot and pie chart representing *NDP* expression in scRNA-seq clusters in week 7 organoids. Each cell expressing *NDP* is marked by a black dot. The percentages reported in the pie chart represent the proportion of cells within the specific cluster where *NDP* expression was detected. (**C**) UMAP representation of retinal clusters from week 23 *NDP^WT^* and *NDP^KO^* organoids. The following clusters were identified: proliferative retinal progenitor cells (pRPC), retinal progenitor cells (RPC), Müller glia cells (MGC), transient retinal progenitor cells (tRPC), photoreceptor precursors (PRp), rod photoreceptors (Rod PR), cone photoreceptors (Cone PR), OFF-bipolar cells (OFF BC), ON-bipolar cells (ON BC), horizontal cells (HC) and amacrine cells (AC). (**D**) Violin plot and pie chart representing *NDP* expression in scRNA-seq clusters in week 23 organoids. Each cell expressing *NDP* is marked by a black dot. The percentages reported on the pie chart represent the proportion of cells within the specific cluster where *NDP* expression was observed. (**E**) Violin plots representing the localization of *FZD4* expression in week 7 and week 23 organoids. Each cell expressing *FZD4* is marked by a black dot. (**F**) Violin plots representing the localization of *LGR4* expression in week 7 and week 23 organoids. Each cell expressing *LGR4* is marked by a black dot. Abbreviations: KO, knockout; scRNA-seq, single cell RNA sequencing; UMAP, uniform manifold approximation and projection; WT, wildtype.

The analysis of a later timepoint (week 23) allowed for the identification of more cell types. Analysing the transcriptomic profiles of the clusters, 11 distinct major cell types were identified (Figure 3C): pRPC expressing marker genes such as *MKI67* and *TOP2A*, RPC with *SFRP2* and *HES1*, Müller glia cells (MGC) with *LGALS3* and *GLUL*, tRPC with *GADD45G* and *HES6*, PRp with *DDIT3* and *DDIT4*, rod photoreceptors (rod PR) with *NRL* and *NR2E3*, cone photoreceptors (cone PR)with *PDE6H* and *GNAT2*, ON and OFF bipolar cells (ON BC; OFF BC) with *NETO1* and *VSX1* or *PRDM8* and *VSX2* respectively, amacrine cells (AC) with *PAX6* and *TFAP2A*, and horizontal cells (HC) with *ONECUT1* and *PROX1* (Figure S08)[23,26,35337].

Notably, no RGC could be detected in week 23 organoids (Figure 3C), neither in *NDP*^WT^ nor *NDP*^KO^. Compared to the week 7 organoids, *NDP* was found to be expressed in an increased proportion of cells (11.4% in WT), mostly in the MGC (61.9%) and progenitor (RPC = 19.7% and pRPC = 7.8%) clusters (Figure 3D, Data 04).

### Localization of Norrin receptors in neuroretinal cell types

Norrin has been mainly described to act by binding to Frizzled-4, encoded by the *FZD4* gene, and activating WNT-signaling pathways stimulating the proliferation of vascular endothelial cells [9,38]. To understand which cell types could potentially be affected by Norrin signaling in the neuroretina, we analyzed the expression of *FZD4* transcripts in our scRNA-seq data. In week 7 retinal organoids, we detected *FZD4* expression in a small fraction of cells (average proportion of expressing cells: WT = 7.34%, KO = 7.62%). *FZD4* was expressed at higher levels in RPC and pRPC populations, while only lower levels were detected in the remaining cell types present at this developmental stage (Figure 3E, Figure S09A). In week 23 organoids, the proportion of cells expressing *FZD4* was found to be increased (average WT = 9.54%; KO = 8.41%). In addition to RPC and pRPC populations, higher levels of *FZD4* expression were detected in Cone PR, and HC, with traces also found in other cell types (Figure 3E, Figure S09A).

Alternatively, Norrin has been described to be able to bind also to the LGR4 receptor, a G-protein coupled receptor, that has been reported to inhibit negative regulators of Wnt-signaling upon activation [14,39]. Therefore, we sought to find which cells may be targets of Norrin signaling through the binding to LGR4 and found that in week 7 organoids the gene is expressed virtually ubiquitously (average proportion of expressing cells: WT = 89.98%, KO = 91.86%), with higher levels detected in RPC, pRPC, tRPC, H&A and eRGC (Figure 3F, Figure S09B). In week 23 organoids, *LGR4* expression was detected in a smaller proportions of cells compared to week 7 (average WT = 67.59%; KO = 64.42%). Also in this case, transcripts were found in all clusters, with higher levels in pRPC, RPC, MGC, HC, and AC (Figure 3F, Figure S09B).

### Transcriptomic alterations in Norrin-deficient human retinal organoids

To better understand the direct effect of Norrin on cells of the neural retina, we hypothesized that human retinal organoids might offer a suitable model, as these lack a vascular component that might confound or mask these effects. Therefore, we performed bulk RNA-seq on RNA extracted from pooled KO/MUT vs. WT retinal organoids (respectively: *NDP^KO^* and *NDP^MUT^*; *NDP^WT^* and *NDP^WTR^*) at weeks 7 and 23 of differentiation. Week 7 represents a differentiation stage in organoids, in which RGCs are still found in the organoids in a considerable number, followed by a progressive loss [40]. Additionally, we previously described how *NDP* transcripts were found in a subpopulation of *ATOH7*-expressing RGC [23]. Conversely, week 23 was chosen because at this time of organoids differentiation, both, late-born retinal neurons as well as mature MGC normally have already differentiated, representing a more mature model of the retina [41,42]. We performed differential expression analysis on the pseudoaligned reads (week 7: Data 05; week 23 Data 06) using DESeq2 and applied a significance cutoff of adjusted *p*-value < 0.05 [30], based on Benjamini-Hochberg false discovery rate (FDR), and found a total of 196

DEGs (128 upregulated and 68 downregulated in KO/MUT) in week 7 and 607 (344 upregulated and 263 downregulated in KO/MUT) in week 23 organoids (week 7: Figure 4A, Data 07; week 23: Figure 4C, Data08). Additionally, we extracted the differentially expressed genes from our scRNA- seq data and focused on genes with a significant difference between the WT and KO/MUT organoids (adjusted *p*-value < 0.05). In week 7 retinal organoids, we detected 228 DEGs (137 downregulated and 91 upregulated in KO/MUT; Figure S10A, Data 09), 18 of which overlapped with the bulk RNA-seq DEGs at the same developmental stage (Figure 4B). In week 23 organoids we found 933 DEGs (581 downregulated and 352 upregulated in KO/MUT; Figure S10B, Data 10), of which 94 overlapped with the ones detected in the week 23 bulk RNA-seq results (Figure 4D).

**Figure 4:**
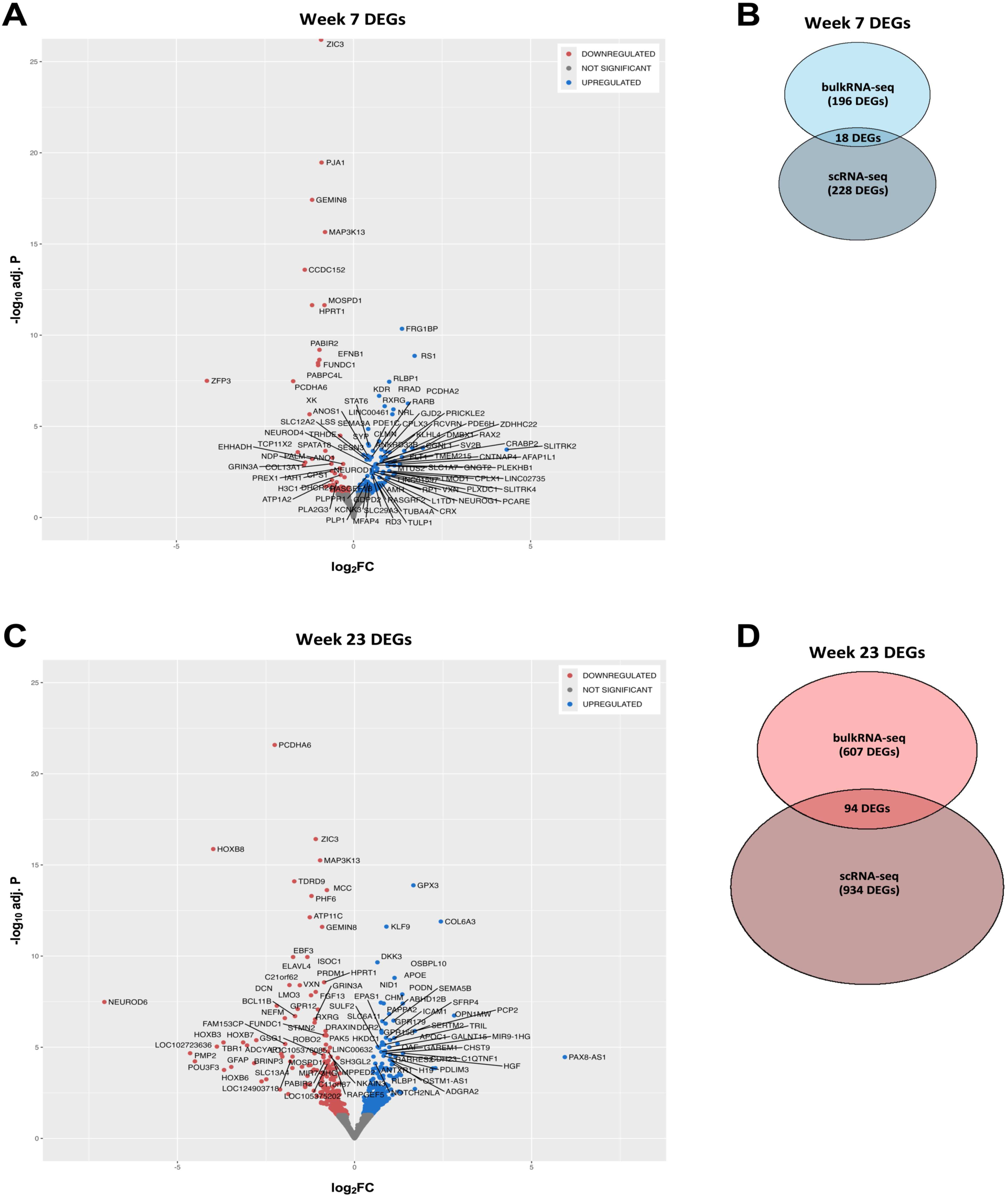
Differentially expressed genes (DEGs) in week 7 and week 23 organoids. (**A**) Volcano plot representing 196 DEGs (128 upregulated and 68 donwregulated in KO/MUT) detected in the bulk RNA-seq data in week 7 retinal organoids. A threshold of adjusted p-value of <0.05 was used. (**B**) Venn diagram representing the overlap of the bulk RNA-seq DEGs and the scRNA-seq DEGs. 18 DEGs were found in both lists. (**C**) Volcano plot representing the 607 DEGs (344 upregulated and 263 downregulated in KO/MUT) detected in the bulk RNA-seq data in week 23 retinal organoids. A threshold of adjusted p-value of <0.05 was used. (**D**) Venn diagram representing the overlap of the bulk RNA-seq DEGs and the scRNA-seq DEGs. 94 DEGs were found in both lists. Abbreviations: bulk RNA-seq, bulk RNA sequencing; DEGs, differentially expressed genes; FC, fold-change; KO, knockout; MUT, mutant reporter; scRNA-seq, single-cell RNA sequencing; WT, wildtype.

### Differential expression of genes associated with glutamate signaling and homeostasis in week 7 retinal organoids

Given the higher probability of gene dropout in scRNA-seq [43], we decided to focus our differential expression analyses on bulk RNA-seq data. Of the 196 DEGs identified in week 7 human retinal organoids, 128 were found to be upregulated and 68 downregulated in KO (Figure 4A, Data 07). The top 30 up- and downregulated genes are represented in Figure 5A. We annotated the DEGs using database for annotation, visualization and integrated discovery (DAVID; https://david.ncifcrf.gov, accessed 15.06.2024) and performed over-representation analysis (ORA) using WebGestalt (https://2024.webgestalt.org, accessed 07.06.2024). We were interested in the enrichment of Gene Ontology (GO) terms and pathways (KEGG, Reactome and WikiPathways). The top enriched terms of each category are listed in Figure 5B,C.

**Figure 5:**
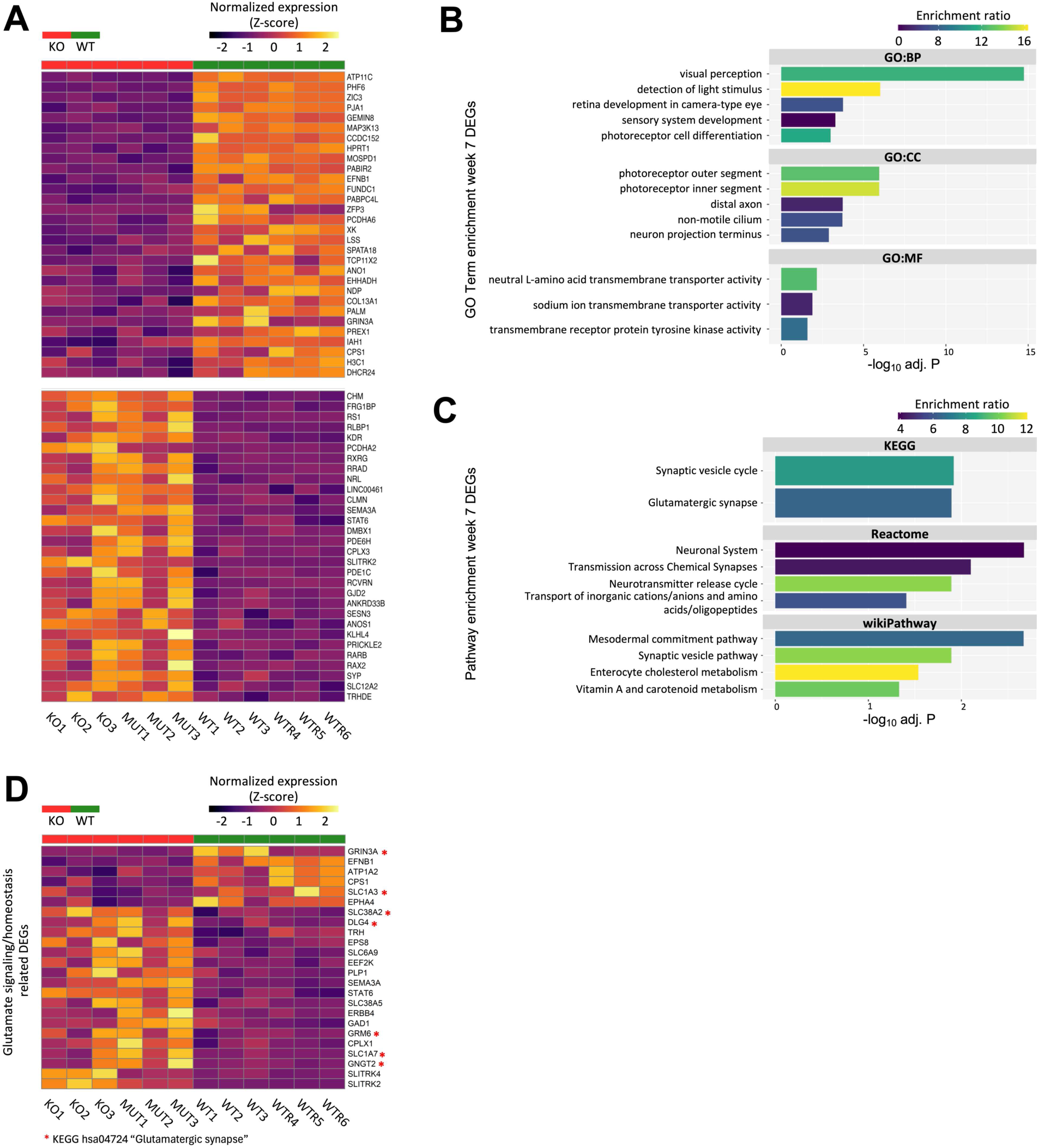
Differential expression in week 7 KO/MUT human retinal organoids. (**A**) Top 30 Up- and downregulated genes detected in week 7 retinal organoids. A threshold of adjusted p-value (FDR) of <0.05 was used. (**B**) Over-representation analysis (ORA) of the week 7 DEGs in GO terms (GO:BP, GO:CC, GO:MF), performed by WebGestalt, after reduction using REVIGO. The graphs show the top five terms according to the adjusted *p*-value. (**C**) Pathway ORA by WebGestalt of the DEGs detected in week 7 retinal organoids after reduction using REVIGO. The graphs show the top 5 terms, where applicable, according to the adjusted *p*-value. (**D**) Heatmap showing the DAVID annotated DEGs that were found to be related to the glutamate signaling system. *Genes detected by the ORA under the KEGG pathway term “Glutamatergic synapse” (hsa04724). Abbreviations: DEGs, differentially expressed genes; FDR, false discovery rate; GO:BP, GO biological process; GO:CC, GO cellular compartment; GO:MF, GO molecular function; KO, knockout; MUT, mutant; ORA, over-representation analysis; WT, wildtype; WTR, wildtype reporter.

Interestingly, among the enriched terms in ORA, we detected “glutamatergic synapse” (KEGG: hsa04724, FDR = 0.013; Figure 5C, Data 11). Glutamate is recognized as being one of the main neurotransmitter systems of the mammalian retina, where glutamatergic synapses are found between PRs-BCs as well as between BCs and RGCs [44347]. Interestingly, altered glutamate signaling and metabolism have been found as common underpinning of different conditions accompanied by degeneration of retinal cells, in particular of RGCs, such as ischemia and diabetic retinopathy [48,49]. Among the genes associated to the KEGG term in our data, ORA identified *SLC1A3*, *SLC1A7*, *SLC38A2*, *GNGT2*, *GRM6*, *DLG4* and *GRIN3A* (Figure 5D). Next, we expanded our search and included genes associated with glutamate signaling and metabolism terms (GO or pathway annotation) according to DAVID annotation, that were not detected by ORA, and found 17 additional DEGs (Figure 5D, Data 07). These results suggest that the absence or reduction *NDP* transcript levels results in transcriptional alterations of genes coding for proteins involved in glutamate signaling, its homeostasis and possibly also in the organization of glutamatergic synapses.

### Differential expression of ECM-related genes in week 23 retinal organoids

In week 23 organoids, we detected an increased number of DEGs with a total 607, 344 of which were upregulated and 263 were downregulated in KO/MUT organoids (Figure 4C, Data 08). The top 30 up and downregulated genes found in our bulk RNA-seq data are presented in Figure 6A. We repeated DAVID annotation and performed overrepresentation analysis using WebGestalt, looking for enrichment in GO terms or in pathways.

**Figure 6:**
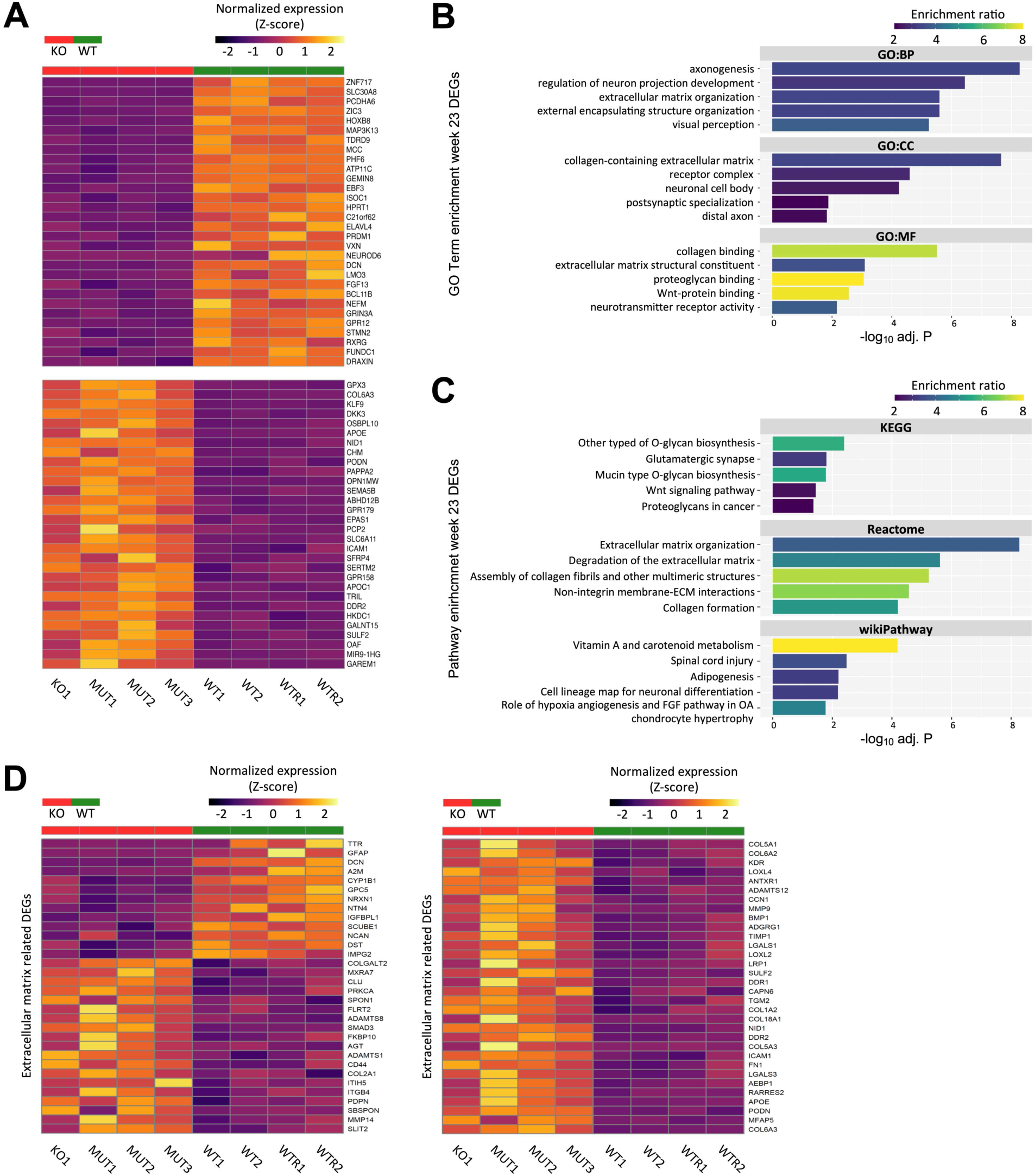
Differential expression in week 23 KO/MUT human retinal organoids. (**A**) Top 30 Up- and downregulated genes detected in week 23 retinal organoids according to the adjusted *p*-value. A threshold of adjusted *p*-value (FDR) <0.05 was used. (**B**) Over-representation analysis (ORA) of the week 23 DEGs in GO terms (GO:BP, GO:CC, GO:MF), performed by WebGestalt, after reduction using REVIGO. The graphs show the top five terms according to the adjusted *p*-value. (**C**) Pathway ORA by WebGestalt of the DEGs detected in week 23 retinal organoids after reduction using REVIGO. The graphs show the top five terms according to the adjusted *p*-value. (**D**) Heatmap showing the DAVID annotated DEGs that were found to be related to the extracellular matrix (ECM) after integration of lists related to different GO terms or pathways detected by ORA (e.g. “extracellular matrix organization”, GO0030198). Abbreviations: DEGs, differentially expressed genes; ECM, extracellular matrix; FDR, false discovery rate; GO:BP, GO biological process; GO:CC, GO cellular compartment; GO:MF, GO molecular function; KO, knockout; MUT, mutant; ORA, over-representation analysis WT, wildtype; WTR, wildtype reporter.

Among the enriched terms found in ORA performed with the week 23 DEGs as input list, we found an enrichment of terms related to the ECM or its regulation and organization. Specifically, among the top enriched GO terms, we detected “extracellular matrix organization” (GO:BP, GO:0030198), “collagen-containing extracellular matrix” (GO Cellular Component, GO:0062023) or “collagen binding” (GO Molecular Function, GO:0005518) (Data 12). Enrichment of extracellular matrix-related terms was also found in the top enriched pathways, with terms like “other types of O-glycan biosynthesis” (KEGG, hsa00514), “degradation of the extracellular matrix” (Reactome, R-HSA1474228) or “collagen formation” (Reactome, R-HSA-147290) (Figure 6B,C). Among the DEGs related to this category of terms, we found several coding for metallopeptidases (*ADAMTS1*, *ADAMTS12*, *ADAMTS8*, *MMP9* and *MMP14*), as well as several different collagen coding genes (*COL1A2*, *COL2A1*, *COL5A1*, *COL5A3*, *COL6A2*, *COL6A3* and *COL18A1*) and other genes associated with the ECM (Figure 6D).

Intriguingly, also in week 23 DEGs, we found enrichment of genes related to glutamate signaling and homeostasis. We found 35 DEGs that were related to glutamate signaling, its regulation, and homeostasis, based on DAVID annotation and confirmed by literature research (Figure S11). Surprisingly, only two of these were found in both timepoints, namely *GRIN3A* and *GAD1*. This may also be related to the differentiation stage of the organoids. Indeed, at week 23 more cell types have differentiated and matured, likely contributing to the differences in the transcriptomic profiles of the samples. In addition, some compensatory mechanism may have set in contributing to the differences detected.

These results suggest that lack or reduced levels of *NDP* transcript may result in the dysregulation of genes related to the composition, organization, and regulation of the ECM and possibly in transcriptomic alterations of genes involved in glutamate signaling.

### *NDP^KO^* human retinal organoids display a shift in cell composition towards RPCs and MGCs

The direct involvement of Norrin in developmental processes that do not relate to the vascular system has been so far described in early ectodermal specification in *Xenopus laevis*, in cochlear hair cell maturation and during decidualization in mice [15,19,50]. Furthermore, cytoarchitectural abnormalities, such as a disorganization and early loss of cells in the RGCs layer have been previously described in the *Ndp^KO^* mouse model [16,17]. Additionally, given the distribution of Norrin receptors in retinal organoids (Figure 3E,F), we hypothesize that Norrin- signaling may affect more cell populations than previously reported, potentially exerting yet undescribed effects. Therefore, we analyzed the cellular composition in the scRNA- seq data.

Although no significant differences were detected between *NDP^KO^* and *NDP^WT^* organoids at week 7, we found a higher proportion of RPC in KO organoids compared to WT (+18,3%). Conversely, cells belonging to the remaining clusters were generally decreased in KO, with the highest differences found in H&A (-11,6%), eRGC (-10,5%), and lRGC (-26,1%) (Figure 7A, Figure S12A,B, Data 13). Next, we analyzed the composition of the week 23 organoids and found significant differences in the proportions of cells belonging to the RPC, MGC, and HC clusters. We observed that RPC and MGC were overrepresented in KO organoids at this timepoint, with a 34,1% increase in RPC (t-test *p*-value = 0.017) and a 27,3% increase in MGC (t-test *p*-value = 0.029). HC were found to be underrepresented in the KO organoids, with a 33.3% decrease (t-test *p*-value = 0.018)(Figure 8B, Figure S13A,B, Data 14). Except for OFF BC and AC, all other clusters were generally underrepresented in KO organoids compared to their WT counterparts.

**Figure 7:**
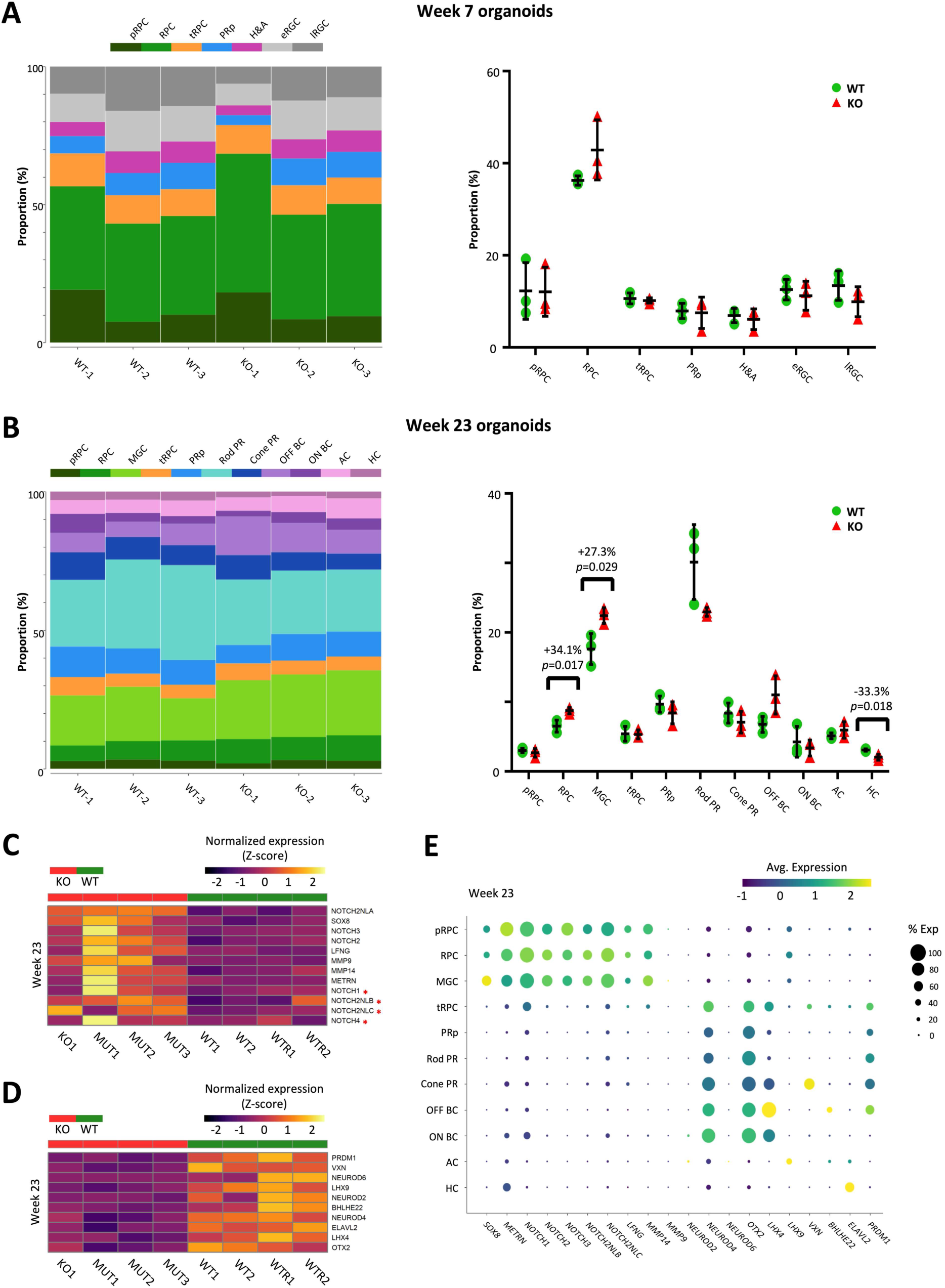
Cell composition of *NDP^KO^* retinal organoids and neurogenesis-related DEGs. (**A**) Representation of the proportional cell composition of week 7 retinal organoids analyzed by scRNA-seq. The values represent the percentage of the total cells in the respective cluster. (**B**) Representation of the proportional cell composition of week 23 retinal organoids analyzed by scRNA-seq. The values represent the percentage of the total cells in the respective cluster. Significant differences were detected in the RPCs, MGCs, and HCs clusters (t-test p-value <0.05). (**C**) Normalized expression at week 23 of genes associated with either maintenance of a progenitor population pool (NOTCH signaling-related genes) or with the specification of MGC identity (*SOX8* and *METRN*). Genes sorted according to the adj. *p*-value. *Upregulated (log2FC>0), but not significant (adjusted p-value >0.05). (**D**) Normalized expression at week 23 of DEGs associated with neurogenesis and specific retinal neuronal identities. Genes sorted according to the adj. *p*-value. (E) Dot-plot displaying the cell type specific expression in week 23 scRNA-seq data of the genes found in the heatmap, showing the proportion of cells expressing them and the average expression in each cluster. Abbreviations: DEGs, differentially expressed genes; FC, fold-change; HCs, horizontal cells; KO, knockout ; MUT, mutant; MGCs, Müller glia cells; RPCs, retinal progenitor cells; scRNA-seq, single-cell RNA sequencing, WT, wildtype; WTR, wildtype reporter.

The difference in composition of KO human retinal organoids is compatible with a possible role of Norrin in neuroretinal development and differentiation independent from its role in promoting vascular development. In week 23 organoids, we detected a tendency of RPC as well as MGC populations to be overrepresented in KO organoids, with an underrepresentation of neuronal populations. For this reason, we speculated that there may be transcriptomic changes underlying this shift in cell populations. Among the week 23 DEGs, we detected upregulation, although not always significant, of NOTCH receptors, *NOTCH1* (adj. *p*-value = 0.187), *NOTCH2* (adj. *p*-value = 0.010), *NOTCH3* (adj. *p*-value = 0.008) and *NOTCH4* (adj. *p*-value = 0.378), as well as some NOTCH signaling modulators, *LFNG* (*adj. p-value* = 0.029), *MMP9* (adj. p-value = 0.030)*, MMP14* (adj. *p*- value = 0.032), *NOTCH2NLA* (*adj. p-value* = 7.85x10^-5^), *NOTCH2NLB* (adj. p-value = 0.229), *NOTCH2NLC* (adj. *p*-value = 0.268) (Figure 7C, Data 08). Interestingly, NOTCH signaling is required for maintaining RPC in an undifferentiated proliferative state, thus slowing the generation of late-born neuroretinal cells [51354]. Moreover, we found significant upregulation in the KO organoids of *SOX8* (*adj. p-value* = 0.002), and *METRN* (*adj. p-value* = 0.037)(Figure 7C, Data 08). *SOX8* codes for the transcription factor SOX-8, whereas *METRN* codes for Meteorin, a secreted neurotrophic factor. Both proteins have been described to be involved in the differentiation of MGC [55,56]. Interestingly, NOTCH signaling has also been demonstrated to promote differentiation of glial cells from neural multipotent progenitors in the central nervous system and of MGC from both retinal progenitor cells and human embryonic stem cells (hESCs) [57359]. Next, we were interested in identifying where these genes were expressed in human retinal organoids in the scRNA-seq data. Expression of *NOTCH1*, *NOTCH2*, *NOTCH3*, *NOTCH2NLB*, *NOTCH2NLC*, *LFNG*, *MMP9*, *MMP14*, *SOX8* and *METRN* was detected at different levels in pRPC, RPC and MGC (Figure 7E). Conversely, among significantly downregulated genes, we detected several genes coding for factors associated with the differentiation of neuronal cell-types of the retina such as *OTX2, NEUROD2, NEUROD4, NEUROD6*, *LHX4*, *LHX9*, *VXN*, *BHLHE22*, *ELAVL2* and *PRDM1* (Figure 7E, Data 08). Together, these observations suggest that Norrin may be involved in the regulation of neurogenesis during retinal development, possibly supporting differentiation of RPC towards neuronal cell types such as photoreceptors (PR), RGC HC, AC and BC.

## Discussion

### *NDP* is expressed in RPC, MGC, and a subset of RGC

The exact source of Norrin in the human retina is currently still not known. The first study describing *Ndp* expression used mRNA in-situ hybridization in mouse eyes and reported signals in the INL and RGC layer, suggesting that RGC may be the main source of the protein in the retina [16]. While others have reported expression in the outer nuclear layer (ONL) and inner nuclear layer (INL) [60]. More recently, researchers studying an *Ndp* AP reporter mouse line, detected signals spanning the whole thickness of the neuroretina at P4, P7 and in adult mice, suggesting that MGC may be the principal source of the protein [12]. Based on our scRNA-seq data, we demonstrate that during early (week 7) human embryonic retinal development *NDP* is mainly expressed in RPCs populations and a subset of RGCs (Figure 3B). Intriguingly, in one of our previous scRNA-seq studies, we found that *NDP* was significantly downregulated in *ATOH7^KO^* human retinal organoids, and that *NDP* expression in the RGC was restricted to *ATOH7*-expressing cells [23]. These findings suggest a possible link between the two genes. Phenotypic similarities between the conditions associated with pathogenic sequence variants in *ATOH7* and *NDP* support this idea.

In week 23 human retinal organoids, *NDP* transcripts were mainly detected in RPC populations and MGC (Figure 3D), indicating that MGC are the main source of Norrin during later stages of human retinal development [12]. It is important to mention that human retinal organoids are lacking RGC at this timepoint (Figure 3C). This is a known phenomenon, since RGC are indeed progressively lost during long-term culture. They are located at the innermost cell layer of the organoids, where diffusion of oxygen and nutrients is limited [40]. For this reason, we cannot exclude RGC as a source of Norrin in later stages of human retinal differentiation in addition to MGC and RPC.

### Norrin affects different signaling pathways and ECM components

To the best of our knowledge, this is the first study exploring the global gene expression patterns resulting from the lack of Norrin in a human organoid model. A literature search allowed us to identify four previous publications that reported transcriptomic data derived from mouse models (*Ndp^KO^*) of the disease [19,61363] . Lenzner et al. analyzed the global gene expression patterns in the adult stage retina (2 years of age), whereas Schäfer et al. focused on young mouse pups at postnatal day 7 (P7). In both studies, microarrays were used to analyze gene expression [61,62]. Hayashi et al. and Pauzuolyte et al. focused on the cochlear transcriptomic differences applying RNA-seq in a subpopulation of cochlear cells isolated at P4-P6 in mice, and whole cochleae collected at 2 months of age, respectively [19,63]. We compared the lists of DEGs reported in these studies with our week 7 and week 23 DEGs, which revealed some overlapping genes. Within our week 7 DEGs, we found for *PDC*, *RCVRN*, *IMPG1*, *RS1*, *SLC38A5*, *CNTNAP4* and *FLT1*, whereas within the week 23 DEGs we detected *OPN1SW*, *GNAT2*, *BAG3*, *ARHGAP10*, *A2M*, *SLA*, *CNTNAP4*, *FLT1*, *CLU* and *AEBP1* (Figure S14A,B). Despite the restricted number of identical genes present in the different studies, we found an overlap with Wnt signaling related terms at both timepoints, week 7 and week 23 (Figure 6B,C; Figure S14C; Data 11,12), which had been described previously [19,62]. Additionally, an overlap was found with MAPK signaling pathway terms (Figure S14D; Data 12), which had been reported previously as being the top-ranking affected signaling pathway affected in the retinae of P7 *Ndp^KO^* mice [62].

Interestingly, in week 23 KO/MUT organoids we found differential expression of genes coding for proteins of the ECM, and for proteins involved in its organization and regulation (Figure 6D, Data 12). The ECM is a complex and dynamic network of proteins and other molecules that participate in several developmental processes in the retina, including cell proliferation, migration, adhesion, differentiation, and maturation [64]. Additionally, it plays an essential role in the growth of cellular extensions, a process involved in both axonal growth and guidance and angiogenesis [65,66]. Alterations affecting the ECM can have severe detrimental consequences on different aspects of retinal development. One of the first morpho-histological findings reported in the *Ndp^KO^* mouse model was a cellular disorganization in the RGC layer, with displacement of the cell bodies and nuclei towards the INL [16,17]. This observation may be compatible with a defect in the migration of the differentiated or differentiating RGCs. Alterations related to the ECM, which we detected, could play a role in the migration of differentiating retinal cells towards their final position. Moreover, secreted signaling molecules (e.g. Norrin), diffuse through the ECM before reaching their targets. Norrin has been described being tightly associated with specific ECM components, and specific pathogenic variants were found to increase or decrease the amount of the protein detectable in the ECM [67].

We therefore conclude that our data indicate that different signaling pathways, such as the Wnt and MAPK signaling pathway, as well as the composition, the regulation and the organization of the ECM are affected when Norrin is absent or decreased. Our data are consistent with previously reported findings, which described differential expression of genes involved in these pathways in the *Ndp^KO^* mouse [19,62].

### Norrin may be involved in the differentiation of late-born retinal neurons

Until now, direct consequences of diminished or missing Norrin signaling on neuronal cell types have been elusive. The possibility that Norrin plays a direct role in the development and differentiation of retinal neuros independent from its role in angiogenesis has been hypothesized previously [16]. Berger et al described a disorganization in the RGCs-INL and occasional morphological abnormalities in the ONL and photoreceptor outer segments regions in retinae of *Ndp^KO^* mice [16]. These mice were later found to have an increased number of proliferating retinal cells and, in particular, an increased number of RGCs when Norrin was ectopically expressed in the lens, suggesting the possibility that Norrin is involved in the differentiation of retinal neurons [68]. Additionally, Norrin has been found to be essential for the development and maintenance of hair cells in the cochlea [19]. Moreover, both canonical Wnt-signaling as well as LGR4-mediated signaling, which also acts through alternative signaling pathways [69], have been implicated in multiple developmental processes [70].

In this context, our data indicate an arrest of neuronal differentiation in human retinal organoids in the absence of Norrin and a significant overrepresentation of RPCs and MGC in the week 23 *NDP^KO^* organoids as compared to WT ones (Figure 7A,B). Intriguingly, several genes involved in NOTCH signaling, which we found to be expressed mainly in pRPC, RPC and MGC, were upregulated in KO/MUT organoids (although not all to a significant level; Figure 7C,E). This could likely result in the arrest of neuronal differentiation, as NOTCH signaling has been reported to be required to maintain neural progenitor cells in an undifferentiated state through lateral inhibition [51354]. The secreted factors NOTCH2NL-A, -B, and -C have been found to delay neuronal differentiation in hESCs-derived cortical organoids by enhancing NOTCH signaling, resulting in upregulation of genes involved in negative regulation of neurogenesis [71]. Furthermore, the signaling pathway has been implicated in the differentiation of glial cells in the CNS in rats, and of MGC in the mouse retina [57,58]. Media supplementation with NOTCH ligands was described as sufficient to induce hESCs-derived RPC commitment to an MGC fate [59]. Moreover, we detected *METRN* and *SOX8* among the upregulated genes (Figure 7C), both of which have been implicated in the differentiation of MGC [55,56]. These observations support the idea that Norrin signaling may be involved in neuronal fate commitment.

In KO/MUT organoids, we detected downregulation of several genes that have been implicated in neurogenesis, and specifically to the commitment to specific neuronal fates in the retina (Figure 7E,D). Of these genes, *OTX2*, *VXN,* and *PRDM1* have been reported to participate in PR specification [72374]. On the other hand, *NEUROD2*, *NEUROD6*, *BHLHE22*, *LHX9*, and *ELAVL2* were implicated in differentiation of AC and their sub-types [75379]. *NEUROD4*, *LHX4*, BHLHE22. and *VXN* have been described to promote BCs identity [73,76,77,80]. Interestingly, *NEUROD2*, *NEUROD4*, *NEUROD6.* and *BHLHE22* are members of the bHLH transcription factors family. These are known downstream targets of NOTCH signaling through the transcriptional repressors of the HES family [81].

In summary, our data suggest an arrest or delay of neuronal differentiation in human retinal organoids due to the lack or reduction of *NDP* transcripts. The exact underlying mechanisms remain to be elucidated, but may involve altered transcription of genes encoding for NOTCH signaling components and other pro-neural factors.

### Alterations of Norrin levels may affect glutamate signaling and homeostasis

In both timepoints, week 7 and 23, we detected enrichment of DEGs associated with GO/pathway terms related to <glutamate signaling and homeostasis= (Figure 5B-D; Figure S11; Data 07,08). Strikingly, retinal angiogenesis as well as the maturation of the blood-retinal barrier (BRB) were found to be regulated by neuronal glutamatergic activity through the modulation of Norrin/³- Catenin signaling. The lack of glutamate release was associated with a delay in angiogenesis and BRB maturation, while constitutive glutamate release led to acceleration of angiogenesis and BRB maturation. This was associated with down- and up-regulation of Norrin expression, respectively [82].

Alterations in glutamate signaling may also explain why RGC seem to be the main cell type affected in the neuroretina in the absence of Norrin. In *Ndp^KO^* mice, disorganization and early loss of cells in the RGCs layer have been described [16,17]. Later studies reported a neuroprotective role of the protein in different conditions. Seitz et al found that Norrin protects RGC from NMDA-mediated damage [83]. Later, the same group reported an increased number of RGC in a mouse model of glaucoma, when Norrin was constitutively expressed in the retina under the control of a *Pax6* fusion promoter [84]. Neuroprotective effects on RGCs were also reported in a model of retinal ischemia, where increased numbers of RGCs survived the ischemic conditions after intravitreal delivery of Norrin [85]. Both, glaucomatous as well as ischemic conditions have been proposed to lead to unbalanced levels of glutamate, resulting in over-activation of NMDA receptors and excitotoxic damage [86389]. In this context, we found the downregulation of *GRIN3A* in our *NDP^KO^* organoids at both timepoints of particular interest. *GRIN3A* codes for the GluN3A subunit of GluN3A-NMDA receptors (NMDAr). Compared to conventional NMDAr, GluN3A-NMDAr have an inhibitory effect on conventional NMDAr-mediated activity, due to their substantially lower Ca^2+^ permeability [90]. Interestingly, the gene has also been described to have neuroprotective functions in excitotoxic and ischemic conditions [91,92]. In particular, cultivated cortical neurons from newborn *Grin3a^KO^* mice were found to be more sensitive to NMDA-mediated damage. Strikingly, in the same study, NMDA was injected in the retina of adult mice, where a higher level of Grin3a expression persists, and a decreased amount of surviving RGCs were detected in *Grin3a*-deficient mice [91,92].

Our results suggest that Norrin may be involved, directly or indirectly, in maintaining glutamatergic activity as a prerequisite for normal vascular development and BRB maturation in the human retina. Moreover, missing or decreased levels of the protein may confer higher susceptibility of neuronal cells (in particular RGC) to glutamate-mediated damage. The previously reported effect of glutamate on Norrin/³-Catenin signaling, along with our finding that altering the transcription levels of *NDP* interferes with the transcription of genes involved in glutamate signaling suggests a bidirectional modulation of the two pathways.

### Limitations

Our results were obtained from iPSC-derived human retinal organoids at weeks 7 and 23. It has been described previously that RGCs are lost during long-term organoid culture [40]. Therefor we cannot exclude additional effects on this cell type during late stages of differentiation. In addition, retinal organoids lack virtually all components of the vascular and immune systems [93]. In our case, we took advantage of this limitation to identify the effects of Norrin on neuroretinal differentiation exclusively.

Additionally, qPCR and bulkRNA-seq data revealed the presence of residual WT *NDP* transcripts in the MUT organoids. Nevertheless, this was only approximately 2% in week 7 organoids and 20% in week 23 organoids. Furthermore, we performed additional analyses for the separate datasets, using only WT vs KO, WTR vs MUT and scRNA-seq DEGs, which revealed a certain degree of overlap between the datasets (Data S11 and Data S12). For these reasons we decided to maintain the analysis of the pooled samples to reveal the effects of a strong reduction of *NDP* transcripts.

The scRNA-seq method used in this study relies on probe hybridization-based capturing of transcripts, which may increase the rate of dropouts or false positives. Also, we limited our analysis to two timepoints in differentiation (week 7 and week 23). Additional developmental timepoints may provide a more detailed view of the molecular processes involved in neuroretinal development and differentiation. Nevertheless, we consider this model a valuable tool to study the function of Norrin in neuronal differentiation.

### Conclusions

Here we describe, for the first time, a putative role of Norrin in the differentiation of retinal neurons in a human retinal organoids model derived from hiPSC. Our data support the findings of previous studies, which described transcriptional alterations in genes related to Wnt- and MAPK signaling, as well as in genes associated to the ECM. In addition, we detected a significant increase in the proportions of RPC and MGC in *NDP^KO^* organoids. This was associated with a reduction in the proportions of other neuronal retinal populations, including PR, BC, and HC. The upregulation of several genes associated with NOTCH signaling and a downregulation of pro- neural factors that were detected in the KO/MUT organoids, may explain the neurogenic arrest or delay in *NDP^KO^* organoids.

## Author contributions

**Conceptualization**: Kevin Maggi, David Atac, Samuel Koller, Wolfgang Berger; **Data curation**: Kevin Maggi; **Formal Analysis**: Kevin Maggi; Funding: Wolfgang Berger, David Atac, Jordi Maggi; **Investigation**: Kevin Maggi, David Atac, Silke Feil, Jordi Maggi; **Methodology**: Kevin Maggi, David Atac; **Project administration**: Wolfgang Berger; **Resources**, Wolfgang Berger; **Software**: Wolfgang Berger; **Supervision**: Wolfgang Berger, Kevin Maggi, David Atac; **Validation**: Kevin Maggi; **Visualization**: Kevin Maggi; **Writing 3 original draft**: Kevin Maggi; **Writing 3 review & editing**: Wolfgang Berger; Samuel Koller, Jordi Maggi, David Atac.

## Institutional Review Board Statement

Not applicable.

## Informed Consent Statement

Not applicable.

## Acknowledgements

We are grateful to Jane Sowden and Elisa Cuevas for sharing their retinal organoids differentiation protocol with us. We would also like to acknowledge the Functional Genomics Center Zurich for performing high-throughput sequencing experiments and support in analyses, with special thanks to Qin Zhang, Susanne Kreutzer, and Lennart Opitz. We express our gratitude to the Center for Microscopy and Image Analysis at the University of Zurich, in particular, to Tatiane Groski and Philipp Schätzle for their assistance during usage of equipment.

