## Supplementary material for "Putative Role of Norrin in Neuroretinal Differentiation Revealed by bulk and scRNA Sequencing of Human Retinal Organoids": Data01_NDP-WT-GFP_repair-template.pdf

```
                                                        10        20        30        40        50        60        70        80        90       100       110
                                                ACGCGGCCTTTTTACGGTTCCTGGCCTTTTGCTGGCCTTTTGCTCACATGTTCTTTCCGATTCCTTTTTTCCATTCCGTTAAGCTGTATGCAAATTGTGTGTGTGCTTAT
H12-W59500-U029XGJ200-4-1.8K-SEQR.ab1(1>540)  → ACGCGGCCTTTTTACGGTTCCTGGCCTTTTGCTGGCCTTTTGCTCACATGTTCTTTCCGATTCCTTTTTTCCATTCCGTTAAGCTGTATGCAAATTGTGTGTGTGCTTAT
U029XGJ200-4.seq(1>2973)                      →                                                              TCCTTTTTTCCATTCCGTTAAGCTGTATGCAAATTGTGTGTGTGCTTAT
E02-W59500-U029XGJ200-4-4-SEQ2R.ab1(1>823)    ←                                                                               TTAAGCTGTATGCAAATTGTGTGTGTGCTTAT

                                                       120       130       140       150       160       170       180       190       200       210       220
                                                GTACATTTTTCTGGGGATAACTTTGGTAGTTTTCATCAGATTCTTAAAGGTGTCTGTTTGCTAAAGCCAGATTAAGAGCTACTATAAAGAACACATAGCCTCCGTTATTC
H12-W59500-U029XGJ200-4-1.8K-SEQR.ab1(1>540)  → GTACATTTTTCTGGGGATAACTTTGGTAGTTTTCATCAGATTCTTAAAGGTGTCTGTTTGCTAAAGCCAGATTAAGAGCTACTATAAAGAACACATAGCCTCCGTTATTC
U029XGJ200-4.seq(1>2973)                      → GTACATTTTTCTGGGGATAACTTTGGTAGTTTTCATCAGATTCTTAAAGGTGTCTGTTTGCTAAAGCCAGATTAAGAGCTACTATAAAGAACACATAGCCTCCGTTATTC
E02-W59500-U029XGJ200-4-4-SEQ2R.ab1(1>823)    ← GTACATTTTTCTGGGGATAACTTTGGTAGTTtTCATCAGATTCTTAAAGGTGTCTGTTTGCTAAAGCCAGATTAAGAGCTACTATAAAGAACACATAGCCTCCGTTATTC
F10-W59500-U029XGJ200-4-4-SEQ1F.ab1(1>423)    →      TTTTTCTGGGGAtAACTTTGGTAGTTTTCATCAGATTCTTAAAGGTGTCTGTTTGCTAAAGCCAGATTAAGAGCTACTATAAAGAACACATAGCCTCCGTTATTC

                                                       230       240       250       260       270       280       290       300       310       320       330
                                                CAGCCCAATCTTTGGAAATAGAGATTATCAGAATGGGTACATCCTAATATTTGCCGCAAATGACTCTGAAGATGCGGATGCAATTACATAAATCCCAGGCTCCTTCAACA
H12-W59500-U029XGJ200-4-1.8K-SEQR.ab1(1>540)  → CAGCCCAATCTTTGGAAATAGAGATTATCAGAATGGGTACATCCTAATATTTGCCGCAAATGACTCTGAAGATGCGGATGCAATTACATAAATCCCAGGCTCCTTCAACA
U029XGJ200-4.seq(1>2973)                      → CAGCCCAATCTTTGGAAATAGAGATTATCAGAATGGGTACATCCTAATATTTGCCGCAAATGACTCTGAAGATGCGGATGCAATTACATAAATCCCAGGCTCCTTCAACA
E02-W59500-U029XGJ200-4-4-SEQ2R.ab1(1>823)    ← CAGCCCAATCTTTGGAAATAGAGATTATCAGAATGGGTACATCCTAATATTTGCCGCAAATGACTCTGAAGATGCGGATGCAATTACATAAATCCCAGGCTCCTTCAACA
F10-W59500-U029XGJ200-4-4-SEQ1F.ab1(1>423)    → CAGCCCAATCTTTGGAAATAGAGATTATCAGAATGGGTACATCCTAATATTTGCCGCAAATGACTCTGAAGATGCGGATGCAATTACATAAATCCCAGGCTCCTTCAACA

                                                       340       350       360       370       380       390       400       410       420       430       440
                                                TATAAACCCAGAAGTGTGATTTCAGGGCCCCAGACATGCCCTGTCTCTACTTCATTCTCTTGTCCCAAATTGCAGCACCCAGAATATTTCTTCCTAAGCATTACTCAGGC
H12-W59500-U029XGJ200-4-1.8K-SEQR.ab1(1>540)  → TATAAACCCAGAAGTGTGATTTCAGGGCCCCAGACATGCCCTGTCTCTACTTCATTCTCTTGTCCCAAATTGCAGCACCCAGAATATTTCTTCCTAAGCATTACTCAGGC
U029XGJ200-4.seq(1>2973)                      → TATAAACCCAGAAGTGTGATTTCAGGGCCCCAGACATGCCCTGTCTCTACTTCATTCTCTTGTCCCAAATTGCAGCACCCAGAATATTTCTTCCTAAGCATTACTCAGGC
E02-W59500-U029XGJ200-4-4-SEQ2R.ab1(1>823)    ← TATAAACCCAGAAGTGTGATTTCAGGGCCCCAGACATGCCCTGTCTCTACTTCATTCTCTTGTCCCAAATTGCAGCACCCAGAATATTTCTTCCTAAGCATTACTCAGGC
F10-W59500-U029XGJ200-4-4-SEQ1F.ab1(1>423)    → TATAAACCCAGAAGTGTGATTTCAGGGCCCCAGACATGCCCTGTCTCTACTTCATTCTCTTGTCCCAAATTGCAGCACCCAGAATATTTCTTCCTAAGCATTACTCAGGC

                                                       450       460       470       480       490       500       510       520       530       540       550
                                                TATCCTATTTTCTTATTCAGAAACTATATTCCTGTTCTTTAGCTATTTCCTGCACAAATCAATTTCTTTTTTTTTTTTTTTTTTTTGGTAGTCATTTTAATTTACTTCTT
H12-W59500-U029XGJ200-4-1.8K-SEQR.ab1(1>540)  → TATCCTATTTTCTTATTCAGAAACTATATTCCTGTTCTTTAGCTATTTCCTGCACAAATCAATTTCTTTTTTTTTTTTTTTTTTTTGGtAgTCatTTTAA
U029XGJ200-4.seq(1>2973)                      → TATCCTATTTTCTTATTCAGAAACTATATTCCTGTTCTTTAGCTATTTCCTGCACAAATCAATTTCTTTTTTTTTTTTTTTTTTTTGGTAGTCATTTTAATTTACTTCTT
E02-W59500-U029XGJ200-4-4-SEQ2R.ab1(1>823)    ← TATCCTATTTTcTTATTCAGAAACTATATTCCTGTTCTTTAGCTATTTCCTGCaCAAATCAATTTcTTTTTTTTTTTTTTTTTTTTGGTAGTCATTTTAATTTACTTCTT
F10-W59500-U029XGJ200-4-4-SEQ1F.ab1(1>423)    → TATCCTATTTTCTTATTCAGAAACTATATTCCTGTTCTTTAGCTATTTCCTGCACAAATCAATTTCTTTTTTTTTTTTTTTTTTTTGGtAgTCaTTTT

                                                       560       570       580       590       600       610       620       630       640       650       660
                                                GGTTTCAATTTGGATTGACTTTCTGCATTTTGCTACTGGTAGAATTGCAATGCATTGGGCCAGTGGTAGACCACTACCTGTCAGGAGAAATTTCTTTTGTTCTTTGGTTT
U029XGJ200-4.seq(1>2973)                      → GGTTTCAATTTGGATTGACTTTCTGCATTTTGCTACTGGTAGAATTGCAATGCATTGGGCCAGTGGTAGACCACTACCTGTCAGGAGAAATTTCTTTTGTTCTTTGGTTT
E02-W59500-U029XGJ200-4-4-SEQ2R.ab1(1>823)    ← GGTTTCAATTTGGATTGACTTTCTGCATTTTGCTACTGGTAGAATTGCAATGCATTGGGCCAGTGGTAGACCACTACCTGTCAGGAGAAATTTCTTTTGTTCTTTGGTTT

                                                       670       680       690       700       710       720       730       740       750       760       770
                                                TTACTCGCACATCACAGTGATGGGCGTCATTCAGTGGTTCTAAACCCTCAGTACACTTAGAAATCACCTAAGAAGCTTTGAAAACCATGAAAGCCTGGTCCCTATCTCCA
U029XGJ200-4.seq(1>2973)                      → TTACTCGCACATCACAGTGATGGGCGTCATTCAGTGGTTCTAAACCCTCAGTACACTTAGAAATCACCTAAGAAGCTTTGAAAACCATGAAAGCCTGGTCCCTATCTCCA
E02-W59500-U029XGJ200-4-4-SEQ2R.ab1(1>823)    ← TTACTCGCACATCACAGTGATGGGCGTCATTCAGTGGTTCTAAACCCTCAGTACACTTAGAAATCACCTAAGAAGCTTTGAAAACCATGAAAGCCTGGTCCCTATCTCCA
```

```
                                                   780       790       800       810       820       830       840       850       860       870       880
                                                 AGGATTTTAATTCAATCGATCTGAATGAGGCCTGGGTGTTGACTTAAAAAAAATGCAACATGCATTGAGCCACTGGTCTAATCTAGCAGGAATTGGCAACGAGTGTGAGT
U029XGJ200-4.seq(1>2973)                     →   AGGATTTTAATTCAATCGATCTGAATGAGGCCTGGGTGTTGACTTAAAAAAAATGCAACATGCATTGAGCCACTGGTCTAATCTAGCAGGAATTGGCAACGAGTGTGAGT
E02-W59500-U029XGJ200-4-4-SEQ2R.ab1(1>823)   ←   AGGATTTTAATTCAATCGATCTGAATGAGGCCTGGGTGTTGACTTAAAAAAAATGCAACATGCATTGAGCCACTGGTCTAATCTAGCAGGAATTGGCAACGAGTGTGAGT
E03-W59500-U029XGJ200-4-4-SEQ3F.ab1(1>825)   →                                                                                        GCAGGAATTGGCAACGAGTGTGAGT

                                                   890       900       910       920       930       940       950       960       970       980       990
                                                 GTCTTTCAAAAGATGCTGTTTTACCTGGCTAAGGTTGTGGCATGCCCACAGAGTAACCACCTCCTTTTCTTTCCCCAGATGGTGCTCCTGGCCAGGTGCGAGGGGCACTG
U029XGJ200-4.seq(1>2973)                     →   GTCTTTCAAAAGATGCTGTTTTACCTGGCTAAGGTTGTGGCATGCCCACAGAGTAACCACCTCCTTTTCTTTCCCCAGATGGTGCTCCTGGCCAGGTGCGAGGGGCACTG
E02-W59500-U029XGJ200-4-4-SEQ2R.ab1(1>823)   ←   GTCTtTCAAAAGATGCTGTTT
E03-W59500-U029XGJ200-4-4-SEQ3F.ab1(1>825)   →   GTCTTTCAAAAGATGCTGTTTTACCTGGcTAAGGTTGTGGCATGCCCACAGAGTAACCACCTCCTTTTCTTTCCCCAGATGGTGCTCCTGGCCAGGTGCGAGGGGCACTG

                                                  1000      1010      1020      1030      1040      1050      1060      1070      1080      1090      1100
                                                 CAGCCAGGCGTCACGCTCCGAGCCTTTGGTGTCGTTCAGCACTGTCCTCAAGCAACCCTTCCGTTCCTCCTGTCACTGCTGCCGGCCCCAGACTTCCAAGCTGAAGGCAC
U029XGJ200-4.seq(1>2973)                     →   CAGCCAGGCGTCACGCTCCGAGCCTTTGGTGTCGTTCAGCACTGTCCTCAAGCAACCCTTCCGTTCCTCCTGTCACTGCTGCCGGCCCCAGACTTCCAAGCTGAAGGCAC
E03-W59500-U029XGJ200-4-4-SEQ3F.ab1(1>825)   →   CAGCCAGGCGTCACGCTCCGAGCCTTTGGTGTCGTTCAGCACTGTCCTCAAGCAACCCTTCCGTTCCTCCTGTCACTGCTGCCGGCCCCAGACTTCCAAGCTGAAGGCAC

                                                  1110      1120      1130      1140      1150      1160      1170      1180      1190      1200      1210
                                                 TGCGGCTGCGATGCTCAGGGGGCATGCGACTCACTGCCACCTACCGGTACATCCTCTCCTGTCACTGCGAGGAATGCAATTCCGGTAAGCCTATCCCTAACCCTCTCCTC
U029XGJ200-4.seq(1>2973)                     →   TGCGGCTGCGATGCTCAGGGGGCATGCGACTCACTGCCACCTACCGGTACATCCTCTCCTGTCACTGCGAGGAATGCAATTCCGGTAAGCCTATCCCTAACCCTCTCCTC
E03-W59500-U029XGJ200-4-4-SEQ3F.ab1(1>825)   →   TGCGGCTGCGATGCTCAGGGGGCATGCGACTCACTGCCACCTACCGGTACATCCTCTCCTGTCACTGCGAGGAATGCAATTCCGGTAAGCCTATCCCTAACCCTCTCCTC

                                                  1220      1230      1240      1250      1260      1270      1280      1290      1300      1310      1320
                                                 GGTCTCGATTCTACGGGATCCGGAGCTACTAATTTCTCCTTGCTTAAGCAAGCTGGTGATGTTGAAGAAAATCCTGGTCCTATGGTGAGCAAGGGCGAGGAGCTGTTCAC
U029XGJ200-4.seq(1>2973)                     →   GGTCTCGATTCTACGGGATCCGGAGCTACTAATTTCTCCTTGCTTAAGCAAGCTGGTGATGTTGAAGAAAATCCTGGTCCTATGGTGAGCAAGGGCGAGGAGCTGTTCAC
E03-W59500-U029XGJ200-4-4-SEQ3F.ab1(1>825)   →   GGTCTCGATTCTACGGGATCCGGAGCTACTAATTTCTCCTTGCTTAAGCAAGCTGGTGATGTTGAAGAAAATCCTGGTCCTATGGTGAGCAAGGGCGAGGAGCTGTTCAC

                                                  1330      1340      1350      1360      1370      1380      1390      1400      1410      1420      1430
                                                 CGGGGTGGTGCCCATCCTGGTCGAGCTGGACGGCGACGTAAACGGCCACAAGTTCAGCGTGTCCGGCGAGGGCGAGGGCGATGCCACCTACGGCAAGCTGACCCTGAAGT
U029XGJ200-4.seq(1>2973)                     →   CGGGGTGGTGCCCATCCTGGTCGAGCTGGACGGCGACGTAAACGGCCACAAGTTCAGCGTGTCCGGCGAGGGCGAGGGCGATGCCACCTACGGCAAGCTGACCCTGAAGT
E03-W59500-U029XGJ200-4-4-SEQ3F.ab1(1>825)   →   CGGGGTGGTGCCCATCCTGGTCGAGCTGGACGGCGACGTAAACGGCCACAAGTTCAGCGTGTCCGGCGAGGGCGAGGGCGATGCCACCTACGGCAAGCTGACCCTGAAGT

                                                  1440      1450      1460      1470      1480      1490      1500      1510      1520      1530      1540
                                                 TCATCTGCACCACCGGCAAGCTGCCCGTGCCCTGGCCCACCCTCGTGACCACCCTGACCTACGGCGTGCAGTGCTTCAGCCGCTACCCCGACCACATGAAGCAGCACGAC
U029XGJ200-4.seq(1>2973)                     →   TCATCTGCACCACCGGCAAGCTGCCCGTGCCCTGGCCCACCCTCGTGACCACCCTGACCTACGGCGTGCAGTGCTTCAGCCGCTACCCCGACCACATGAAGCAGCACGAC
E03-W59500-U029XGJ200-4-4-SEQ3F.ab1(1>825)   →   TCATCTGCACCACCGGCAAGCTGCCCGTGCCCTGGCCCACCCTCGTGACCACCCTGACCTACGGCGTGCAGTGCTTCAGCCGCTACCCCGACCACATGAAGCAGCACGAC
E05-W59500-U029XGJ200-4-4-SEQ4F.ab1(1>833)   →                                                  ACCACCCTGACCTACGGCGTGCAGTGCTTCAGCCGCTACCCCGACCACATGAAGcAGCACGAC

                                                  1550      1560      1570      1580      1590      1600      1610      1620      1630      1640      1650
                                                 TTCTTCAAGTCCGCCATGCCCGAAGGCTACGTCCAGGAGCGCACCATCTTCTTCAAGGACGACGGCAACTACAAGACCCGCGCCGAGGTGAAGTTCGAGGGCGACACCCT
U029XGJ200-4.seq(1>2973)                     →   TTCTTCAAGTCCGCCATGCCCGAAGGCTACGTCCAGGAGCGCACCATCTTCTTCAAGGACGACGGCAACTACAAGACCCGCGCCGAGGTGAAGTTCGAGGGCGACACCCT
E03-W59500-U029XGJ200-4-4-SEQ3F.ab1(1>825)   →   TTCTTCAAGTCCGCCATGCCCGAAGGCTACGTCCAGGAGCGCACCATCTTCTTCAAGGACGACGGCAACTACAAGACCCGCGCCGAGGTGAAGTTCGAGGGCGACACCCT
E05-W59500-U029XGJ200-4-4-SEQ4F.ab1(1>833)   →   TTCTTCAAGTCCGCCATGCCCGAAGGCTACGTCCAGGAGCGCACCATCTTCTTCAAGGACGACGGCAACTACAAGACCCGCGCCGAGGTGAAGTTCGAGGGCGACACCCT
```

```
                                                  1660      1670      1680      1690      1700      1710      1720      1730      1740      1750      1760
                                                  GGTGAACCGCATCGAGCTGAAGGGCATCGACTTCAAGGAGGACGGCAACATCCTGGGGCACAAGCTGGAGTACAACTACAACAGCCACAACGTCTATATCATGGCCGACA
U029XGJ200-4.seq(1>2973)                       →  GGTGAACCGCATCGAGCTGAAGGGCATCGACTTCAAGGAGGACGGCAACATCCTGGGGCACAAGCTGGAGTACAACTACAACAGCCACAACGTCTATATCATGGCCGACA
E03-W59500-U029XGJ200-4-4-SEQ3F.ab1(1>825)     →  GGTGAACCGCATCGAGCTGAAGGGCATCGA
E05-W59500-U029XGJ200-4-4-SEQ4F.ab1(1>833)     →  GGTGAACCGCATCGAGCTGAAGGGCATCGACTTCAAGGAGGACGGCAACATCCTGGGGCACAAGCTGGAGTACAACTACAACAGCCACAACGTCTATATCATGGCCGACA

                                                  1770      1780      1790      1800      1810      1820      1830      1840      1850      1860      1870
                                                  AGCAGAAGAACGGCATCAAGGTGAACTTCAAGATCCGCCACAACATCGAGGACGGCAGCGTGCAGCTCGCCGACCACTACCAGCAGAACACCCCCATCGGCGACGGCCCC
U029XGJ200-4.seq(1>2973)                       →  AGCAGAAGAACGGCATCAAGGTGAACTTCAAGATCCGCCACAACATCGAGGACGGCAGCGTGCAGCTCGCCGACCACTACCAGCAGAACACCCCCATCGGCGACGGCCCC
E05-W59500-U029XGJ200-4-4-SEQ4F.ab1(1>833)     →  AGCAGAAGAACGGCATCAAGGTGAACTTCAAGATCCGCCACAACATCGAGGACGGCAGCGTGCAGCTCGCCGACCACTACCAGCAGAACACCCCCATCGGCGACGGCCCC

                                                  1880      1890      1900      1910      1920      1930      1940      1950      1960      1970      1980
                                                  GTGCTGCTGCCCGACAACCACTACCTGAGCACCCAGTCCGCCCTGAGCAAAGACCCCAACGAGAAGCGCGATCACATGGTCCTGCTGGAGTTCGTGACCGCCGCCGGGAT
U029XGJ200-4.seq(1>2973)                       →  GTGCTGCTGCCCGACAACCACTACCTGAGCACCCAGTCCGCCCTGAGCAAAGACCCCAACGAGAAGCGCGATCACATGGTCCTGCTGGAGTTCGTGACCGCCGCCGGGAT
E05-W59500-U029XGJ200-4-4-SEQ4F.ab1(1>833)     →  GTGCTGCTGCCCGACAACCACTACCTGAGCACCCAGTCCGCCCTGAGCAAAGACCCCAACGAGAAGCGCGATCACATGGTCCTGCTGGAGTTCGTGACCGCCGCCGGGAT

                                                  1990      2000      2010      2020      2030      2040      2050      2060      2070      2080      2090
                                                  CACTCTCGGCATGGACGAGCTGTACAAGTGAGGCCCGCTGCTGTGTGTGGCTTCTGGATGGGACAACTGTAGAGGCAGTTCGACCAGCCAGTGAAAGACTGGCAAGAAAA
U029XGJ200-4.seq(1>2973)                       →  CACTCTCGGCATGGACGAGCTGTACAAGTGAGGCCCGCTGCTGTGTGTGGCTTCTGGATGGGACAACTGTAGAGGCAGTTCGACCAGCCAGTGAAAGACTGGCAAGAAAA
E05-W59500-U029XGJ200-4-4-SEQ4F.ab1(1>833)     →  CACTCTCGGCATGGACGAGCTGTACAAGTGAGGCCCGCTGCTGTGTGTGGCTTCTGGATGGGACAACTGTAGAGGCAGTTCGACCAGCCAGTGAAAGACTGGCAAGAAAA

                                                  2100      2110      2120      2130      2140      2150      2160      2170      2180      2190      2200
                                                  GAGTTAAGGCAAAAAAGGATGCAACAATTCTCCCGGGACTCTGCATATTCTAGTAATAAAGACTCTACATGCTTGTTGACAGAGAGAGATACTCTGGGAACTTCTTTGCA
U029XGJ200-4.seq(1>2973)                       →  GAGTTAAGGCAAAAAAGGATGCAACAATTCTCCCGGGACTCTGCATATTCTAGTAATAAAGACTCTACATGCTTGTTGACAGAGAGAGATACTCTGGGAACTTCTTTGCA
E05-W59500-U029XGJ200-4-4-SEQ4F.ab1(1>833)     →  GAGTTAAGGCAAAAAAGGATGCAACAATTCTCCCGGGACTCTGCATATTCTAGTAATAAAGACTCTACATGCTTGTTGACAGAGAGAGATACTCTGGGAACTTCTTTGCA
E06-W59500-U029XGJ200-4-4-SEQ5F.ab1(1>833)     →                               CTCCCGGGACTCTGCATATTCTAGTAATAAAGACTCTACATGCTTGTTGACAGAGAGAGATACTCTGGGAACTTCTTTGCA

                                                  2210      2220      2230      2240      2250      2260      2270      2280      2290      2300      2310
                                                  GTTCCCATCTCCTTTCTCTGGTACAATTTCTTTTGGTTCATTTTCAGATTCAGGCATTTTCCCCCTTGGCTCTCAATGCTGTTTGGGTTTCCAACAATTCAGCATTAGTG
U029XGJ200-4.seq(1>2973)                       →  GTTCCCATCTCCTTTCTCTGGTACAATTTCTTTTGGTTCATTTTCAGATTCAGGCATTTTCCCCCTTGGCTCTCAATGCTGTTTGGGTTTCCAACAATTCAGCATTAGTG
E05-W59500-U029XGJ200-4-4-SEQ4F.ab1(1>833)     →  GTTCCCATCTCCTTTCTCTGGTACAATTTCTTTTGGTTCATTTTCAGATTCAGGCATTTTCCCCCTTGGCTCTCAATGCTGTTTGGGTTTCCAACAATTCAGCATTAGTG
E06-W59500-U029XGJ200-4-4-SEQ5F.ab1(1>833)     →  GTTCCCATCTCCTTTCTCTGGTACAATTTCTTTTGGTTCATTTTCAGATTCAGGCATTTTCCCCCTTGGCTCTCAATGCTGTTTGGGTTTCCAACAATTCAGCATTAGTG

                                                  2320      2330      2340      2350      2360      2370      2380      2390      2400      2410      2420
                                                  GGAAAAAGTGGGCCCTCATACACAAGCGTGTCAGGCTGTCAGTGTTTGGTGCACGCTGGGGAAGAATTTACTTTGGAAAGTAGAAAAGCCCAGCTTTTCCTGGGACATCT
U029XGJ200-4.seq(1>2973)                       →  GGAAAAAGTGGGCCCTCATACACAAGCGTGTCAGGCTGTCAGTGTTTGGTGCACGCTGGGGAAGAATTTACTTTGGAAAGTAGAAAAGCCCAGCTTTTCCTGGGACATCT
E06-W59500-U029XGJ200-4-4-SEQ5F.ab1(1>833)     →  GGAAAAAGTGGGCCCTCATACACAAGCGTGTCAGGCTGTCAGTGTTTGGTGCACGCTGGGGAAGAATTTACTTTGGAAAGTAGAAAAGCCCAGCTTTTCCTGGGACATCT
E04-W59500-U029XGJ200-4-Regen-seqR.ab1(1>833)  ←                            CGTGTCAGGCTGTCAGTGTTTGGTGCACGCTGGGGAAGAATTTACTTTGGAAAGTAGAAAAGCCCAGCTTTTCCTGGGACATCT

                                                  2430      2440      2450      2460      2470      2480      2490      2500      2510      2520      2530
                                                  TCTGTTATTGTTGATGTTTTTTTTTACCTTGTCATTTTGGTCTAAGGTTGCCATTGCTGCTAAAGGTTACCGATTTCAAAGTCCAGATACCAAGCATGTGGATATGTTTA
U029XGJ200-4.seq(1>2973)                       →  TCTGTTATTGTTGATGTTTTTTTTTACCTTGTCATTTTGGTCTAAGGTTGCCATTGCTGCTAAAGGTTACCGATTTCAAAGTCCAGATACCAAGCATGTGGATATGTTTA
E06-W59500-U029XGJ200-4-4-SEQ5F.ab1(1>833)     →  TCTGTTATTGTTGATGTTTTTTTTTACCTTGTCATTTTGGTCTAAGGTTGCCATTGCTGCTAAAGGTTACCGATTTCAAAGTCCAGATACCAAGCATGTGGATATGTTTA
E04-W59500-U029XGJ200-4-Regen-seqR.ab1(1>833)  ←  TCTGTTATTGTTGATGTTTTTTTTTACCTTGTCATTTTGGTCTAAGGTTGCCATTGCTGCTAAAGGTTACCGATTTCAAAGTCCAGATACCAAGCATGTGGATATGTTTA
```

```
                                                   2540      2550      2560      2570      2580      2590      2600      2610      2620      2630      2640
                                               GCTACGTTTACTCACAGCCAGCGAACTGACATTAAAATAACTAACAAACAGATTCTTTTATGTGATGCTGGAACTCTTGACAGCTATAATTATTATTCAGAAATGACTTT
U029XGJ200-4.seq(1>2973)                    →  GCTACGTTTACTCACAGCCAGCGAACTGACATTAAAATAACTAACAAACAGATTCTTTTATGTGATGCTGGAACTCTTGACAGCTATAATTATTATTCAGAAATGACTTT
E06-W59500-U029XGJ200-4-4-SEQ5F.ab1(1>833)  →  GCTACGTTTACTCACAGCCAGCGAACTGACATTAAAATAACTAACAAACAGATTCTTTTATGTGATGCTGGAACTCTTGACAGCTATAATTATTATTCAGAAATGACTTT
E04-W59500-U029XGJ200-4-Regen-seqR.ab1(1>833) ← GCTACGTTTACTCACAGCCAGCGAACTGACATTAAAATAACTAACAAACAGATTCTTTTATGTGATGCTGGAACTCTTGACAGCTATAATTATTATTCAGAAATGACTTT

                                                   2650      2660      2670      2680      2690      2700      2710      2720      2730      2740      2750
                                               TTGAAAGTAAAAGCAGCATAAAGAATTTGTCACAGGAAGGCTGTCTCAGATAAATTATGGTAAAATTTTGTAAGGGAGCAGACTTTTAAAGACTTGCACAAATACGGATC
U029XGJ200-4.seq(1>2973)                    →  TTGAAAGTAAAAGCAGCATAAAGAATTTGTCACAGGAAGGCTGTCTCAGATAAATTATGGTAAAATTTTGTAAGGGAGCAGACTTTTAAAGACTTGCACAAATACGGATC
E06-W59500-U029XGJ200-4-4-SEQ5F.ab1(1>833)  →  TTGAAAGTAAAAGCAGCATAAAGAATTTGTCACAGGAAGGCTGTCTCAGATAAATTATGGTAAAATTTTGTAAGGGAGCAGACTTTTAAAGACTTGCACAAATACGGATC
E04-W59500-U029XGJ200-4-Regen-seqR.ab1(1>833) ← TTGAAAGTAAAAGCAGCATAAAGAATTTGTCACAGGAAGGCTGTCTCAGATAAATTATGGTAAAATTTTGTAAGGGAGCAGACTTTTAAAGACTTGCACAAATACGGATC

                                                   2760      2770      2780      2790      2800      2810      2820      2830      2840      2850      2860
                                               CTGCACTGACTCTGGAAAAGGCATATATGTACTAGTGGCATGGAGAATGCACCATACTCATGCATGCAAATTAGACAACCAAGTATGAATCTATTTGTGGGTGTGCTATA
U029XGJ200-4.seq(1>2973)                    →  CTGCACTGACTCTGGAAAAGGCATATATGTACTAGTGGCATGGAGAATGCACCATACTCATGCATGCAAATTAGACAACCAAGTATGAATCTATTTGTGGGTGTGCTATA
E06-W59500-U029XGJ200-4-4-SEQ5F.ab1(1>833)  →  CTGCACTGACTCTGGAAAAGGCATATATGTACTAGTGGCATGGAGAATGCACCATACTCATGCATGCAAATTAGACAACCAAGTATGAATCTATTTGTGGGTGTGCTATA
E04-W59500-U029XGJ200-4-Regen-seqR.ab1(1>833) ← CTGCACTGACTCTGGAAAAGGCATATATGTACTAGTGGCATGGAGAATGCACCATACTCATGCATGCAAATTAGACAACCAAGTATGAATCTATTTGTGGGTGTGCTATA

                                                   2870      2880      2890      2900      2910      2920      2930      2940      2950      2960      2970
                                               GCTTTAGCCGTGTCACGGGCATCATTCTCTAATATCCACTTGTCCATGTGAAACATGTTGCCAAAATGGTGGCCTGGCTTGTCTTCTGAACGTTTGGTTCAAATGTGTTT
U029XGJ200-4.seq(1>2973)                    →  GCTTTAGCCGTGTCACGGGCATCATTCTCTAATATCCACTTGTCCATGTGAAACATGTTGCCAAAATGGTGGCCTGGCTTGTCTTCTGAACGTTTGGTTCAAATGTGTTT
E06-W59500-U029XGJ200-4-4-SEQ5F.ab1(1>833)  →  GCTTTAGCCGTGTCACGGGCATCATTCTCTAATATCCACTTGTCCATGTGAAACATGTTGCCAAAATGGTGGCCTGGCTTGTCTTCTGAACG
E04-W59500-U029XGJ200-4-Regen-seqR.ab1(1>833) ← GCTTTAGCCGTGTCACGGGCATCATTCTCTAATATCCACTTGTCCATGTGAAACATGTTGCCAAAATGGTGGCCTGGCTTGTCTTCTGAACGTTTGGTTCAAATGTGTTT

                                                   2980      2990      3000      3010      3020      3030      3040      3050      3060      3070      3080
                                               TGGTCCTGGAGGCTCAAATTTTGAGTTATTCCCACGTTTTGAAATAAAAAGAGTATATTCAAAAATCATTGGGACGTCAGGTGGCACTTTTCGGGGAAATGTGCGCGGAA
U029XGJ200-4.seq(1>2973)                    →  TGGTCCTGGAGGCTCAAATTTTGAGTTATTCCCACGTTTTGAAATAAAAAGAGTATATTCAAAA
E04-W59500-U029XGJ200-4-Regen-seqR.ab1(1>833) ← TGGTCCTGGAGGCTCAAATTTTGAGTTATTCCCACGTTTTGAAATAAAAAGAGTATATTCAAAAATCATTGGGACGTCAGGTGGCACTTTTCGGGGAAATGTGCGCGGAA

                                                   3090      3100      3110      3120      3130      3140      3150      3160
                                               CCCCTATTTGTTTATTTTTCTAAATACATTCAAATATGTATCCGCTCATGAGACAATAACCCTGATAAATGCTTCAATAATATTGAAAA
E04-W59500-U029XGJ200-4-Regen-seqR.ab1(1>833) ← CCCCTATTTGTTTATTTTTCTAAATACATTCAAATATGTATCCGCTCATGAGACAATAACCCTGATAAATGCTTCAATAATAtTGAAAA
```
