## Supplementary material for "Putative Role of Norrin in Neuroretinal Differentiation Revealed by bulk and scRNA Sequencing of Human Retinal Organoids": Data02_NDP-KO-GFP_repair-template.pdf

```
                                             10        20        30        40        50        60        70        80        90       100       110
                                    TTCCTGGCCTTTTGCTGGCCTTTTGCTCACATGTTCTTTCCGATGCGTGTGATACCTTAACCATGCTGGAGGCCAGTTAGCATAATGTGTATTTCAGATGCAGTTTGTCA
F01-V72508-U029XGJ200-2-1.8K-SEQR.ab1(1>806)  →  TTCCTGGCCTTTTGCTGGCCTTTTGCTCACATGTTCTTTCCGATGCGTGTGATACCTTAACCATGCTGGAGGCCAGTTAGCATAATGTGTATTTCAGATGCAGTTTGTCA
U029XGJ200-2.seq(1>2594)                      →                                              GCGTGTGATACCTTAACCATGCTGGAGGCCAGTTAGCATAATGTGTATTTCAGATGCAGTTTGTCA

                                            120       130       140       150       160       170       180       190       200       210       220
                                    TCAGTGCTGGGTGCTATGGGGCTTAAAACAAAAACAGGATGAGCCTAGAGCACAGAAGTGGAAAATTAGGTGGGTGGGTGAGTATGGACTTGGAAATGGAAAATGTGCTT
F01-V72508-U029XGJ200-2-1.8K-SEQR.ab1(1>806)  →  TCAGTGCTGGGTGCTATGGGGCTTAAAACAAAAACAGGATGAGCCTAGAGCACAGAAGTGGAAAATTAGGTGGGTGGGTGAGTATGGACTTGGAAATGGAAAATGTGCTT
U029XGJ200-2.seq(1>2594)                      →  TCAGTGCTGGGTGCTATGGGGCTTAAAACAAAAACAGGATGAGCCTAGAGCACAGAAGTGGAAAATTAGGTGGGTGGGTGAGTATGGACTTGGAAATGGAAAATGTGCTT

                                            230       240       250       260       270       280       290       300       310       320       330
                                    CCCTGTGGGACATGGCTTGTGTTCATTAGTGCTGACCTGCCTCCTCTCTAGCCCTGTAAGCTGACGCTGACACCAAAGCTTGCTGAGAAACTGAATAGACCCTCCCGCCC
F01-V72508-U029XGJ200-2-1.8K-SEQR.ab1(1>806)  →  CCCTGTGGGACATGGCTTGTGTTCATTAGTGCTGACCTGCCTCCTCTCTAGCCCTGTAAGCTGACGCTGACACCAAAGCTTGCTGAGAAACTGAATAGACCCTCCCGCCC
U029XGJ200-2.seq(1>2594)                      →  CCCTGTGGGACATGGCTTGTGTTCATTAGTGCTGACCTGCCTCCTCTCTAGCCCTGTAAGCTGACGCTGACACCAAAGCTTGCTGAGAAACTGAATAGACCCTCCCGCCC

                                            340       350       360       370       380       390       400       410       420       430       440
                                    CGACCAACCAAGGACAAAGATGAGACCACCCAATTCGGTTACGTTGTTGCCAGAACAACATGTTTAATCTTTAACATGGGTTCAAACTATTCTTGGCCCTAGGAACATGG
F01-V72508-U029XGJ200-2-1.8K-SEQR.ab1(1>806)  →  CGACCAACCAAGGACAAAGATGAGACCACCCAATTCGGTTACGTTGTTGCCAGAACAACATGTTTAATCTTTAACATGGGTTCAAACTATTCTTGGCCCTAGGAACATGG
U029XGJ200-2.seq(1>2594)                      →  CGACCAACCAAGGACAAAGATGAGACCACCCAATTCGGTTACGTTGTTGCCAGAACAACATGTTTAATCTTTAACATGGGTTCAAACTATTCTTGGCCCTAGGAACATGG

                                            450       460       470       480       490       500       510       520       530       540       550
                                    ACTTCAGCAATTAAAGTCAACATGTGCTTCCATTAAACCATTGTGTCCACCTCCAAATGGTTATAAATATGTAGCATAAGCTATGGGAGTTGGGGTGGAATGGATGACAG
F01-V72508-U029XGJ200-2-1.8K-SEQR.ab1(1>806)  →  ACTTCAGCAATTAAAGTCAACATGTGCTTCCATTAAACCATTGTGTCCACCTCCAAATGGTTATAAATATGTAGCATAAGCTATGGGAGTTGGGGTGGAATGGATGACAG
U029XGJ200-2.seq(1>2594)                      →  ACTTCAGCAATTAAAGTCAACATGTGCTTCCATTAAACCATTGTGTCCACCTCCAAATGGTTATAAATATGTAGCATAAGCTATGGGAGTTGGGGTGGAATGGATGACAG

                                            560       570       580       590       600       610       620       630       640       650       660
                                    CCTTTGCTAATGACGCTCTAGAAACCAATATTCTCCTCTCAAAATAACATGGAAAAATTCTACTTAATATCACCTGGGTTCCATTAGTGGTTCTGGGTAAATAATTCTGG
F01-V72508-U029XGJ200-2-1.8K-SEQR.ab1(1>806)  →  CCTTTGCTAATGACGCTCTAGAAACCAATATTCTCCTCTCAAAATAACATGGAAAAATTCTACTTAATATCACCTGGGTTCCATTAGTGGTTCTGGGTAAATAATTCTGG
U029XGJ200-2.seq(1>2594)                      →  CCTTTGCTAATGACGCTCTAGAAACCAATATTCTCCTCTCAAAATAACATGGAAAAATTCTACTTAATATCACCTGGGTTCCATTAGTGGTTCTGGGTAAATAATTCTGG

                                            670       680       690       700       710       720       730       740       750       760       770
                                    GGAAAGTAATTTCTGTTTTCATTCCAGCTGTGCAGCAGATACTGTGATGATGGATTGCAAGTGCAAAGAGTAAGACAAAACTCCAGCACATAAAGGACAATGACAACCAG
F01-V72508-U029XGJ200-2-1.8K-SEQR.ab1(1>806)  →  GGAAAGTAATTTCTGTTTTCATTCCAGCTGTGCAGCAGATACTGTGATGATGGATTGCAAGTGCAAAGAGTAAGACAAAACTCCAGCACATAAAGGACAATGACAACCAG
U029XGJ200-2.seq(1>2594)                      →  GGAAAGTAATTTCTGTTTTCATTCCAGCTGTGCAGCAGATACTGTGATGATGGATTGCAAGTGCAAAGAGTAAGACAAAACTCCAGCACATAAAGGACAATGACAACCAG
A11-V72508-U029XGJ200-2-2-SEQ1F.ab1(1>828)    →                         CCAGCTGTGCAGCAGATACTGTGATGATGGATTGCAAGTGCAAAGAGTAAGACAAAACTCCAGCACATAAAGGACAATGACAACCAG

                                            780       790       800       810       820       830       840       850       860       870       880
                                    AAAGCTTCAGCCCGATCCTGCCCTTTCCTTGAACGGGACTGGATCCTAGGAGGTGAAGCCATTTCCAATTTTTTGTCCTCTGCCTCCCTCTGCTGTTCTTCTAGAGAAGT
F01-V72508-U029XGJ200-2-1.8K-SEQR.ab1(1>806)  →  AAAGCTTCAGCCCGATCCTGCCCTTTCCTTGAACGG
U029XGJ200-2.seq(1>2594)                      →  AAAGCTTCAGCCCGATCCTGCCCTTTCCTTGAACGGGACTGGATCCTAGGAGGTGAAGCCATTTCCAATTTTTTGTCCTCTGCCTCCCTCTGCTGTTCTTCTAGAGAAGT
A11-V72508-U029XGJ200-2-2-SEQ1F.ab1(1>828)    →  AAAGCTTCAGCCCGATCCTGCCCTTTCCTTGAACGGGACTGGATCCTAGGAGGTGAAGCCATTTCCAATTTTTTGTCCTCTGCCTCCCTCTGCTGTTCTTCTAGAGAAGT
```

```
                                           890       900       910       920       930       940       950       960       970       980       990
                                           TTTTCCTTACAACAATGGTGAGCAAGGGCGAGGAGCTGTTCACCGGGGTGGTGCCCATCCTGGTCGAGCTGGACGGCGACGTAAACGGCCACAAGTTCAGCGTGTCCGGC
U029XGJ200-2.seq(1>2594)                 → TTTTCCTTACAACAATGGTGAGCAAGGGCGAGGAGCTGTTCACCGGGGTGGTGCCCATCCTGGTCGAGCTGGACGGCGACGTAAACGGCCACAAGTTCAGCGTGTCCGGC
A11-V72508-U029XGJ200-2-2-SEQ1F.ab1(1>828) → TTTTCCTTACAACAATGGTGAGCAAGGGCGAGGAGCTGTTCACCGGGGTGGTGCCCATCCTGGTCGAGCTGGACGGCGACGTAAACGGCCACAAGTTCAGCGTGTCCGGC

                                           1000      1010      1020      1030      1040      1050      1060      1070      1080      1090      1100
                                           GAGGGCGAGGGCGATGCCACCTACGGCAAGCTGACCCTGAAGTTCATCTGCACCACCGGCAAGCTGCCCGTGCCCTGGCCCACCCTCGTGACCACCCTGACCTACGGCGT
U029XGJ200-2.seq(1>2594)                 → GAGGGCGAGGGCGATGCCACCTACGGCAAGCTGACCCTGAAGTTCATCTGCACCACCGGCAAGCTGCCCGTGCCCTGGCCCACCCTCGTGACCACCCTGACCTACGGCGT
A11-V72508-U029XGJ200-2-2-SEQ1F.ab1(1>828) → GAGGGCGAGGGCGATGCCACCTACGGCAAGCTGACCCTGAAGTTCATCTGCACCACCGGCAAGCTGCCCGTGCCCTGGCCCACCCTCGTGACCACCCTGACCTACGGCGT

                                           1110      1120      1130      1140      1150      1160      1170      1180      1190      1200      1210
                                           GCAGTGCTTCAGCCGCTACCCCGACCACATGAAGCAGCACGACTTCTTCAAGTCCGCCATGCCCGAAGGCTACGTCCAGGAGCGCACCATCTTCTTCAAGGACGACGGCA
U029XGJ200-2.seq(1>2594)                 → GCAGTGCTTCAGCCGCTACCCCGACCACATGAAGCAGCACGACTTCTTCAAGTCCGCCATGCCCGAAGGCTACGTCCAGGAGCGCACCATCTTCTTCAAGGACGACGGCA
A11-V72508-U029XGJ200-2-2-SEQ1F.ab1(1>828) → GCAGTGCTTCAGCCGCTACCCCGACCACATGAAGCAGCACGACTTCTTCAAGTCCGCCATGCCCGAAGGCTACGTCCAGGAGCGCACCATCTTCTTCAAGGACGACGGCA

                                           1220      1230      1240      1250      1260      1270      1280      1290      1300      1310      1320
                                           ACTACAAGACCCGCGCCGAGGTGAAGTTCGAGGGCGACACCCTGGTGAACCGCATCGAGCTGAAGGGCATCGACTTCAAGGAGGACGGCAACATCCTGGGGCACAAGCTG
U029XGJ200-2.seq(1>2594)                 → ACTACAAGACCCGCGCCGAGGTGAAGTTCGAGGGCGACACCCTGGTGAACCGCATCGAGCTGAAGGGCATCGACTTCAAGGAGGACGGCAACATCCTGGGGCACAAGCTG
A11-V72508-U029XGJ200-2-2-SEQ1F.ab1(1>828) → ACTACAAGACCCGCGCCGAGGTGAAGTTCGAGGGCGACACCCTGGTGAACCGCATCGAGCTGAAGGGCATCGACTTCAAGGAGGACGGCAACATCCTGGGGCACAAGCTG

                                           1330      1340      1350      1360      1370      1380      1390      1400      1410      1420      1430
                                           GAGTACAACTACAACAGCCACAACGTCTATATCATGGCCGACAAGCAGAAGAACGGCATCAAGGTGAACTTCAAGATCCGCCACAACATCGAGGACGGCAGCGTGCAGCT
U029XGJ200-2.seq(1>2594)                 → GAGTACAACTACAACAGCCACAACGTCTATATCATGGCCGACAAGCAGAAGAACGGCATCAAGGTGAACTTCAAGATCCGCCACAACATCGAGGACGGCAGCGTGCAGCT
A11-V72508-U029XGJ200-2-2-SEQ1F.ab1(1>828) → GAGTACAACTACAACAGCCACAACGTCTATATCATGGCCGACAAGCAGAAGAACGGCATCAAGGTGAACTTCAAGATCCGCCACAACATCGAGGACGGCAGCGTGCAGCT
H04-V72508-U029XGJ200-2-2-SEQ2F.ab1(1>845) →                                                            CAAGGTGAACTTCAAGATCCGCCACAACATCGAGGACGGCAGCGTGCAGCT

                                           1440      1450      1460      1470      1480      1490      1500      1510      1520      1530      1540
                                           CGCCGACCACTACCAGCAGAACACCCCCATCGGCGACGGCCCCGTGCTGCTGCCCGACAACCACTACCTGAGCACCCAGTCCGCCCTGAGCAAAGACCCCAACGAGAAGC
U029XGJ200-2.seq(1>2594)                 → CGCCGACCACTACCAGCAGAACACCCCCATCGGCGACGGCCCCGTGCTGCTGCCCGACAACCACTACCTGAGCACCCAGTCCGCCCTGAGCAAAGACCCCAACGAGAAGC
A11-V72508-U029XGJ200-2-2-SEQ1F.ab1(1>828) → CGCCGACCACTACCAGCAGAACACCCCCATCGGCGACGGCCCCGTGCTGCTGCCCGACAACCACTACCTGAGCACCCAGTC
H04-V72508-U029XGJ200-2-2-SEQ2F.ab1(1>845) → CGCCGACCACTACCAGCAGAACACCCCCATCGGCGACGGCCCCGTGCTGCTGCCCGACAACCACTACCTGAGCACCCAGTCCGCCCTGAGCAAAGACCCCAACGAGAAGC

                                           1550      1560      1570      1580      1590      1600      1610      1620      1630      1640      1650
                                           GCGATCACATGGTCCTGCTGGAGTTCGTGACCGCCGCCGGGATCACTCTCGGCATGGACGAGCTGTACAAGGGATCCGGAGCTACTAATTTCTCCTTGCTTAAGCAAGCT
U029XGJ200-2.seq(1>2594)                 → GCGATCACATGGTCCTGCTGGAGTTCGTGACCGCCGCCGGGATCACTCTCGGCATGGACGAGCTGTACAAGGGATCCGGAGCTACTAATTTCTCCTTGCTTAAGCAAGCT
H04-V72508-U029XGJ200-2-2-SEQ2F.ab1(1>845) → GCGATCACATGGTCCTGCTGGAGTTCGTGACCGCCGCCGGGATCACTCTCGGCATGGACGAGCTGTACAAGGGATCCGGAGCTACTAATTTCTCCTTGCTTAAGCAAGCT

                                           1660      1670      1680      1690      1700      1710      1720      1730      1740      1750      1760
                                           GGTGATGTTGAAGAAAATCCTGGTCCTGTAAGACCAAGGTCTCTGTGAGTAGAGCATTTCCCTCAGAAACAGGCAAGAAGCCATATTTCTTTGTCAGGGGATGGAGAACT
U029XGJ200-2.seq(1>2594)                 → GGTGATGTTGAAGAAAATCCTGGTCCTGTAAGACCAAGGTCTCTGTGAGTAGAGCATTTCCCTCAGAAACAGGCAAGAAGCCATATTTCTTTGTCAGGGGATGGAGAACT
H04-V72508-U029XGJ200-2-2-SEQ2F.ab1(1>845) → GGTGATGTTGAAGAAAATCCTGGTCCTGTAAGACCAAGGTCTCTGTGAGTAGAGCATTTCCCTCAGAAACAGGCAAGAAGCCATATTTCTTTGTCAGGGGATGGAGAACT
```

```
                                           1770      1780      1790      1800      1810      1820      1830      1840      1850      1860      1870
                                      AAGATCATAGCCCAAACTTAAGTGATGTGGCTGGCCCACAGGGCTTGTGGGAGAATGAGGCTTTTAGAATTTCCCCCCGTGCTTTGCACTGTTATCAGTAAAAGTGCCTA
U029XGJ200-2.seq(1>2594)            → AAGATCATAGCCCAAACTTAAGTGATGTGGCTGGCCCACAGGGCTTGTGGGAGAATGAGGCTTTTAGAATTTCCCCCCGTGCTTTGCACTGTTATCAGTAAAAGTGCCTA
H04-V72508-U029XGJ200-2-2-SEQ2F.ab1(1>845) → AAGATCATAGCCCAAACTTAAGTGATGTGGCTGGCCCACAGGGCTTGTGGGAGAATGAGGCTTTTAGAATTTCCCCCCGTGCTTTGCACTGTTATCAGTAAAAGTGCCTA

                                           1880      1890      1900      1910      1920      1930      1940      1950      1960      1970      1980
                                      ATCAATCATGTGCACACCCATCCAGTGGGACAGGTGTCCTTTGTCCCTGGCTGTGATGTTTATGAGTGGTAGTGTGCTTTTCCAAAAAAGAAAAATGGAGTTTACAAGTC
U029XGJ200-2.seq(1>2594)            → ATCAATCATGTGCACACCCATCCAGTGGGACAGGTGTCCTTTGTCCCTGGCTGTGATGTTTATGAGTGGTAGTGTGCTTTTCCAAAAAAGAAAAATGGAGTTTACAAGTC
H04-V72508-U029XGJ200-2-2-SEQ2F.ab1(1>845) → ATCAATCATGTGCACACCCATCCAGTGGGACAGGTGTCCTTTGTCCCTGGCTGTGATGTTTATGAGTGGTAGTGTGCTTTTCCAAAAAAGAAAAATGGAGTTTACAAGTC

                                           1990      2000      2010      2020      2030      2040      2050      2060      2070      2080      2090
                                      ATCATAACTCTCTAGCTCCATCATCAATAAAGAGGAAGAGAATGCAAAGAATGCGATTGAGCAGTGGAACATACTTCTTGGAGGGCTCTTTAAAGAGATTCTTGGTGAAA
U029XGJ200-2.seq(1>2594)            → ATCATAACTCTCTAGCTCCATCATCAATAAAGAGGAAGAGAATGCAAAGAATGCGATTGAGCAGTGGAACATACTTCTTGGAGGGCTCTTTAAAGAGATTCTTGGTGAAA
H04-V72508-U029XGJ200-2-2-SEQ2F.ab1(1>845) → ATCATAACTCTCTAGCTCCATCATCAATAAAGAGGAAGAGAATGCAAAGAATGCGATTGAGCAGTGGAACATACTTCTTGGAGGGCTCTTTAAAGAGATTCTTGGTGAAA
A07-V72508-U029XGJ200-2-2-SEQ3F.ab1(1>840) →                                                                               TGGAGGGCTCTTTAAAGAGATTCTTGGTGAAA

                                           2100      2110      2120      2130      2140      2150      2160      2170      2180      2190      2200
                                      TAACTGGCTTTGTGGTCTGAGCCTTTCTTCAGGTGCTACATTTCTGTTCAAATGATTTCAAAAGATAAGATGGCAATATGATCATTTATCATTTTTACATTGCTTATCCA
U029XGJ200-2.seq(1>2594)            → TAACTGGCTTTGTGGTCTGAGCCTTTCTTCAGGTGCTACATTTCTGTTCAAATGATTTCAAAAGATAAGATGGCAATATGATCATTTATCATTTTTACATTGCTTATCCA
H04-V72508-U029XGJ200-2-2-SEQ2F.ab1(1>845) → TAACTGGCTTTGTGGTCTGAGCCTTTCTTCAGGTGCTACATTTCTGTTCAAATGATTTCAAAAGATAAGATGGCAATATGATCATTTATCATTTTTACATTGCTTATCCA
A07-V72508-U029XGJ200-2-2-SEQ3F.ab1(1>840) → TAACTGGCTTTGTGGTCTGAGCCTTTCTTCAGGTGCTACATTTCTGTTCAAATGATTTCAAAAGATAAGATGGCAATATGATCATTTATCATTTTTACATTGCTTATCCA

                                           2210      2220      2230      2240      2250      2260      2270      2280      2290      2300      2310
                                      AATCGTTCTCATTTGGAAAAGGATGCATGTTTCTTGCAGAGGCAGTCCGTGCCTCAGTGGTGTGCAAGGGAATATCATGCTTAAAAGGCCACCAAGAGTGGCCAGCTACA
U029XGJ200-2.seq(1>2594)            → AATCGTTCTCATTTGGAAAAGGATGCATGTTTCTTGCAGAGGCAGTCCGTGCCTCAGTGGTGTGCAAGGGAATATCATGCTTAAAAGGCCACCAAGAGTGGCCAGCTACA
H04-V72508-U029XGJ200-2-2-SEQ2F.ab1(1>845) → AATCGTTCTCATTTGGAAAAGGAT
A07-V72508-U029XGJ200-2-2-SEQ3F.ab1(1>840) → AATCGTTCTCATTTGGAAAAGGATGCATGTTTCTTGCAGAGGCAGTCCGTGCCTCAGTGGTGTGCAAGGGAATATCATGCTTAAAAGGCCACCAAGAGTGGCCAGCTACA

                                           2320      2330      2340      2350      2360      2370      2380      2390      2400      2410      2420
                                      TGGACCACCCAGAAGGGAGGCACATTGTAGGGTGGAACATACGAGATCTCACGACTGTCACTGCCCCTAAACAGGCACAATGAATGAAATTTGTCCACACTTGGTAAATA
U029XGJ200-2.seq(1>2594)            → TGGACCACCCAGAAGGGAGGCACATTGTAGGGTGGAACATACGAGATCTCACGACTGTCACTGCCCCTAAACAGGCACAATGAATGAAATTTGTCCACACTTGGTAAATA
A07-V72508-U029XGJ200-2-2-SEQ3F.ab1(1>840) → TGGACCACCCAGAAGGGAGGCACATTGTAGGGTGGAACATACGAGATCTCACGACTGTCACTGCCCCTAAACAGGCACAATGAATGAAATTTGTCCACACTTGGTAAATA

                                           2430      2440      2450      2460      2470      2480      2490      2500      2510      2520      2530
                                      ATAATTTGTTTAGCCCAACCTCAGAGATTTTTACATTTAGTTTTGAGATGAAAATCACTTAAACATTCTCGTATGGCAAGTAATTTTTGAAATTACCAGTATAATTAAAT
U029XGJ200-2.seq(1>2594)            → ATAATTTGTTTAGCCCAACCTCAGAGATTTTTACATTTAGTTTTGAGATGAAAATCACTTAAACATTCTCGTATGGCAAGTAATTTTTGAAATTACCAGTATAATTAAAT
A07-V72508-U029XGJ200-2-2-SEQ3F.ab1(1>840) → ATAATTTGTTTAGCCCAACCTCAGAGATTTTTACATTTAGTTTTGAGATGAAAATCACTTAAACATTCTCGTATGGCAAGTAATTTTTGAAATTACCAGTATAATTAAAT

                                           2540      2550      2560      2570      2580      2590      2600      2610      2620      2630      2640
                                      AATAATTTGTTTAGCCCAACCTCAGAGATTTTTACATTTAGTTTTGAGATGAAAATCACTTAAACATTCTCGTATGGCAAGTAATTTTTGAAATTACCAGTATAATTAAT
U029XGJ200-2.seq(1>2594)            → AATAATTTGTTTAGCCCAACCTCAGAGATTTTTACATTTAGTTTTGAGATGAAAATCACTTAAACATTCTCGTATGGCAAGTAATTTTTGAAATTACCAGTATAATTA
A07-V72508-U029XGJ200-2-2-SEQ3F.ab1(1>840) → AATAATTTGTTTAGCCCAACCTCAGAGATTTTTACATTTAGTTTTGAGATGAAAATCACTTAAACATTCTCGTATGGCAAGTAATTTTTGAAATTACCAGTATAATTAAT
```

```
                                              2650      2660      2670      2680      2690      2700      2710      2720      2730      2740      2750
                                          CATTGGGACGTCAGGTGGCACTTTTCGGGGAAATGTGCGCGGAACCCCTATTTGTTTATTTTTCTAAATACATTCAAATATGTATCCGCTCATGAGACAATAACCCTGAT
A07-V72508-U029XGJ200-2-2-SEQ3F.ab1(1>840)  →  CATTGGGACGTCAGGTGGCACTTTTCGGGGAAATGTGCGCGGAACCCCTATTTGTTTATTTTTCTAAATACATTCAAATATGTATCCGCTCATGAGACAATAACCCTGAT

                                              2760      2770      2780      2790      2800      2810      2820      2830      2840      2850      2860
                                          AAATGCTTCAATAATATTGAAAAAGGAAGAGTATGAGTATTCAACATTTCCGTGTCGCCCTTATTCCCTTTTTTGCGGCATTTTGCCTTCCTGTTTTTGCTCACCCAGAA
A07-V72508-U029XGJ200-2-2-SEQ3F.ab1(1>840)  →  AAATGCTTCAATAATATTGAAAAAGGAAGAGTATGAGTATTCAACATTTCCGTGTCGCCCTTATTCCCTTTTTTGCGGCATTTTGCCTTCCTGTTTTTGCTCACCCAGAA

                                              2870      2880      2890
                                          ACGCTGGTGAAAGTAAAAGATGCTGAAGATCAGTTGGG
A07-V72508-U029XGJ200-2-2-SEQ3F.ab1(1>840)  →  ACGCTGGTGAAAGTAAAAGATGCTGAAGATCAGTTGGG
```
