## Supplementary Figures and Tables for "Putative Role of Norrin in Neuroretinal Differentiation Revealed by bulk and scRNA Sequencing of Human Retinal Organoids"

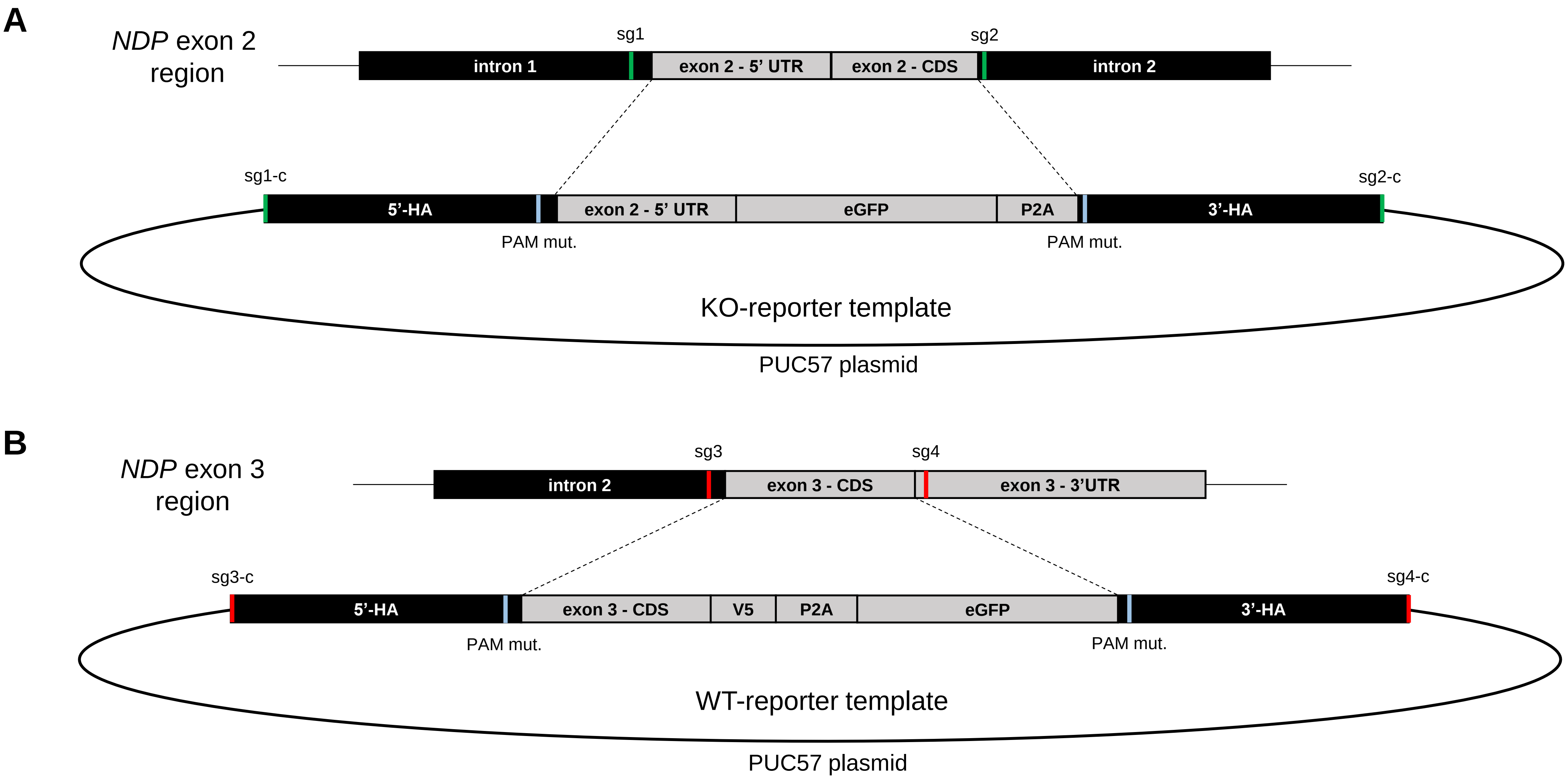


**Figure S01: CRISPR/Cas9 editing of human iPSCs**. (**A**) Dual sgRNA (sg1 and sg2) targeting regions flanking exon 2 were used to generate HDR-mediated *NDP*$^{KO\text{-}GFP}$ reporter lines, using a PUC57 vector harboring the KO-reporter template, consisting of an exon2 (5'UTR)-eGFP-P2A cassette, flanked by homology arms (HA), which included PAM site mutations and sgRNA-cleavage sites for plasmid linearization (sg1-c and sg2-c). NHEJ-mediated *NDP*$^{KO}$ (*NDP*$^{\Delta exon2}$), as well as *NDP*$^{WT}$ isogenic clones were generated in the process as well. (**B**) A similar approach was used to generate *NDP*$^{WT\text{-}GFP}$ reporter lines. Dual sgRNA (sg3 and sg4) flanking the CDS of exon 3 and a plasmid harboring the repair template, consisting of an exon3(CDS)-V5-P2A-eGFP cassette, flanked by homology arms, which included PAM site mutations and sgRNA cleavage sites for plasmid linearization (sg3-c and sg4-c) were used. sgRNA, small-guide RNA; HDR, homology directed repair; KO-GFP, knock-out reporter; KO, knockout; WT, wildtype; PAM, protoscpacer adjacent motif; UTR, untranslated region; CDS, coding sequence; HA, homology arm; eGFP, enhanced green fluorescent protein; P2A, porcine teschovirus-1 2A self-cleaving peptide; V5, V5 tag.

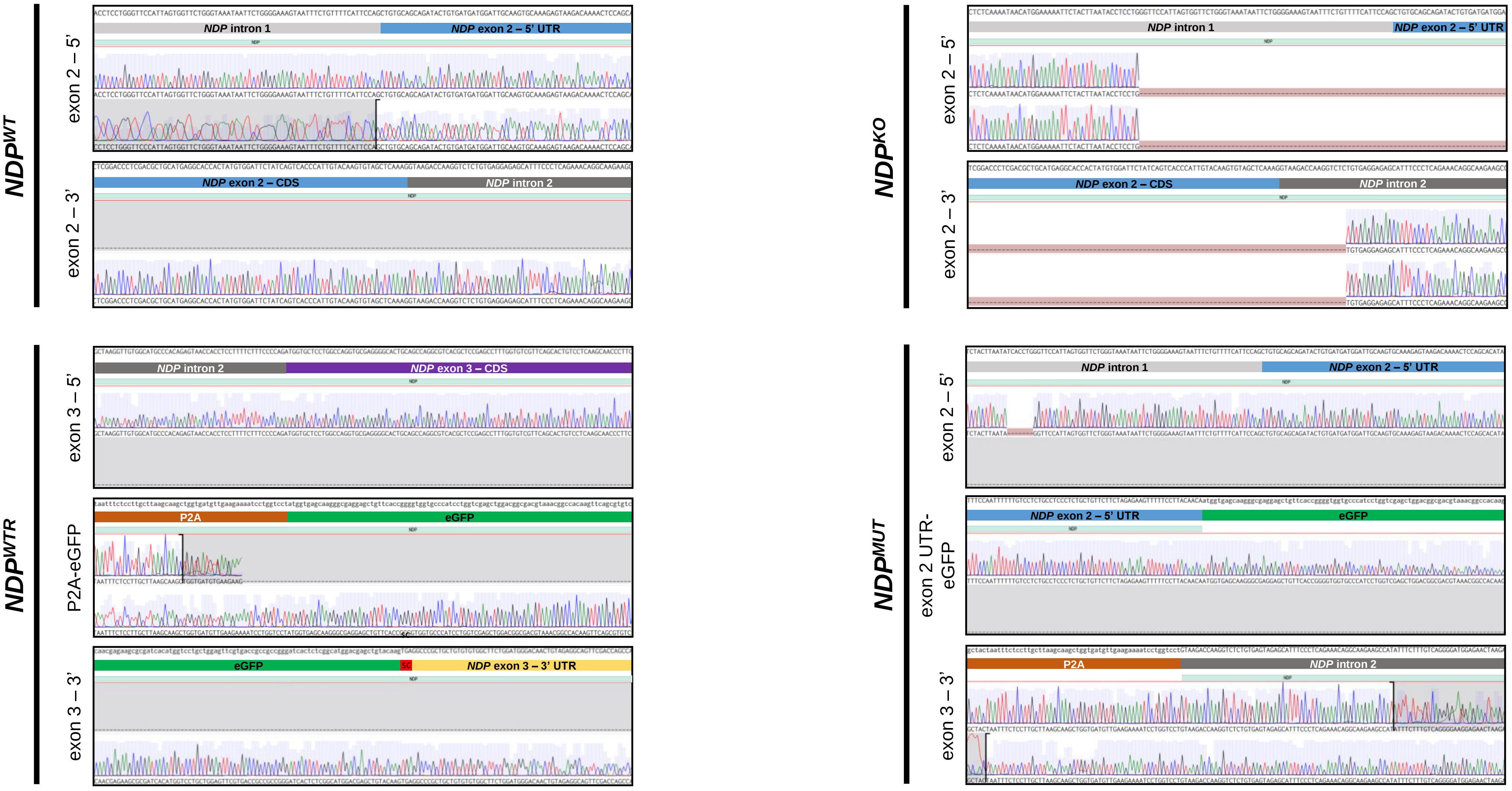


**Figure S02: Examples electropherograms of edited human iPSC lines**. Genomic DNA was extracted from selected clones and amplified using either a combination of NDP_e2_5'_fw and NDP_e2_3'_rev (*$NDP^{KO}$*, *$NDP^{KO\text{-}GFP}$*, *$NDP^{WT}$*) or NDP_e3_5'_fw and NDP_e3_3'_rev (*$NDP^{WT\text{-}GFP}$*)(Table S2A). Sanger sequencing was performed using NDP_e2_5'_fw and NDP_e2_3'_rev2 (*$NDP^{KO}$*, *$NDP^{KO\text{-}GFP}$*, *$NDP^{WT}$*) or NDP_e3_5'_fw and NDP_e3_3'_rev (*$NDP^{WT\text{-}GFP}$*) and in addition, for the reporters, eGFP internal primers (GFP_nterm_rev and GFP_int_rev) were used (Table S2A). Sequences from WT or KO clones were aligned to canonical NDP reference (GRCh38p14), while for the reporter lines (*$NDP^{WT\text{-}GFP}$* and *$NDP^{KO\text{-}GFP}$*), custom reporter reference were constructed. Visualization of alignments was performed on Benchling (https://www.benchling.com, accessed 01.08.2024). UTR, untranslated region; CDS, coding sequence, iPSC, induced pluripotent stem cells; KO, knockout, KO-GFP, knockout reporter; WT, wildtype; WT-GFP, wildtype reporter.

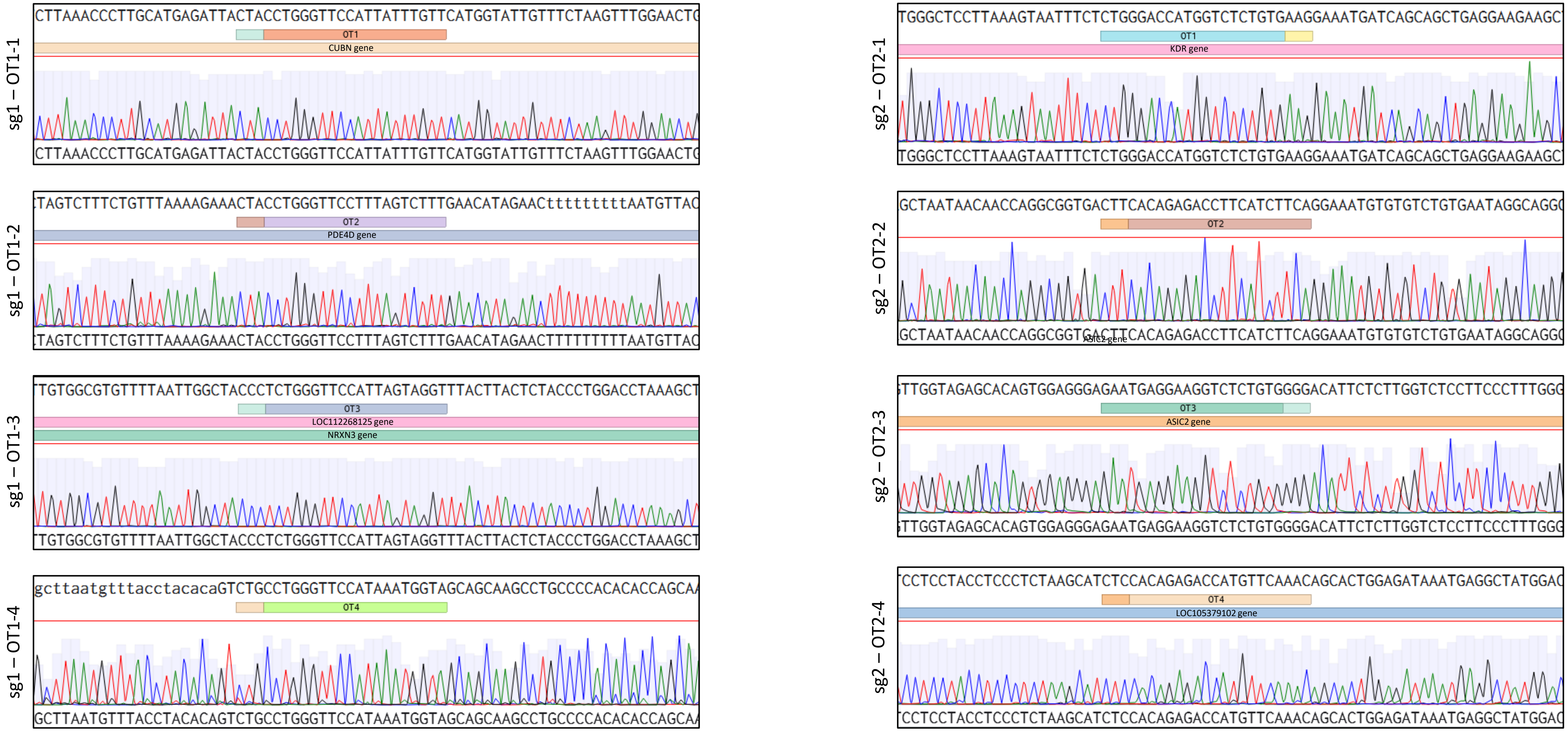


**Figure S03. Example electropherograms of top putative off-target binding sites in an *NDP*$^{KO}$ iPSC line.** The top four off-target sites of each sgRNA were selected for screening (Table S1B,C). Genomic DNA from edited clones was amplified using primer pairs OT1_1-4-fw/rev and OT2_1-4-fw/rev (Table 2B). Sanger sequencing was performed using a single primer of the pair. Sequences were aligned to GRCh38p14 reference where the predicted off-target cut site is marked by the top annotation under the reference sequence (OT1-4) together with its PAM-site (small adjacent annotation). Gene elements are visualized under the putative off-target site. Alignments were visualized using Benchling (https://www.benchling.com). iPSC, induced pluripotent stem cell; KO, knockout; PAM, protospacer adjacent motif; OT, off-target site.

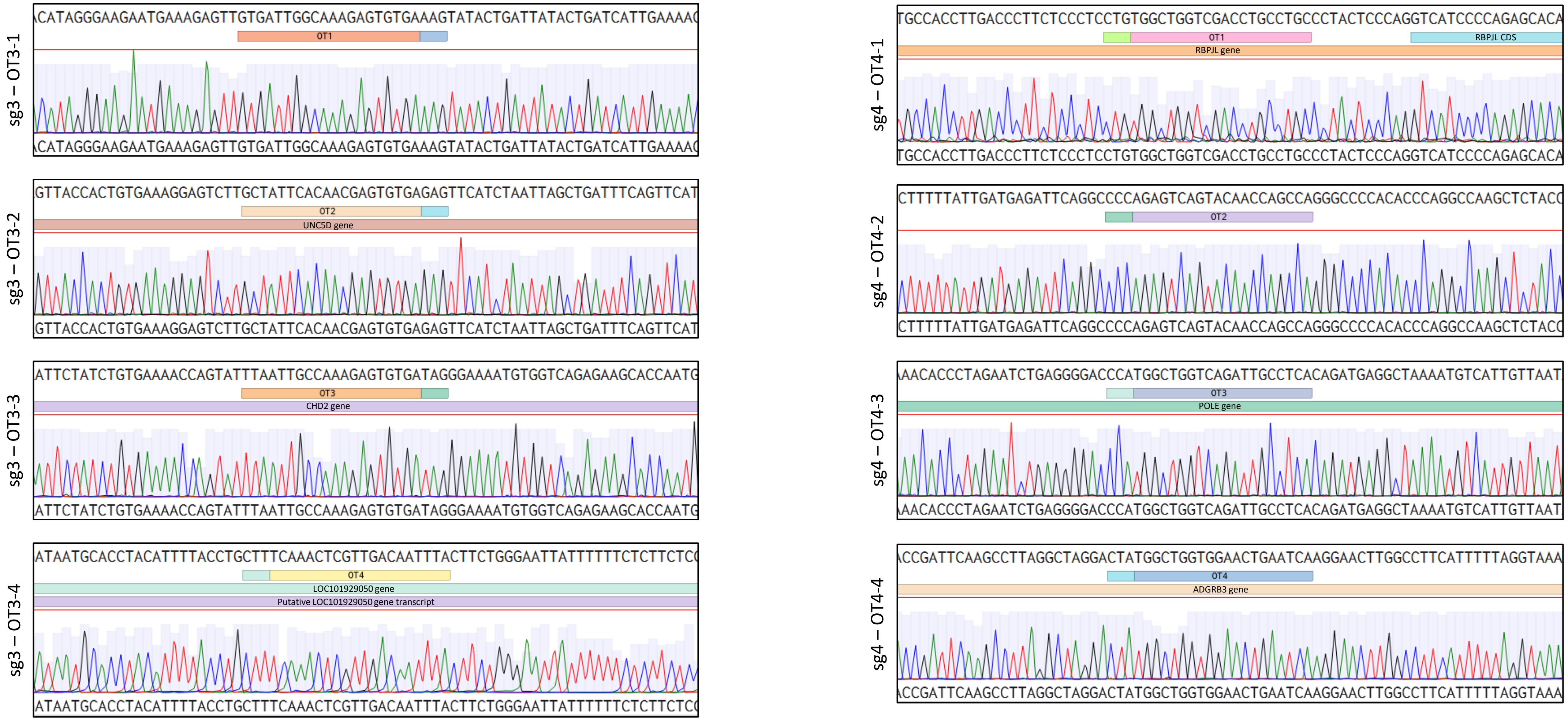


**Figure S04. Example electropherograms of top putative off-target binding sites in an *NDP*$^{WT\text{-}GFP}$ iPSC line.** The top four off-target sites of each sgRNA were selected for screening (Table S1C,D). Genomic DNA from edited clones was amplified using primer pairs OT1_1-4-fw/rev and OT2_1-4-fw/rev (Table 2B). Sanger sequencing was performed using a single primer of the pair. Sequences were aligned to GRCh38p14 reference where the predicted off-target cut site is marked by the top annotation under the reference sequence (OT1-4) together with its PAM-site (small adjacent annotation). Gene elements are visualized under the putative off-target site. Alignments were visualized using Benchling (https://www.benchling.com). iPSC, induced pluripotent stem cell; KO, knockout; PAM, protospacer adjacent motif; OT, off-target site.

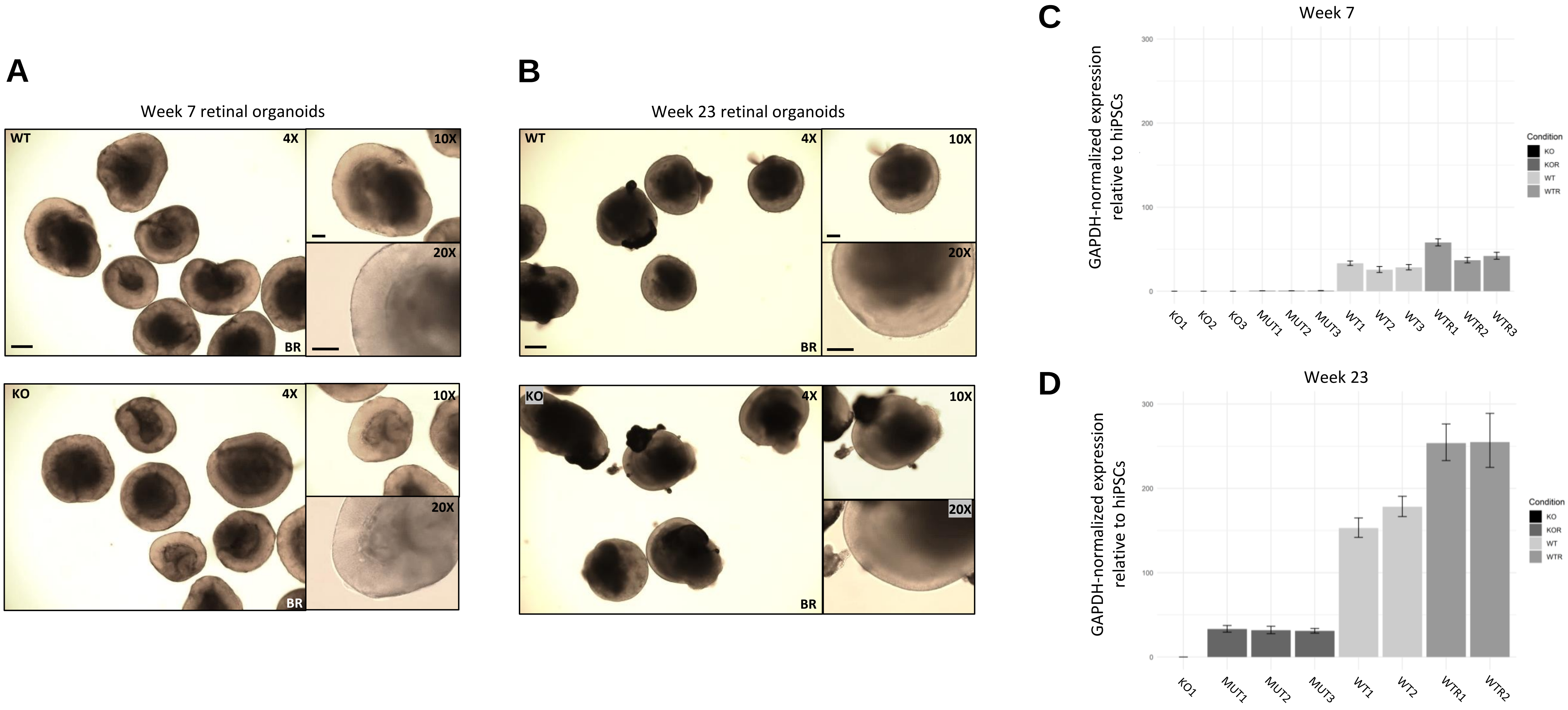


**Figure S05: Retinal organoids morphology**. (**A**) Brightfield (BR) images of human retinal organoids, portrayed at different magnifications (4X, 10X and 20X) displaying comparable morphology. Scale bars represent 200 µm, 100 µm and 100 µm respectively. (**B**) Brightfield (BR) images of human retinal organoids, portrayed at different magnifications (4X, 10X and 20X) displaying comparable morphology. Scale bars represent 200 µm, 100 µm and 100 µm respectively. (**C**) RT-qPCR performed with primers targeting sequences in the exon2 CDS and exon 3 CDS, respectively. Values represent cycle threshold (Ct) normalized to *GAPDH* levels and are displayed relative to the expression detected in WT iPSC lines. KO, knockout; MUT, mutant; WT, wildtype; WTR, wildtype reporter.

**A**

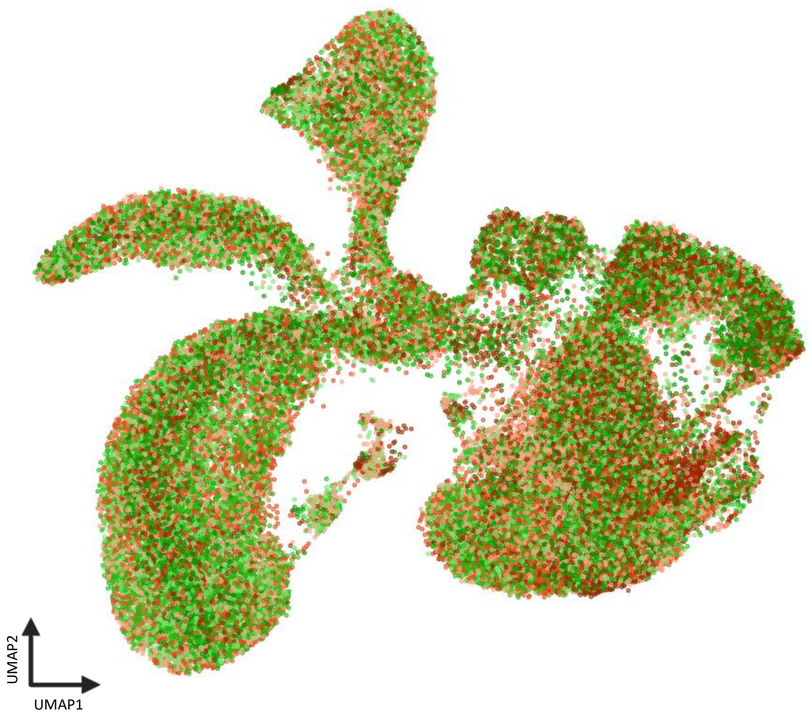


| Sample | Cells after quality filtering |
|---|---|
| • **W7_WT1** | 10661 |
| • **W7_WT2** | 12091 |
| • **W7_WT3** | 10558 |
| • **W7_KO1** | 10279 |
| • **W7_KO2** | 11070 |
| • **W7_KO1** | 11717 |

**B**

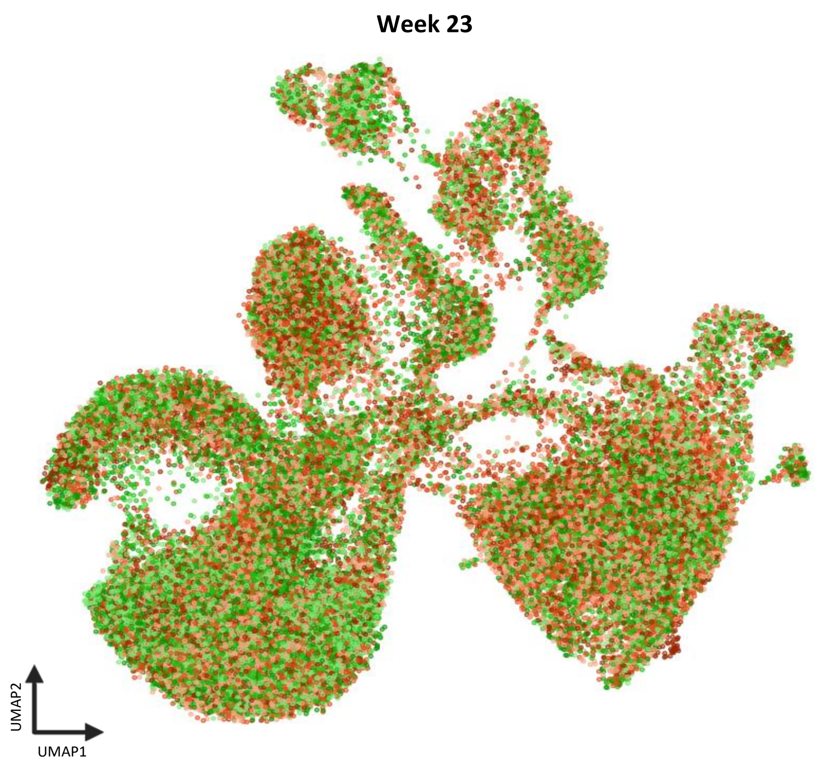


| Sample | Cells after quality filtering |
|---|---|
| • **W23_WT1** | 6365 |
| • **W23_WT2** | 7243 |
| • **W23_WT3** | 9236 |
| • **W23_KO1** | 7051 |
| • **W23_KO2** | 9724 |
| • **W23_KO3** | 10260 |

**Figure S06: UMAP representation of single cells recovered from scRNA-seq experiments**. (**A**) UMAP of integrated single cells derived from the sequencing of WT and KO week 7 human retinal organoids. Sequencing was performed using fixed scRNA-sequencing approach. *NDP*$^{WT}$-derived cells are represented in green colors whereas *NDP*$^{KO}$ cells are represented in red. The table reports the numbers of cells from each sample recovered after quality filters were applied. Processing and visualization of data was performed on Trailmaker (https://app.trailmaker.parsebiosciences.com, accessed 15.07.2024). (**B**) UMAP of integrated single cells derived from the sequencing of WT and KO week 23 human retinal organoids. Sequencing was performed using fixed scRNA-sequencing approach. *NDP*$^{WT}$-derived cells are represented in green colors whereas *NDP*$^{KO}$ cells are represented in red. The table reports the numbers of cells from each sample recovered after quality filters were applied. Processing and visualization of data was performed on Trailmaker (https://app.trailmaker.parsebiosciences.com, accessed 15.07.2024). UMAP, uniform manifold approximation and projection; W7, week 7; W23, week 23; scRNA-seq, single cell RNA sequencing; WT, wildtype; KO, knock-out.

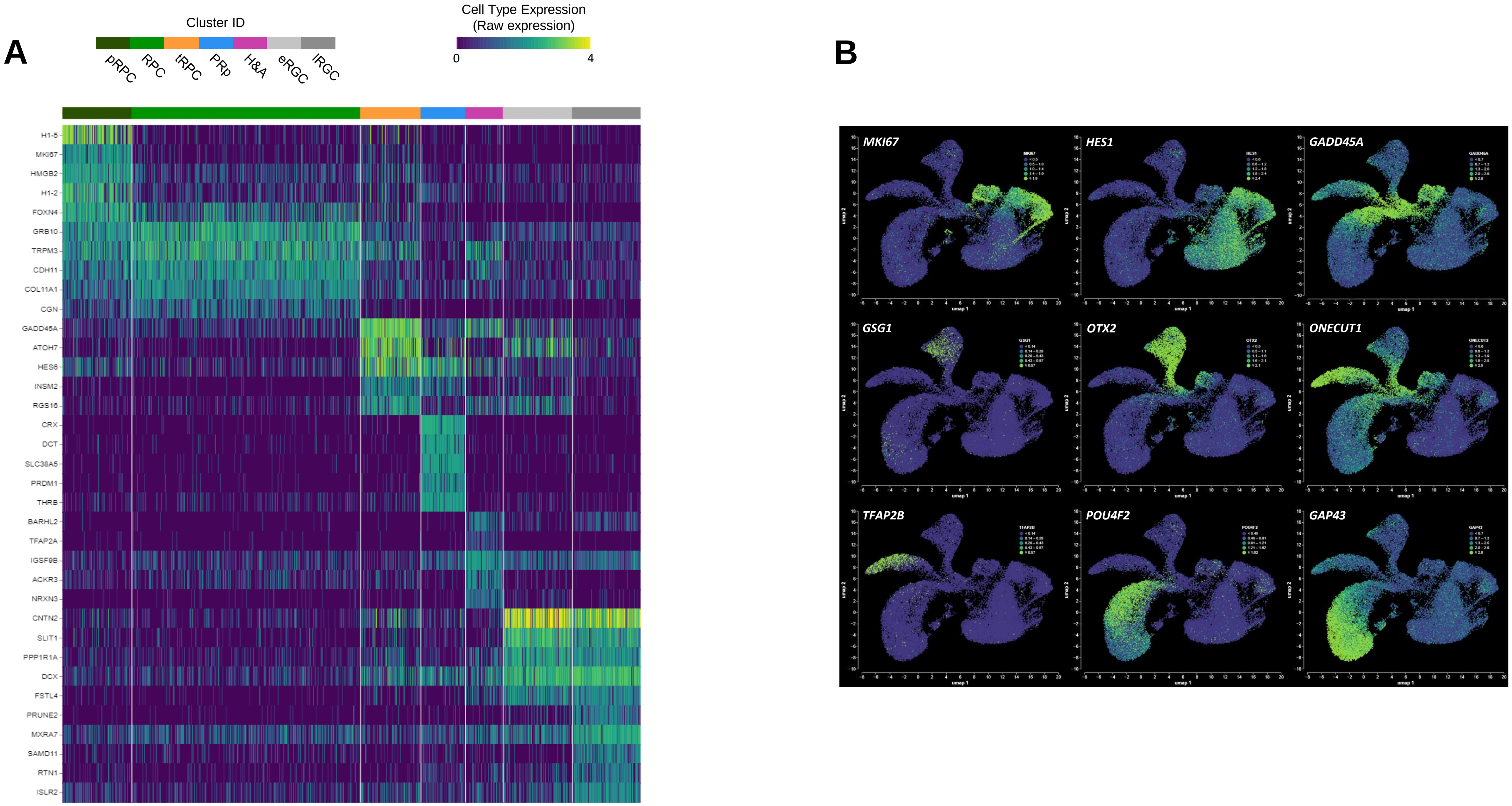


**Figure S07: Cell type specific markers in scRNA-seq week 7 data**. Hybridization-based PFA-fixed scRNA-seq was performed on W7 retinal organoids. The analysis of scRNA-seq data allowed recovering seven distinct cell clusters, which were identified respectively as: proliferating retinal progenitor cells (pRPC), retinal progenitor cells (RPC), transient retinal progenitor cells (tRPC), photoreceptor precursors (PRp), horizontal and amacrine cells (H&A), early retinal ganglion cells (eRGC) and late retinal ganglion cells (lRGC). (**A**) Heatmap representing the raw expression of the top-five markers expressed in in the identified clusters. (**B**) UMAPs displaying expression localization of selected marker genes in organoids samples. scRNA-seq, single-cell RNA sequencing; PFA, paraformaldehyde; UMAP, uniform manifold approximation and projection.

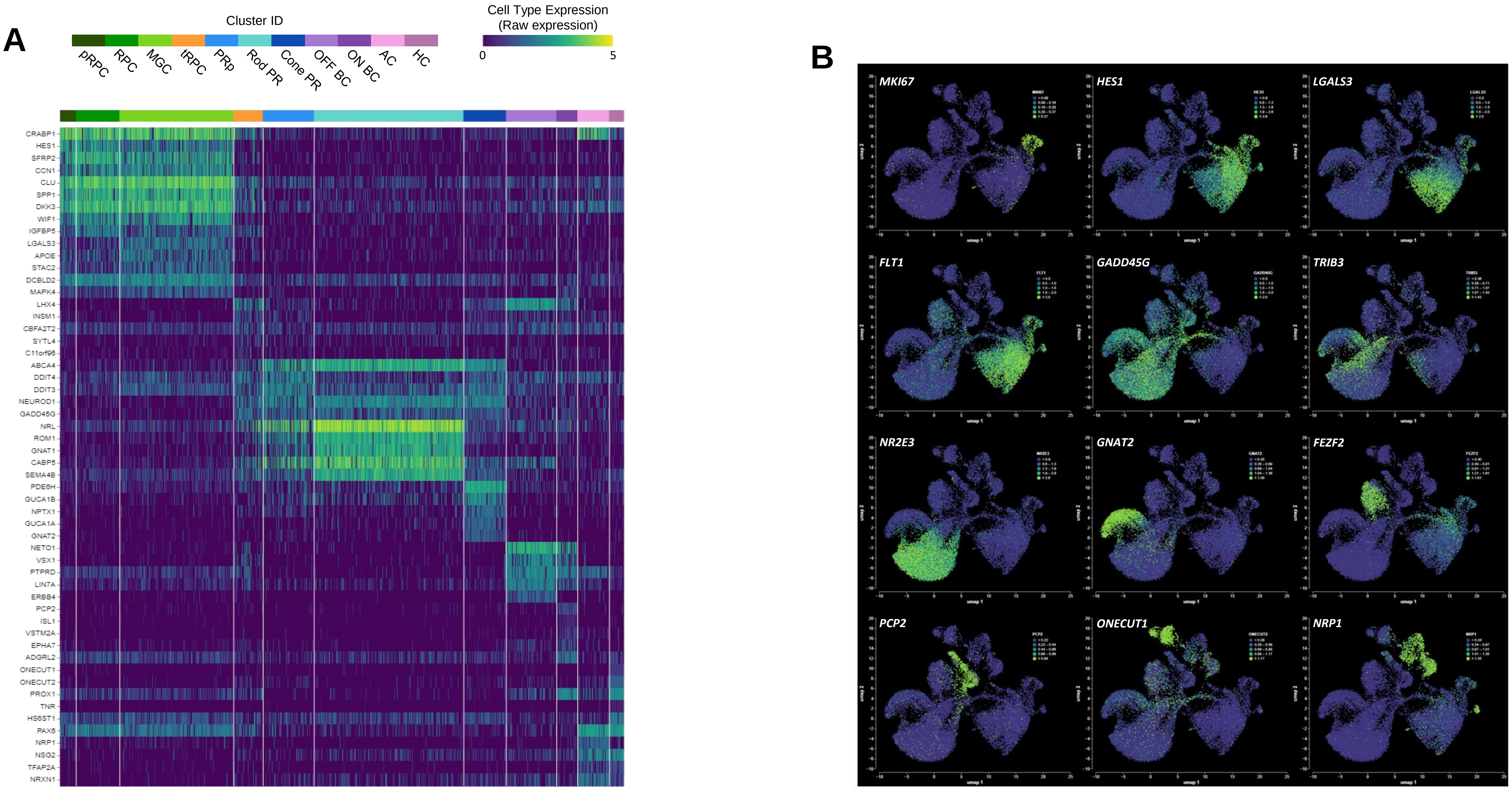


**Figure S08: Cell type specific markers in scRNA-seq week 23 data**. Hybridization-based PFA-fixed scRNA-seq was performed on W23 retinal organoids. The analysis of scRNA-seq data allowed recovering seven distinct cell clusters, which were identified respectively as: proliferating retinal progenitor cells (pRPC), retinal progenitor cells (RPC), Müller glia cells (MGC), transient retinal progenitor cells (tRPC), photoreceptor precursors (PRp), rod photoreceptors (Rod PR), cone photoreceptors (cone PR), OFF-bipolar cells (OFF BC), ON-bipolar cells (ON BC), amacrine cells (AC) and horizontal cells (HC). (**A**) Heatmap representing the raw expression of the top-five markers expressed in in the identified clusters. (**B**) UMAPs displaying expression localization of selected marker genes in organoids samples. scRNA-seq, single-cell RNA sequencing; PFA, paraformaldehyde; UMAP, uniform manifold approximation and projection.

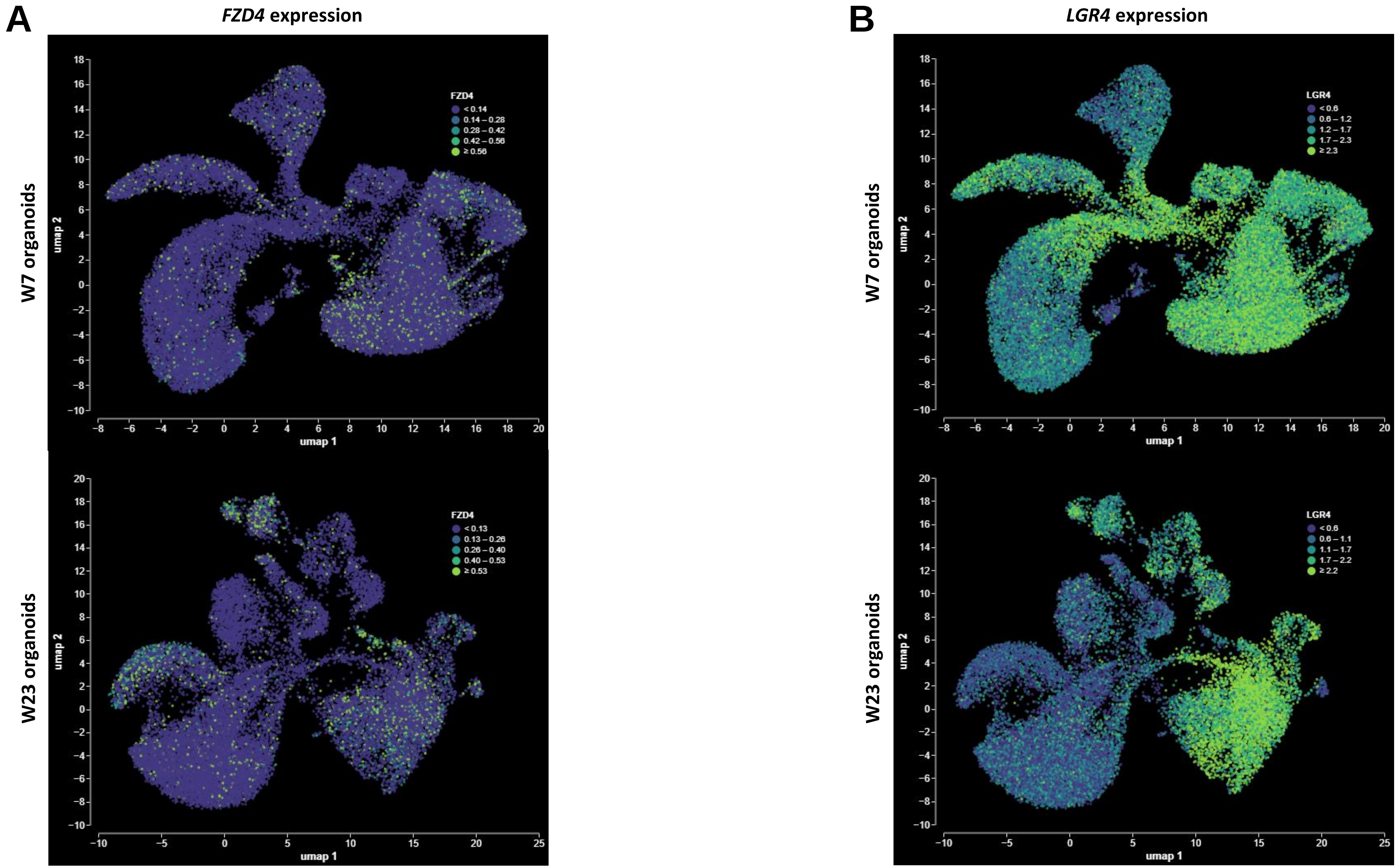


**Figure S09: Localization of expression of genes coding for Norrin-receptors in WT human retinal organoid, identified from scRNA-seq data**. (**A**) UMAPs displaying the localization of *FZD4* expression in retinal organoids at week 7 and week 23 of differentiation. (**B**) UMAPs displaying the localization of LGR4 expression in retinal organoids at week 7 and week 23 of differentiation. WT, wildtype; scRNA-seq, single-cell RNA sequencing; UMAP, uniform manifold approximation and projection.

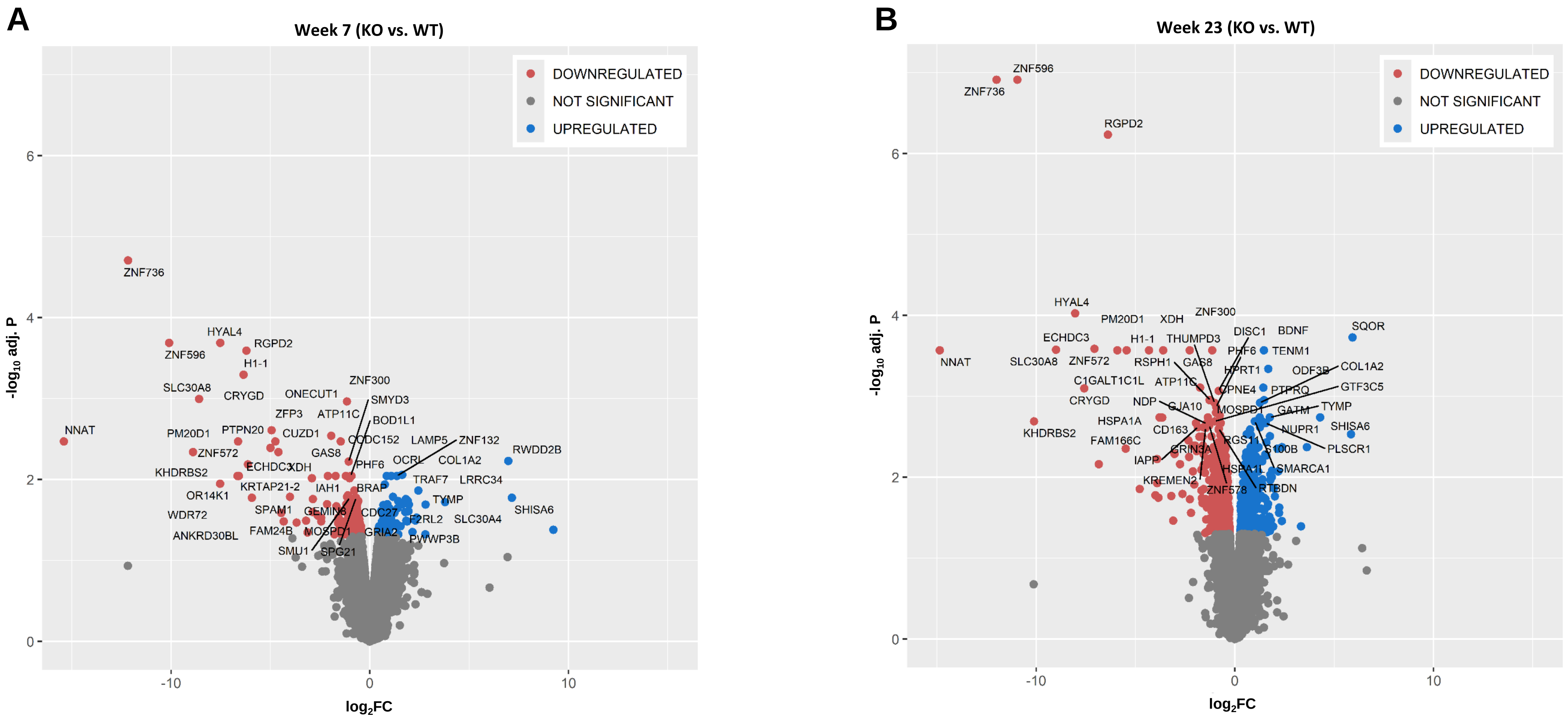


**Figure S010: DEGs detected in the scRNA-sequencing data in week 7 and week 23 retinal organoids**. (**A**) Volcano plot representing the 228 DEGs detected in the *NDP*$^{KO}$ retinal organoids analyzed at week 7 of differentiation (91 downregulated and 137 upregulated). For DEGs identification, a significance threshold of adjusted *p*-value <0.05 was applied. (**B**) Volcano plot representing the 933 DEGs detected in the *NDP*$^{KO}$ retinal organoids analyzed at week 23 of differentiation (581 downregulated and 352 upregulated). For DEGs identification, a significance threshold of adjusted *p*-value <0.05 was applied. DEGs, differentially expressed genes; KO, knock-out; WT, wildtype.

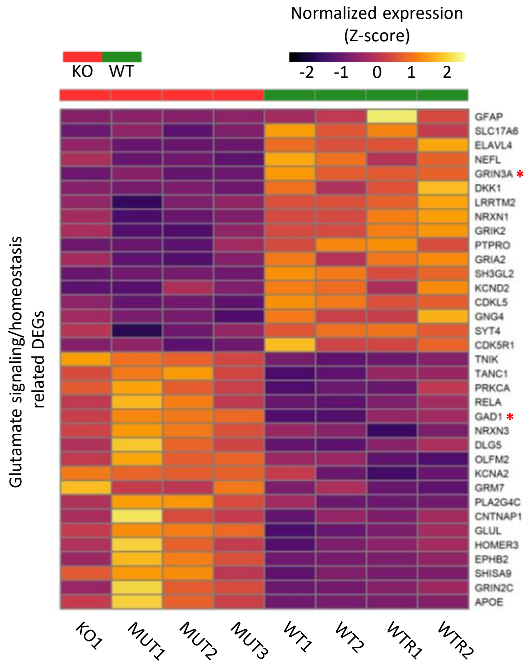


**Figure S11: Glutamate signaling related DEGs detected in week 23 KO/MUT retinal organoid**s. Heatmap representing the DEGs found in week 23 retinal organoids that are associated to the glutamate signaling system based on DAVID annotation (e.g. homeostastis, trafficking, synapse,..). *DEGs were detected also in the week 7 DEGs. DEGs, differentially expressed genes; KO, knock-out.

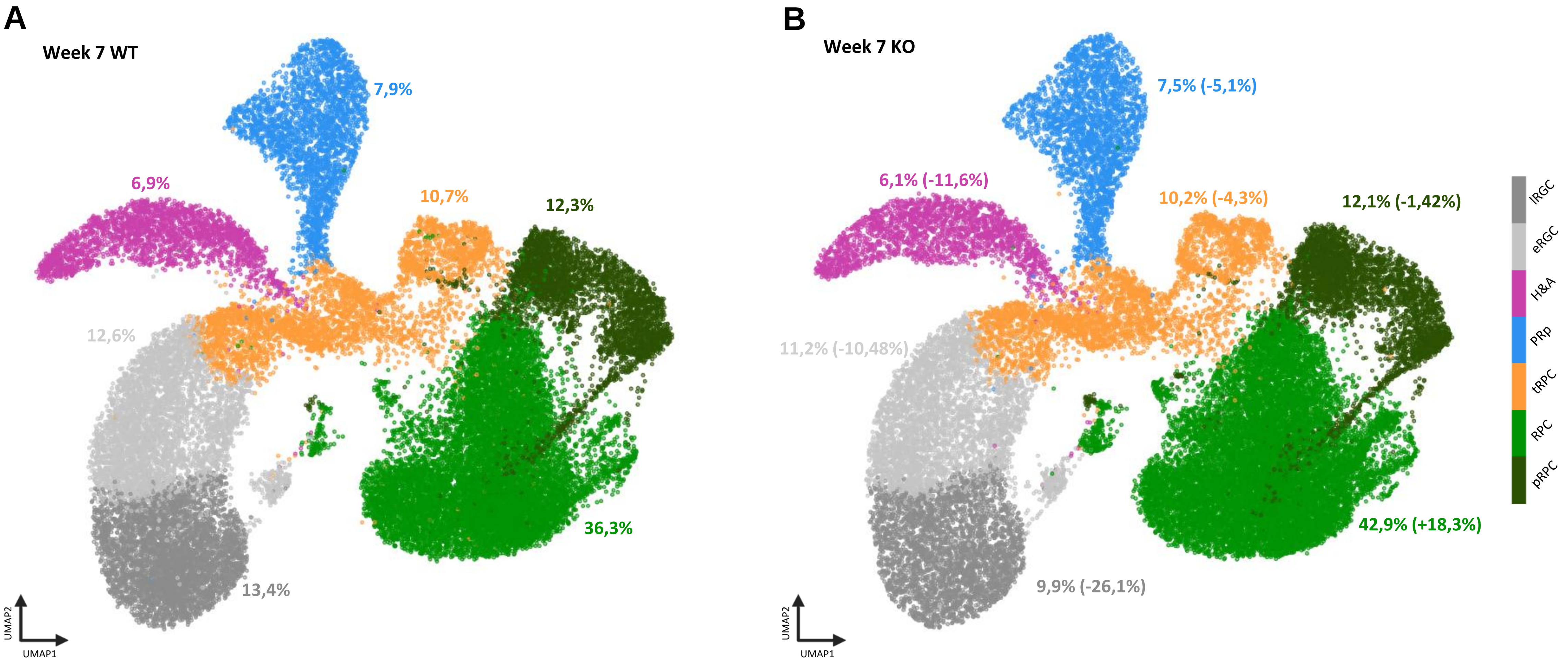


**Figure S12: Cell composition in week 7 *NDP*$^{WT}$ and *NDP*$^{KO}$ retinal organoids**. The analysis of week 7 scRNA-seq data allowed recovering seven distinct cell clusters, which were identified respectively as: proliferative retinal progenitor cells (pRPC), retinal progenitor cells (RPC), transient retinal progenitor cells (tRPC), photoreceptor precursors (PRp), horizontal and amacrine cells (H&A), early retinal ganglion cells (eRGC) and late retinal ganglion cells (lRGC). (**A**) UMAP of week 7 *NDP*$^{WT}$ retinal organoids reporting the cell composition. The percentages represent the proportion of the total cells ascribed to each cluster. (**B**) UMAP of week 7 *NDP*$^{KO}$ retinal organoids reporting the cell composition. The percentages represent the proportion of the total cells ascribed to each cluster. The values between brackets represent the change of the specific cluster proportion when compared to the WT proportion. WT, wildtype; KO, knock-out; UMAP, uniform manifold approximation and projection

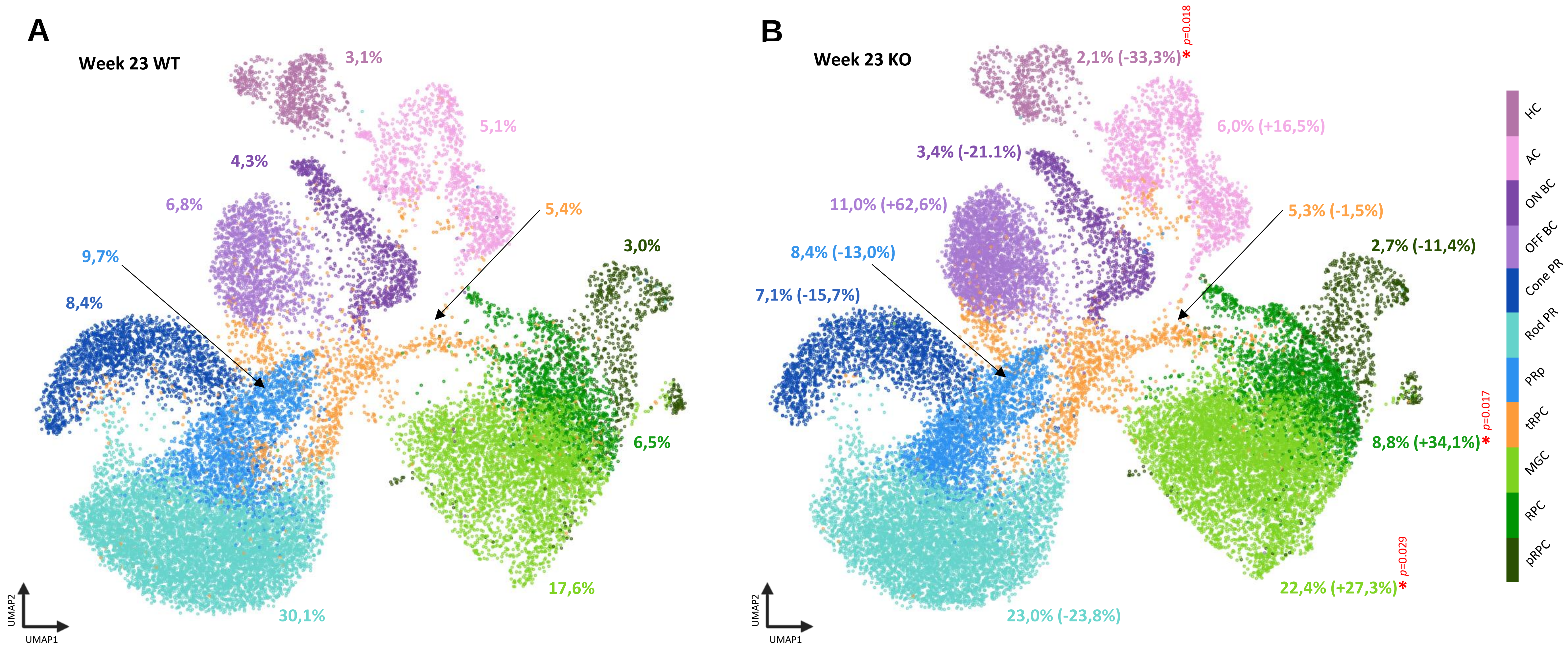


**Figure S13: Cell composition in week 23 *NDP*$^{WT}$ and *NDP*$^{KO}$ retinal organoids**. The analysis of week 23 scRNA-seq data allowed recovering seven distinct cell clusters, which were identified respectively as: proliferative retinal progenitor cells (pRPC), retinal progenitor cells (RPC), Müller glia cells (MGC), transient retinal progenitor cells (tRPC), photoreceptor precursors (PRp), rod photoreceptors (Rod PR), cone photoreceptors (Cone PR), OFF-bipolar cells (OFF BC), ON-bipolar cells (ON BC), amacrine cells (AC) and horizontal cells (HC) early retinal ganglion cells (eRGC) and late retinal ganglion cells (lRGC). (**A**) UMAP of week 23 *NDP*$^{WT}$ retinal organoids reporting the cell composition. The percentages represent the proportion of the total cells ascribed to each cluster. (**B**) UMAP of week 23 *NDP*$^{KO}$ retinal organoids reporting the cell composition. The percentages represent the proportion of the total cells ascribed to each cluster. The values between brackets represent the change of the specific cluster proportion when compared to the WT proportion. *Changes were found to be significant (*p*-value < 0.05). WT, wildtype; KO, knockout; UMAP, uniform manifold approximation and projection

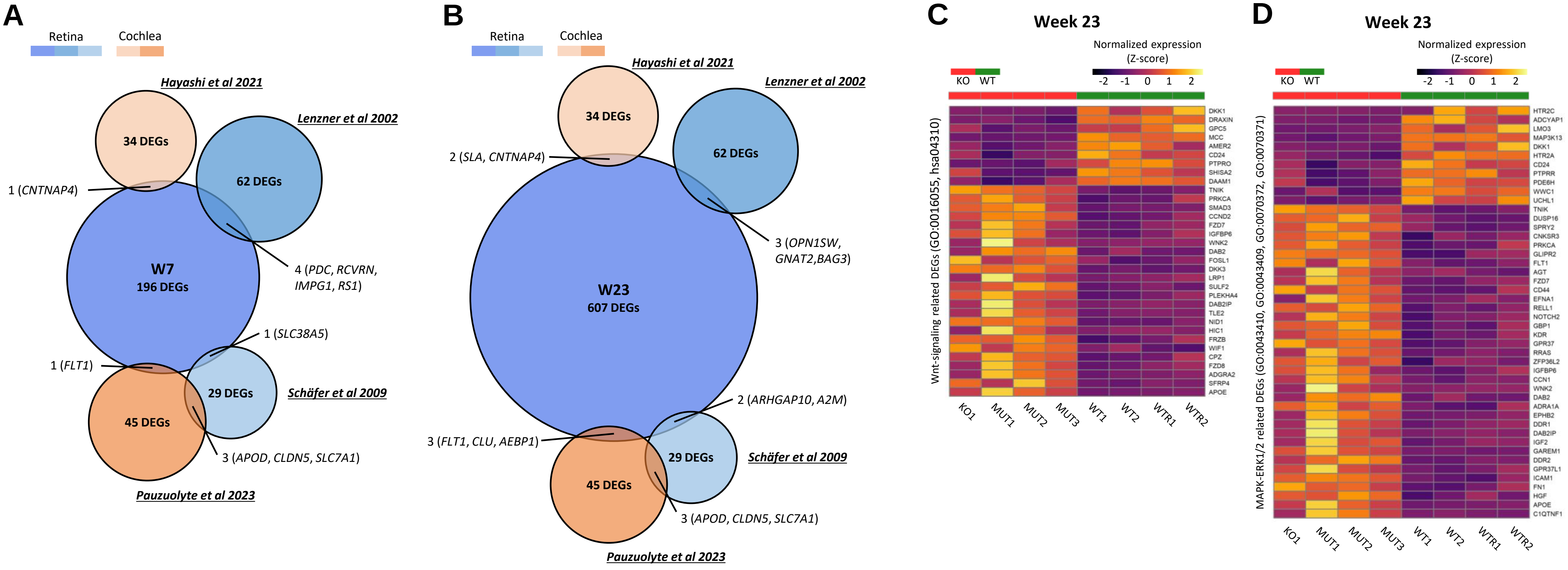


**Figure S14: Results of the comparison with previous studies**. Four publications reporting differentially expressed genes in $Ndp^{KO}$ mouse models were identified, two lists referred to DEGs in the retina (Lenzner et al. 2002 and Schäfer et al. 2009) and two to DEGs in the cochlea (Hayashi et al. 2021 and Pauzuolyte et al. 2023). Lists of human orthologs of the respective DEGs lists were compiled and compared to both, the week 7 and the week 23 DEGs detected in our study. Overlaps in affected pathways were also subject of analysis. (**A**) Venn diagram representing the overlapping DEGs found when comparing the lists of DEGs with the week 7 DEGs detected in the present study. (**B**) Venn diagram representing the overlapping DEGs found when comparing the lists of DEGs with the week 23 DEGs detected in the present study. (**C**) Heatmap representing the DEGs associated to Wnt-signaling found in our week 23 DEGs, that were detected by the ORA within the indicated terms. (**D**) Heatmap representing the DEGs associated to MAPK-ERK1/2-signaling cascade found in our week 23 DEGs, that were detected by the ORA within the indicated terms. DEGs, differentially expressed genes; KO, knockout; ORA, over-representation analysis.

**A**

| Name | Sequence | PAM | Score | Gene | Chromosome | Strand | Position | Mismatches |
|---|---|---|---|---|---|---|---|---|
| **sg1** | **AACCACTAATGGAACCCAGG** | **AGG** | **100** | | **chrX** | **+** | **43958915** | **0** |
| OT1-1 | AACAAATAATGGAACCCAGG | TAG | 3.303 | | chr10 | - | 17019724 | 2 |
| OT1-2 | AAAGACTAAAGGAACCCAGG | TAG | 2.512 | | chr5 | - | 59316341 | 3 |
| OT1-3 | ACCTACTAATGGAACCCAGA | GGG | 1.873 | | chr14 | - | 79242508 | 3 |
| OT1-4 | TACCATTTATGGAACCCAGG | CAG | 1.673 | | chr20 | - | 13693373 | 3 |
| OT1-5 | CATCACTAATGAAACCCAGG | AGG | 1.560 | | chr20 | - | 13173738 | 3 |
| OT1-6 | AATCAGTCATGGAACCCAGG | TGG | 1.542 | | chr8 | - | 78517147 | 3 |
| OT1-7 | AGCAACTGAGGGAACCCAGG | TGG | 1.433 | | chr5 | - | 176876983 | 4 |
| OT1-8 | CAGCCCTAACGGAACCCAGG | TGG | 1.425 | | chrX | + | 9443467 | 4 |
| OT1-9 | AGAGGCTAATGGAACCCAGG | CAG | 1.326 | | chr14 | + | 89308898 | 4 |
| OT1-10 | TATCACTCATTGAACCCAGG | GAG | 0.907 | | chr2 | + | 147082495 | 4 |

**B**

| Name | Sequence | PAM | Score | Gene | Chromosome | Strand | Position | Mismatches |
|---|---|---|---|---|---|---|---|---|
| **sg2** | **GTAAGACCAAGGTCTCTGTG** | **AGG** | **100** | | **chrX** | **-** | **43958455** | **0** |
| OT2-1 | CTGGGACCATGGTCTCTGTG | AAG | 1.413 | | chr4 | + | 55107159 | 4 |
| OT2-2 | TGAAGATGAAGGTCTCTGTG | AAG | 1.044 | | chr7 | - | 41153996 | 4 |
| OT2-3 | GAATGAGGAAGGTCTCTGTG | GGG | 1.001 | | chr17 | + | 33813951 | 4 |
| OT2-4 | GTTTGAACATGGTCTCTGTG | GAG | 0.932 | | chr5 | - | 101573093 | 4 |
| OT2-5 | GTTCCAGCAAGGTCTCTGTG | AAG | 0.926 | | chr14 | - | 101223577 | 4 |
| OT2-6 | GGTAGACTCAGGTCTCTGTG | GAG | 0.921 | | chr7 | + | 30856145 | 4 |
| OT2-7 | GAGAGACTCAGGTCTCTGTG | CAG | 0.921 | | chr10 | + | 79219267 | 4 |
| OT2-8 | GACAGACAAACGTCTCTGTG | GGG | 0.882 | | chr20 | + | 52728548 | 4 |
| OT2-9 | GAAACACAAATGTCTCTGTG | AGG | 0.879 | | chr22 | + | 36693605 | 4 |
| OT2-10 | GAAGGACCAGAGTCTCTGTG | AAG | 0.831 | | chr2 | + | 114796889 | 4 |

**Table S1A,B: CRISPR/Cas9 guide design for gene editing of the *NDP* locus**. The combination of sgRNA sg1 (**Table S1A**) and sg2 (**Table S1B**) was used to edit the region including exon 2 (see Figure S01A). The table reports the on-target sequence (marked in grey) and the top-10 ranking putative off-target binding sites. Target score ranges from 0-100, representing the likelihood of cutting events. sgRNA, small-guide RNA; OT, off-target site.

**C**

| Name | Sequence | PAM | Score | Gene | Chromosome | Strand | Position | Mismatches |
|---|---|---|---|---|---|---|---|---|
| **sg3** | **GGAATTGGCAACGAGTGTGA** | **GGG** | **100** | | **chrX** | **-** | **43950110** | **0** |
| OT3-1 | GTGATTGGCAAAGAGTGTGA | AAG | 1.499 | | chr5 | + | 90124349 | 3 |
| OT3-2 | GCTATTCACAACGAGTGTGA | GAG | 0.995 | | chr8 | + | 35746649 | 4 |
| OT3-3 | TTAATTGCCAAAGAGTGTGA | TAG | 0.847 | ENSG00000173575 | chr15 | + | 92947196 | 4 |
| OT3-4 | TAAATTGTCAACGAGTTTGA | AAG | 0.775 | | chr7 | - | 63388330 | 4 |
| OT3-5 | TAAATTGTCAACGAGTTTGA | AAG | 0.775 | | chr7 | + | 63359786 | 4 |
| OT3-6 | AGAAATTGCAAGGAGTGTGA | AAG | 0.557 | | chr6 | - | 72745397 | 4 |
| OT3-7 | TCAATTGGAAAGGAGTGTGA | GGG | 0.522 | | chr2 | + | 41898574 | 4 |
| OT3-8 | GGAATTACTGACGAGTGTGA | AAG | 0.517 | | chr4 | - | 22412210 | 4 |
| OT3-9 | TGGATTGGGAATGAGTGTGA | GGG | 0.510 | | chr2 | + | 46964381 | 4 |
| OT3-10 | AGAAATGGGAAAGAGTGTGA | CGG | 0.508 | | chr1 | + | 171660075 | 4 |

**D**

| Name | Sequence | PAM | Score | Gene | Chromosome | Strand | Position | Mismatches |
|---|---|---|---|---|---|---|---|---|
| **sg4** | **AGAGGCAGTTCGACCAGCCA** | **GGG** | **100** | **ENSG00000124479** | **chrX** | **-1** | **43949743** | **0** |
| OT4-1 | GCAGGCAGGTCGACCAGCCA | CAG | 1.751 | | chr20 | -1 | 45309539 | 3 |
| OT4-2 | AGAGTCAGTACAACCAGCCA | GGG | 1.253 | | chr1 | 1 | 177814033 | 3 |
| OT4-3 | GCATGCTGTTCGACCAGCCA | GGG | 0.993 | | chrY | -1 | 18336540 | 4 |
| OT4-4 | GCATGCTGTTCGACCAGCCA | GGG | 0.993 | | chrY | 1 | 17569472 | 4 |
| OT4-5 | TGAGGCAATCTGACCAGCCA | TGG | 0.824 | ENSG00000177084 | chr12 | -1 | 132632871 | 4 |
| OT4-6 | TGATTCAGTTCCACCAGCCA | TAG | 0.808 | | chr6 | -1 | 69295342 | 4 |
| OT4-7 | AGGGGCACTGGGACCAGCCA | CAG | 0.771 | | chr11 | 1 | 69617665 | 4 |
| OT4-8 | AGTGGCATTAAGACCAGCCA | TGG | 0.771 | | chr12 | 1 | 2734469 | 4 |
| OT4-9 | AGGTCCAGTTCTACCAGCCA | CAG | 0.755 | | chr15 | 1 | 67147683 | 4 |
| OT4-10 | GTAGGCTGTTTGACCAGCCA | CAG | 0.628 | | chr8 | 1 | 101375263 | 4 |

**Table S1B,C: CRISPR/Cas9 guide design for gene editing of the *NDP* locus**. The combination of sgRNAs sg3 (**Table S1C**) and sg4 (**Table S1D**) was used to edit the region including the exon 3-CDS (see Figure S01B). The table reports the on-target sequence (marked in grey) and the top-10 ranking putative off-target binding sites. Target score ranges from 0-100, representing the likelihood of cutting events. CDS, coding sequence; OT, off-target site; sgRNA, small-guide RNA.

**A**

| Name | Sequence (5' - 3') |
|---|---|
| NDP_e2_5'_Fw | CCCCGACCAACCAAGGACAAAGATG |
| NDP_e2_3'_Rev | AGCCTCATTCTCCCACAAGCCCT |
| NDP_e2_3'_Rev2 | CAGCCACATCACTTAAGTTTGGGCT |
| NDP_e3_5'_Fw | AGCCTCCGTTATTCCAGCCCAATC |
| NDP_e3_3'_Rev | AGCATTGAGAGCCAAGGGGGAA |
| GFP_nterm__Rev | TCCTCGCCCTTGCTCACCAT |
| GFP_int_Rev | AGTTGTACTCCAGCTTGTGCCCC |

**B**

| | Name | Sequence (5' - 3') |
|---|---|---|
| sg1 | OT1-1-Fw | GCACCTGAGATACTCCCACCCC |
| | OT1-1-Rev | TGTACTCAACCCACCTCAGCAGAT |
| | OT1-2-Fw | AAAACTGAGGGCCGTAGACCCT |
| | OT1-2-Rev | CCTCCGTCCACAACTGGAACATTG |
| | OT1-3-Fw | GCAAGCCAGATCTGAGGCCAAG |
| | OT1-3-Rev | CCTCAGGTTGTCTGCCTCACACTT |
| | OT1-4-Fw | GGACTTGGAAGCTCCCTTCAGC |
| | OT1-4-Rev | ATAGCTAAAGGGTTGTGCGGGAAG |
| sg2 | OT2-1-Fw | CAGGATACAGGAGGGAAGCAGCT |
| | OT2-1-Rev | TTTCCCATCTTCAGTCCCACCCC |
| | OT2-2-Fw | GGGACCAGAGTTGGAACTGGGT |
| | OT2-2-Rev | GGTTGAAATCCTCATTTGCCTCTC |
| | OT2-3-Fw | CCACCAGGCTGGAAGGAACAAG |
| | OT2-3-Rev | TTCTCCATGCAAGTCCCCACTCT |
| | OT2-4-Fw | ATTTCTGGTGCTGTTGAGAGAGACA |
| | OT2-4-Rev | GCCTCTGACCATTATTTTGTCCTCT |
| sg3 | OT3-1-Fw | TGCACCACACTTCACAGAGGAAC |
| | OT3-1-Rev | CACCACTCTACTGCCCTGTCCC |
| | OT3-2-Fw | AGAGCAAGAGCCTGTGAGTGAGA |
| | OT3-2-Rev | AGTGACCGATGACAAGCTCTCCT |
| | OT3-3-Fw | ACTGATGTGCTTGGGTCCTCAGTA |
| | OT3-3-Rev | AAAACCCAGCCATCCATCCACTC |
| | OT3-4-Fw | GGACAGGCAGACAGGGAGATG |
| | OT3-4-Rev | ATGTCTCGCAACTGTCCCTGAAAA |
| sg4 | OT4-1-Fw | GGAAAGCCCACACAGCAACAGG |
| | OT4-1-Rev | CAAAGTGGGGGCTAATGGTGGC |
| | OT4-2-Fw | CCTGGGGTTCTCTCTGGCTTGT |
| | OT4-2-Rev | TGGGGGGAGAATGAGTTGGGAAG |
| | OT4-3-Fw | CACACAGAGGAGTTAGGTCACTGG |
| | OT4-3-Rev | CTGCCTTTGCTTGTGTTGGGAG |
| | OT4-4-Fw | CAAGCCTGCTCTGACTGACCT |
| | OT4-4-Rev | GGGGTGGGGGCAGTTAGAAAAT |

**C**

| Name | Sequence (5' - 3') |
|---|---|
| NDP_qPCR_Fw | GCTCATTCATAATGGACTCGGACCC |
| NDP_qPCR_Rev | CTTGAGGACAGTGCTGAACGACAC |

**Table S2: List of primers used in the study**. Sequences are reported from 5' to 3'. (**A**) Primers used for genotyping of CRISPR/Cas9 edited hiPSC lines. (**B**) Primers used for the sequencing of the top-four ranking off-target binding sites of each sgRNA used for the CRISPR/Cas9 editing of the NDP locus. The respective guides are indicated on the side of the table. (**C**) Primer pair used for RT-qPCR. hiPSC, human induces pluripotent stem cells; OT, off-target site; RT-qPCR, reverse transcriptase quantitative polymerase chain reaction; sgRNA, small-guide RNA.
